# Warming makes 20 boreal species grow earlier but not always taller

**DOI:** 10.64898/2025.11.30.691420

**Authors:** Yiluan Song, Kai Zhu, Yang Chen, Artur Stefanski, Raimundo Bermudez, Rebecca A. Montgomery, Peter B. Reich

## Abstract

Climate warming is altering the timing of seasonal growth in cold-temperate and boreal ecosystems, with potential consequences for forests on multiple continents. It has been hypothesized that in a warming climate, earlier growth in spring might lead to a longer growing season and thus higher annual total growth. However, whether the period of active growth is longer in a warming climate is unclear, as is whether changes in growth rate might reinforce or offset the effect of changing growing season length in modulating annual total growth. Understanding responses in growth to warming is crucial to projecting forest dynamics under future climate change. To address this, we experimentally warmed (by +1.62 and 3.26 °C on average) open-air southern boreal forest plots for 14 years (2010 to 2023) and measured juvenile height growth from spring until fall for 20 tree and shrub species, with a total of 11,565 individual-year growth trajectories. Across all species and in various canopy (open, closed) and rainfall (ambient, reduced) contexts, experimental warming consistently advanced the timing of height growth (4.32 days per 3 °C warming). However, species differed in their annual total growth responses. Evergreen gymnosperms such as spruce, fir, and pine tended to exhibit suppressed annual total growth under warming (−15.8% per 3 °C warming), driven by a shorter duration of active growth, despite an earlier start, with little change in relative growth rate. In contrast, deciduous angiosperms and non-native shrubs often show enhanced annual total growth under warming (19.3 and 37.4% per 3 °C warming), driven by faster rate, longer duration, or a combination of both. More broadly, species with a more conservative lifestyle responded less positively in the duration and rate of growth, thus placing them at a disadvantage under warming. These contrasting responses challenge assumptions that an earlier start of the growing season universally enhances growth and highlight how warming may shift both the timing and pace of growth and thus alter community composition within mixed-species forests.

## Introduction

Plant growth, encompassing the increase in biomass, height, and/or diameter over time (Hilty et al., 2021), drives forest population dynamics (McDowell et al., 2020), influences competitive outcomes (Howard & Goldberg, 2001), and interacts with carbon sequestration (Piao, Wang, et al., 2019; Zhang et al., 2019). Plant growth plays a central role in shaping forest structure, composition, and function (Ford et al., 2017). Except in the aseasonal wet tropics, on an intra-annual scale, growth is inherently seasonal and tied to plant phenology, the timing of recurring developmental events such as budburst, shoot elongation, and growth cessation. Both plant growth and phenology are tightly regulated by temperature (Piao, Liu, et al., 2019; Saxe et al., 2001). Under climate warming in seasonally cold biomes, it is critical to quantitatively understand the coupling of plant growth and phenology, in order to anticipate the future of global forests. Whether or not the expected earlier growth onset under warming leads to a greater total annual growth has drastically different implications: forests may become taller, denser, and more carbon-rich, or they may experience restructured seasonal dynamics and community composition without an increase in carbon sink. Yet, from first principles, two apparently contradictory outcomes emerge.

Warming in seasonally cold biomes might enhance plant growth through two pathways, via plant phenology and physiology. Warming might increase productivity indirectly through an earlier start of the growing season, which, if coupled with an unchanged or later end of the growing season, allows for a longer photosynthetically active period (Gunderson et al., 2012). Additionally, higher temperatures can also directly increase the rates of photosynthesis (Sendall et al., 2015; Xia et al., 2015; S. Zhou et al., 2016), further increasing productivity. The stimulated productivity under warming, jointly driven by changing phenology and physiology, could explain the phenomenon of “global greening,” especially in temperate and boreal systems, where productivity is often temperature-limited (Piao et al., 2007; Piao, Wang, et al., 2019; Z. Zhu et al., 2016). Such warming-induced increases in productivity could lead to enhanced plant growth (Piao et al., 2007).

In contrast, warming might be expected to suppress plant growth through several phenological and physiological constraints. First, warming almost always increases evaporative demand and induces water stress (Dai et al., 2018). This water stress would be compounded when reduced total precipitation, more temporally clustered precipitation patterns, and/or lower soil moisture occur in association with warmer temperatures (Buermann et al., 2018; Reich et al., 2018). Under such conditions, plants might exhibit earlier cessation of growth (Ruehr & Nadal-Sala, 2025), downregulation of photosynthesis (Reich et al., 2018; Rennenberg et al., 2006; L. Zhou et al., 2015), or both (Reich et al., 2022). Second, enhanced aboveground growth is not guaranteed despite increased gross primary production (GPP), as plants might shift resource allocation toward belowground growth and/or stress tolerance mechanisms (Wiley & Helliker, 2012; L. Zhou et al., 2022). Finally, warming may disrupt winter chilling requirements that are essential for dormancy release in many temperate and boreal species (Baumgarten et al., 2021; Flynn & Wolkovich, 2018), potentially delaying the onset of growth despite warmer spring temperatures.

There is evidence both supporting enhanced and suppressed total growth in years with warmer springs and earlier growth onset. Satellite remote sensing and flux tower data suggest that the earlier start of annual growth leads to extended growing seasons (Piao et al., 2007) and higher GPP (Keenan et al., 2014). Observational studies at the individual or species level showed mixed growth responses. Warmer conditions over space and time have been associated with increased height growth, with differences among species (Bontemps et al., 2012; Messaoud & Chen, 2011). Dow et al. (2022) found no significant increase in stem diameter growth in warmer springs, despite earlier growth. Girardin et al. (2016) found regional- and species-related trends in stem biomass growth, but no strong overall trend when averaged. An analysis of wood microcores further suggested that trees with a longer growing season produced more cells but not more biomass in the wood (Silvestro et al., 2023). Limited experimental evidence has shown that stem biomass growth responses vary with species’ geographic range, with species near their warm range limits tending to respond negatively to warming (Reich et al., 2015).

These two contradictory expectations and corresponding observational evidence highlight a key knowledge gap: it remains unclear whether and how we can generalize regarding how warming influences growth under these opposing mechanisms. As a result, it is unclear whether we can accurately predict growth under future climate warming. Our understanding remains fragmented for several reasons. First, as observational studies leverage interannual or spatial climatic differences, observed discrepancies in growth responses might or might not reflect inconsistent responses to warming, as they might be results of different summer conditions that follow warm springs. Global change experiments that directly couple warmer springs with warmer summers are needed to accurately represent future climate change, generalize the effect of warming, and identify mechanisms for differential responses (but are rare). Second, despite the anticipated relationship between growing season and total growth, existing studies usually focus on either phenology or growth responses, rarely linking changes in the timing of growth with the magnitude of growth, which could be achieved with seasonal growth trajectories (Dow et al., 2022). To resolve these gaps, we need to examine the growth trajectories of individual plants in manipulated experiments.

Furthermore, species-specific growth responses to warming have often been treated as inconsistencies, but rarely as predictable differences (but see Reich et al., 2015, 2022). Might species’ differences in lifestyle influence how they respond to warming? Being able to predict growth responses under warming will help anticipate potential shifts in competitive dynamics and community composition. It will also help bridge the gap between results from the whole ecosystem or landscape level and the individual plant level.

In this study, we answer three questions. First, does warming lead to both earlier growth and enhanced annual total growth? We hypothesize that warming will lead to earlier growth, but warming will not consistently lead to enhanced annual total growth because of considerable degrees of freedom involving growth duration and growth rate responses to warming (Fig. 1A). Second, linking growth to phenology, are changes in annual total growth related to changes in duration or rate of growth? We hypothesize that enhanced annual total growth, when it occurs, will be driven by both longer duration and faster rate; and when it does not, will be driven by either shorter duration or slower rate. Third, can species’ growth and phenological responses be predicted by their lifestyles, reflected through their distributions and functional traits? We hypothesize that more acquisitive species respond more positively in total magnitude, duration, and rate of growth under warming compared to conservative species.

To test these hypotheses, we conducted a long-term open-air global change field experiment (Fig. S1A) that offers a unique opportunity to synthesize plant responses to simultaneous aboveground and belowground warming across 14 years from 2010 to 2023 (Fig. S2,S3, Table S2). The experiment is rare in implementing controlled warming both aboveground and belowground without chambers (Rich et al., 2015); additionally, contrasting warming treatments were imposed across different sites, canopy conditions (open and understory), and rainfall regimes (ambient and 40% less summer rainfall from June 1 to September 30) (Stefanski et al., 2020), providing a wide set of contexts to examine the generality of responses. Crucially, the experiment provides unique measurements on both the timing and magnitudes of terminal shoot elongation, a key component of early-season vegetative growth, tracked at a high temporal resolution (Fig. S1B,S4). This unique dataset, with 131,510 measurements from 11,565 height growth trajectories, enables us to integrate phenology and growth responses within a single framework. The inclusion of 20 co-occurring plant species with diverse lifestyles (Table S1) further allows us to assess interspecific variation in phenological and growth responses under shared environmental conditions.

**Figure 1.**
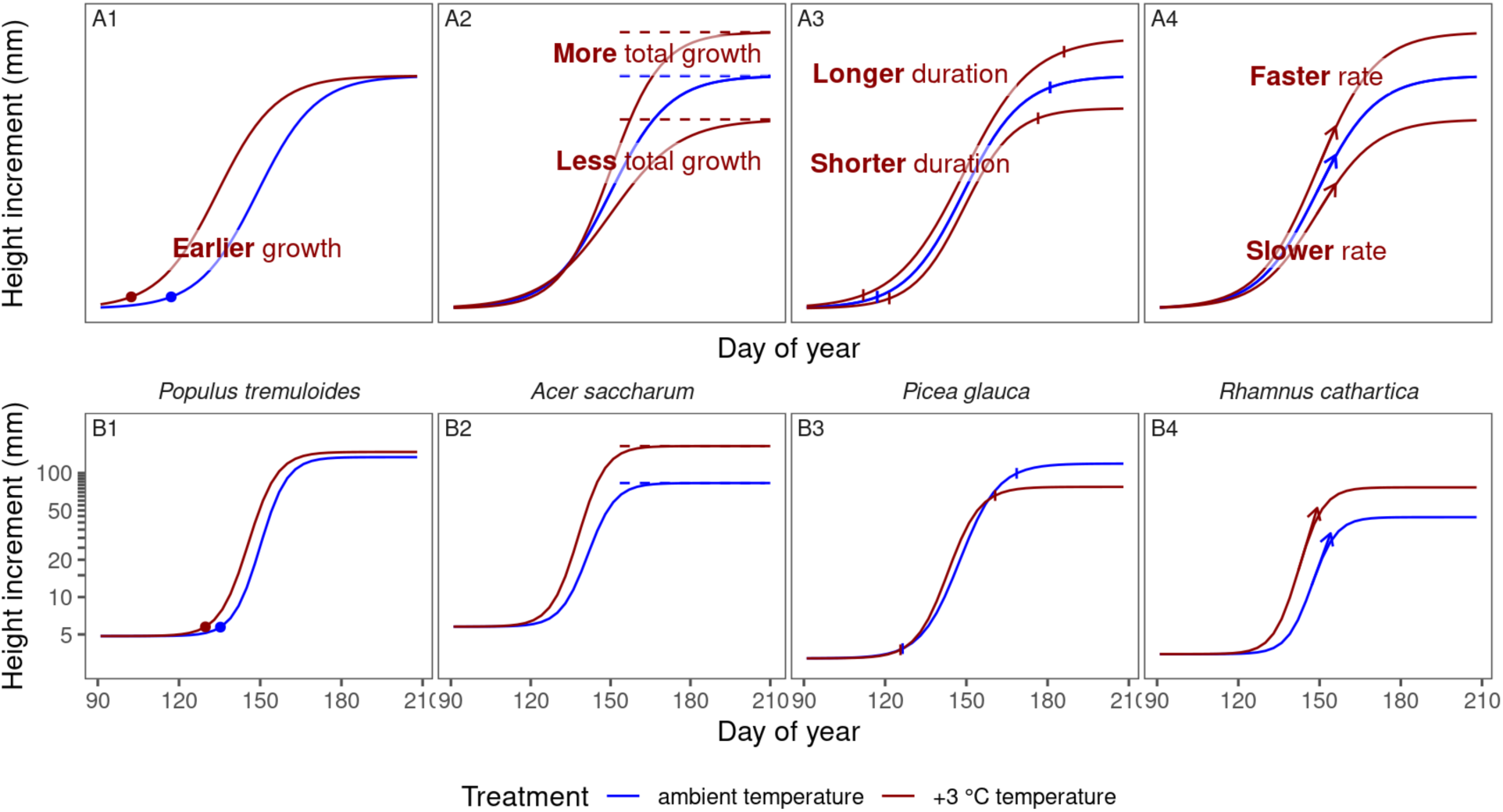
**(A)** Schematic representation of hypotheses: climate warming temporally advances height growth, leading to more, similar, or less annual total growth, through possible changes in duration and rate of growth. Logistic growth curves show modeled height increments (mm) on a log scale. Phenological and growth parameters are annotated: points indicate the start of growth (time of 5% logarithmic growth); horizontal dashed lines indicate the annual total growth (mm; visualized on a log scale); pairs of vertical segments indicate the duration of growth (from 5% to 95% logarithmic growth; day); arrows indicate the maximum relative growth rate (day^−1^). **(B)** Modeled curves of height growth in example species under ambient temperature and 3 °C warming, including (B1) trembling aspen (*P. tremuloides*) with earlier start of growth under warming, (B2) sugar maple (*A. saccharum*) with more annual total growth under warming, (B3) white spruce (*P. glauca*) with shorter duration of growth under warming, and (B4) common buckthorn (*R. cathartica*) with faster relative growth rate under warming, with all examples in open canopy under ambient rainfall. Modeled curves are marginal predictions from fitted Bayesian hierarchical models. Refer to Figs. S6-S8 for modeled curves of all species.

## Results

### Earlier growth, but not always more growth

We inferred the effects of experimental warming on both the timing and magnitude of height growth by fitting Bayesian hierarchical models (Fig. S5-8) (K. Zhu et al., 2018). Here and throughout, we modeled the effects of warming as linear against treatment strength (Fig. S9) and stationary over years (Fig. S10) after validation. Across all 20 species, experimental warming consistently advanced the timing of height growth, estimated as the timing of 5% growth in the log-transformed growth curve (Fig. 1B1,2A1,2B1, Table S3A,S4A). Shoot development advanced in general, with the start of growth changing by −4.32 (−9.62, 0.102, median and 95% interval here and throughout the text) days per 3 °C warming. The extent of advancement in the start of growth in response to warming was marginally significantly different among types of species (ANOVA, *p* = 0.0581), with exotic shrubs showing the greatest advancement, followed by deciduous angiosperm tree species and evergreen gymnosperm tree species. In addition to the start of growth, species consistently advanced in the midpoint and end of growth under warming (Fig. S11,S12) .

Despite a generally similar advancement in timing, species exhibited significantly different responses in annual total height growth, estimated as the asymptote of the log-transformed growth curve (ANOVA *p* < 0.0001, Fig. 1B2, Fig. 2A2,2B2, Table S3B, Table S4B), supporting our first hypothesis that warming will lead to earlier growth, but not always enhanced annual total growth. Evergreen gymnosperm species generally showed weak to negative responses per 3 °C warming, with −15.8 (−32.1, 10.2) % changes in annual total growth. The largest average reduction of 35.1% was seen in white spruce (*Picea glauca*) in the open canopy, under ambient rainfall. In contrast, native deciduous angiosperm species generally showed weak to positive responses per 3 °C warming, with 19.3 (−3.32, 88.0) % changes in annual total growth. The largest increase of 98.7 % was seen in sugar maple (*A. saccharum*) in the open canopy, under ambient rainfall. Non-native shrub species generally showed positive responses, with 37.4 (−14.2, 118) % changes in annual total growth, with the largest increase of 120 % in tatarian honeysuckle (*Lonicera tatarica*) in the open canopy, under reduced rainfall.

The earlier start of growth, and contrasting responses in annual total growth, were found across ambient and reduced summer rainfall, as well as across open and closed canopy conditions (Fig. 2). Eleven species were planted in all three environmental contexts along a water limitation gradient from closed canopy with ambient rainfall, to open canopy with ambient rainfall, and to open canopy with reduced rainfall. In several species, the warming effects on annual total growth were less positive in more water-limited contexts, such as jack pine (*P. banksiana*), eastern white pine (*P. strobus*), trembling aspen (*P. tremuloides*), and common buckthorn (*R. cathartica*). Some species, however, showed more positive response in annual total growth to warming in more water-limited contexts, including paper birch (*B. papyrifera*) and red maple (*A. rubrum*).

Similar patterns in growth responses to warming inferred from the Bayesian hierarchical models were consistent with those from a NLS-LME two-stage approach, based on independent individual-level growth curve fittings followed by a regression model for the estimated parameters for the growth curves (Note S3, Figs. S13-17). These results support our hypothesis that warming advances the timing of growth, but they reveal that growth enhancement is not universal, with contrasting responses among species types.

**Figure 2.**
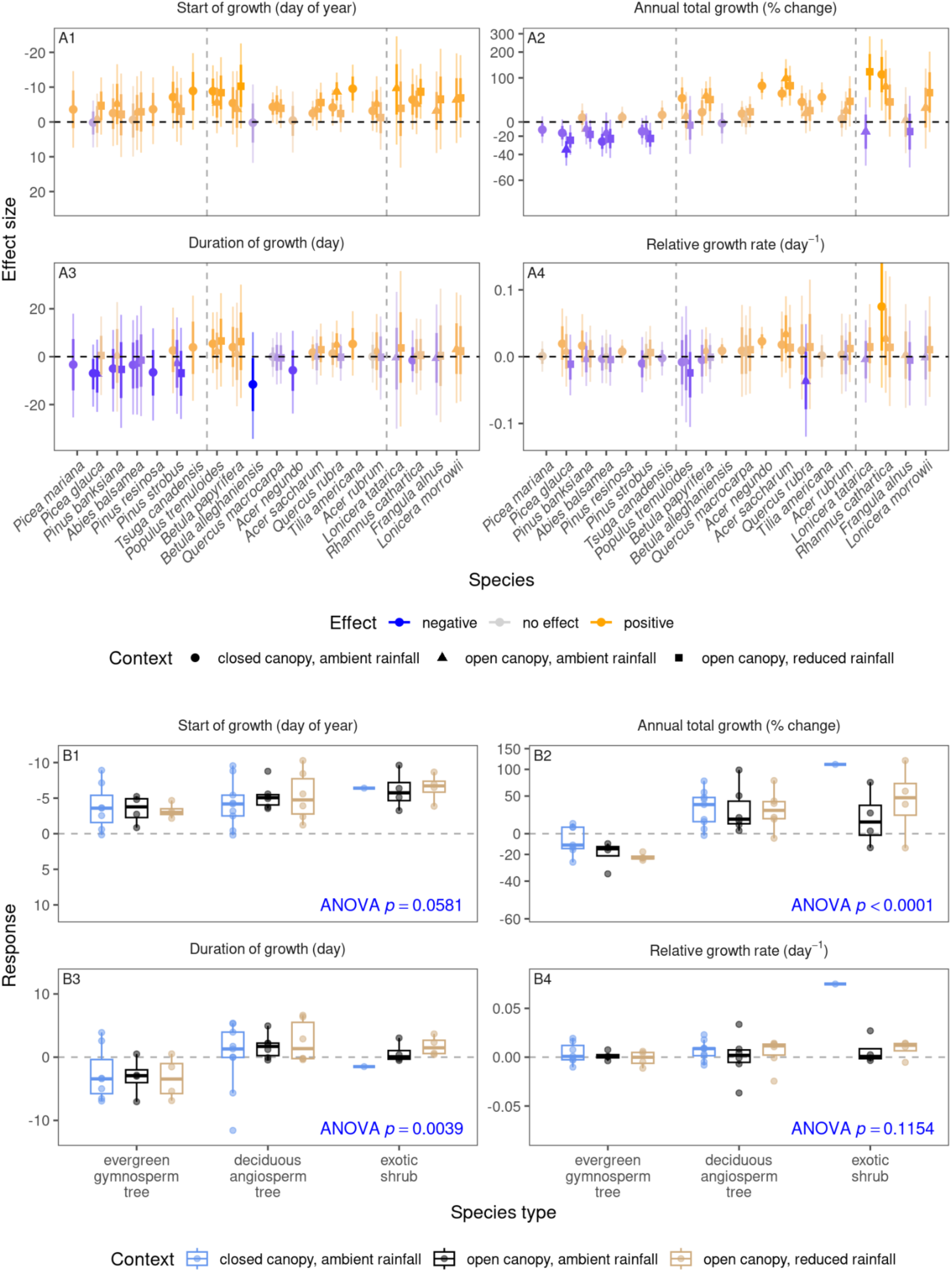
Effects per 3 °C warming on the start of growth (time of 5% logarithmic growth; day of year; effect sizes shown on a reversed scale to highlight advancement), annual total growth (mm; effect sizes on a log scale shown as equivalent percentage changes), duration of growth (duration from 5% to 95% logarithmic growth; day), and maximum relative growth rate (day^−1^) under various environmental contexts. Effect sizes were inferred with Bayesian hierarchical models. **(A)** Points show medians of effect sizes, with thick and thin error bars showing 68% and 95% credible intervals of effect sizes, respectively. Colors indicate the direction and magnitude of warming effects on growth, with positive effects associated with earlier growth, higher annual total growth, longer duration, and faster rate. Species are sorted by species types (evergreen gymnosperm trees, deciduous angiosperm trees, and exotic shrubs), and then by increasing species’ temperature niche. **(B)** Species’ height growth responses to warming treatment are summarized into three species types across environmental contexts. Boxplots show the median (line), interquartile range (box), and 95% quantile range (whiskers) of species’ posterior median estimates. *p*-values are from ANOVA for the predictor of species type, derived from linear mixed-effects models.

### Suppressed through duration, enhanced through rate and duration

To test our second hypothesis, we inferred the effects of warming on the duration and rate of growth. From the fitted models, we estimated the duration of growth as the time interval between 5% and 95% logarithmic growth, and we estimated the maximum relative growth rate, which is the maximum rate of increase per unit of existing height increment (Fig. 1, Note S2). Warming had significantly different effects on growth duration among species types (ANOVA *p* = 0.0039, Fig. 2A3,2B3, Tables S3C,S4C). Evergreen gymnosperm trees responded negatively with compressed duration (−3.32; −7.01, 3.44 day) while deciduous angiosperm tree species (1.30; −8.62, 6.50 day) had differing responses. Warming had non-significant to positive effects on relative growth rate among the 20 species (ANOVA *p* = 0.1154, Fig. 2A4,2B4, Tables S3D,S4D). While evergreen gymnosperm trees had little response (0.000822; −0.0109, 0.0183 day^−1^), deciduous angiosperm trees (0.00838; −0.0306, 0.0283 day^−1^) and exotic shrubs (0.0105; −0.00491, 0.0654 day^−1^) tended to increase in relative growth rate.

We further identified how contrasting growth responses among the 20 species were driven by the combined changes in duration or rate. We summarized how species’ responses in annual total growth to warming can be partitioned into responses in duration of growth and relative growth rate using a conceptual diagram (Fig. 3A): (I) enhanced growth driven by longer duration of growth, (II) enhanced growth driven by faster relative growth rate, (III) suppressed growth driven by shorter duration of growth, and (IV) suppressed growth driven by slower relative growth rate.

We found distinct clustering of phenology-growth responses among evergreen gymnosperm and deciduous angiosperm trees that was maintained across environmental contexts (Fig. 3B). Evergreen gymnosperm tree species predominantly fell into Type III, showing suppressed annual total growth driven by shorter duration, with weak and inconsistent changes in growth rate (Figs. 3B, S6-8). An example of Type III is balsam fir (*Abies balsamea*). Evergreen gymnosperm species occasionally fell into type IV, such as white spruce (*P. glauca*) in the open canopy, under reduced rainfall, showing suppressed annual total growth driven by slower rate.

Deciduous angiosperm tree species fell into either Type II or Type I, occasionally between the two types (Figs. 3C, S6-8). Bur oak (*Q. macrocarpa*) and sugar maple (*A. saccharum*), both with determinate shoot growth, represent those in Type II, showing enhanced annual total growth driven by faster rate, with weak and inconsistent changes in duration. Trembling aspen (*P. tremuloides*) and paper birch (*B. papyrifera*), both with indeterminate shoot growth, represent those that fell into Type I, showing enhanced annual total growth driven by longer duration, with little to negative changes in rate. Exotic shrub species exhibited a range of responses from Type I to Type II, including both faster rate and longer duration in the open canopy, under reduced rainfall.

These findings provide a mechanistic explanation for contrasting growth responses under warming mediated by phenology: the suppressed growth in evergreen gymnosperm trees was generally driven by shorter duration, whereas the enhanced growth in deciduous angiosperm trees and exotic shrubs tended to be driven by faster rate, longer duration, or their combinations.

**Figure 3.**
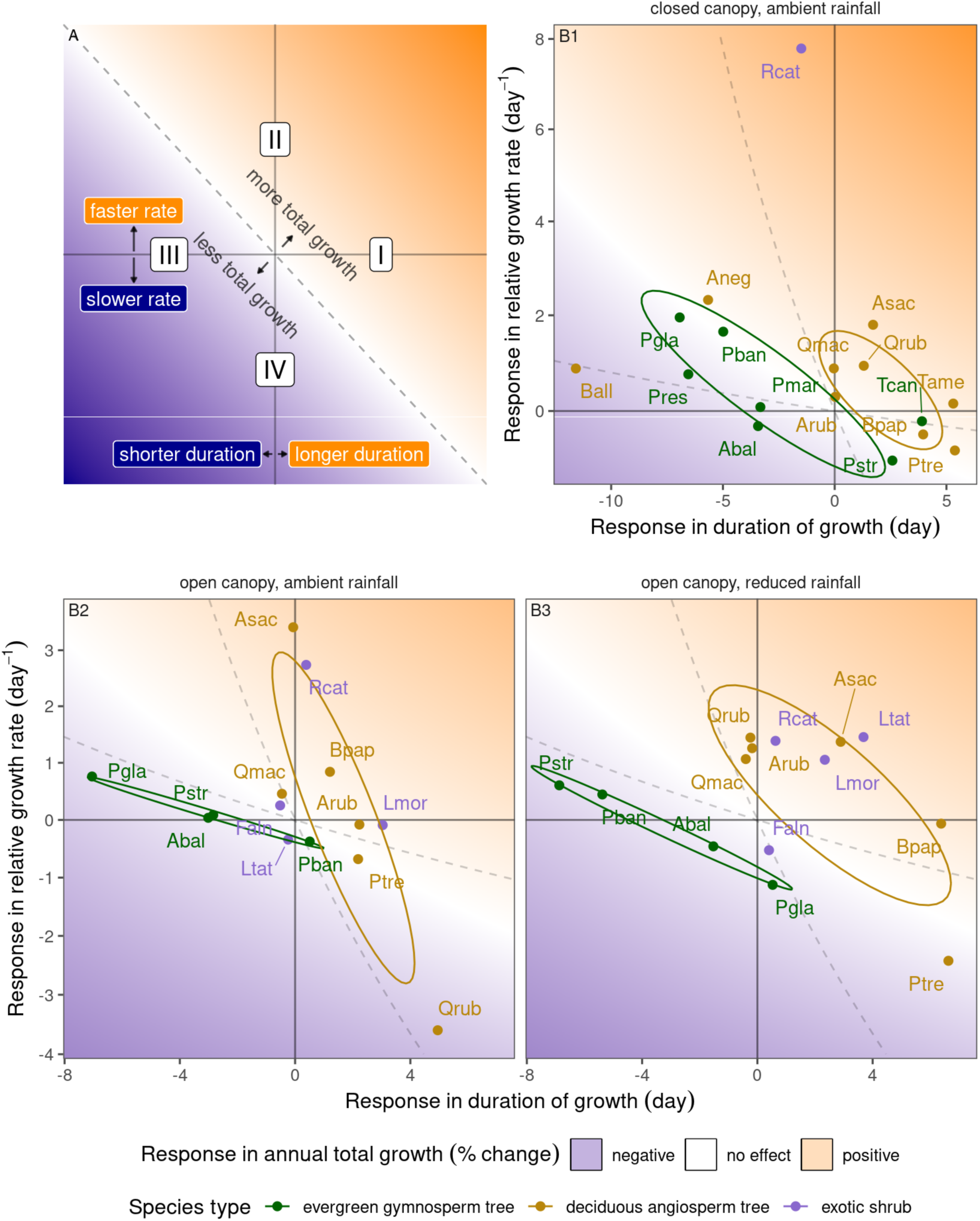
**(A)** Conceptual diagram illustrating how species’ responses in annual total growth to warming can be partitioned into responses in duration of growth and relative growth rate, summarized into four response types: (I) enhanced growth driven by longer duration, (II) enhanced growth driven by faster rate, (III) suppressed growth driven by shorter duration, and (IV) suppressed growth driven by slower rate. The dashed line represents an isoline where combined changes in duration and rate result in no net change in annual total growth. This isoline is different for each species, given their duration and rate in ambient temperature. **(B)** Relationship between species’ inferred responses in maximum relative growth rate (day^−1^) and responses in duration of growth (duration from 5% to 95% logarithmic growth; day) per 3 °C warming, under different environmental contexts. Points indicate median effect sizes for each species. Background colors represent simulated responses in annual total growth (percentage change) given varying combinations of changes in duration and rate, averaged across species. Dashed-line pairs delineate the range of species-specific isolines with no net change in annual total growth. Ellipses represent 50% concentration regions (assuming multivariate normal distributions) for evergreen gymnosperm species and deciduous angiosperm species that were in all three environmental contexts. Species acronyms are provided in Table S1.

### Conservative species at a disadvantage under warming

We examined whether species’ lifestyles reflected through their distributions and functional traits could help explain their growth responses to experimental warming. The start of growth advanced in all species under warming, and the extent of advancement differed negligibly to modestly among species varying in distributions and traits (Fig. S18,S19, Table S5A). In contrast, we found that more positive responses in annual total growth under warming were correlated with distributions and traits that reflect a more acquisitive lifestyle. In terms of distribution, species with an origin from temperate (as opposed to boreal) biomes, a more northern geographical range, a distribution in warmer areas (high temperature niche), and an exotic (as opposed to native) origin had more positive responses in annual total growth (Fig. 4, Fig. S20, Table S5B). In terms of functional traits, species with a deciduous (as opposed to evergreen) leaf habit, an indeterminate (as opposed to determinate) shoot growth type, and larger and more N-rich leaves had more positive responses in annual total growth (Fig. 4, Fig. S21, Table S5B). Variables related to moisture, including precipitation niche and drought tolerance did not strongly predict growth responses to warming (Figs. S20-21, Table S5B). These relationships generally held across different canopy and rainfall treatments.

Species distribution and functional traits predicted growth enhancement under warming through longer duration of growth and faster relative growth rate, but in different ways (see Fig. 4 for following examples). For example, contrasting growth responses between temperate and boreal species were predominantly driven by different responses in relative growth rate (Figs. S20A,S22A,S24A). Different growth responses depending on species’ temperature niche were driven by combined responses in duration of growth and relative growth rate (Figs. S20E,S22E,S24E). The contrast between determinate and indeterminate growth types was explained mostly by different responses in duration of growth (Figs. S21B,S23B,S25B).

These results show that conservative species might be at a disadvantage under future climate change due to more negative responses in annual total growth, duration of growth, and sometimes relative growth rate. The results can be interpreted in light of the phenological rank and growth rank among species. In ambient temperature, species with a more conservative lifestyle (e.g., evergreen gymnosperm trees), compared to those with a more acquisitive lifestyle (e.g., deciduous angiosperm trees and exotic shrubs), tended to have earlier start of growth, longer duration of growth, slower relative growth rate, and higher annual total growth (Fig. S26, Table S6). While experimental warming generally maintained the order in their time of height growth (Fig. S27), it shuffled the order in duration of growth and annual total growth (Fig. S28,S29), placing conservative species at a disadvantage.

**Figure 4.**
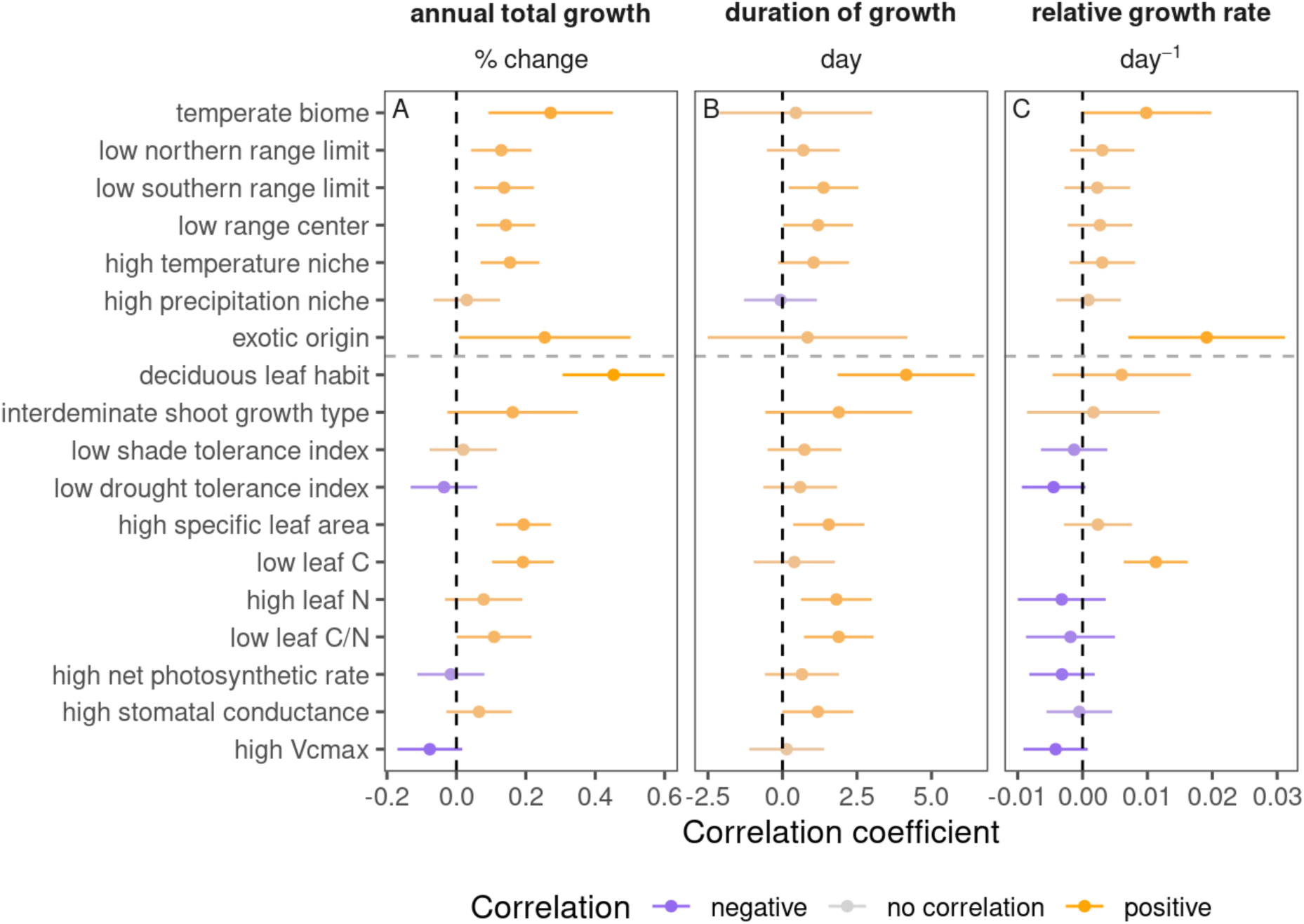
Summary of correlations between species’ distributions and functional traits with their **(A)** responses in annual total growth (% change), **(B)** responses in duration of growth (day), and **(C)** responses in maximum relative growth rate (day^−1^) per 3 °C warming. Signs of correlation coefficients were adjusted to reflect the effect of distributions and traits associated with a more acquisitive lifestyle. Correlation coefficients quantify either differences between discrete trait categories or effects per one standard deviation change in continuous trait values.

## Discussion

Our study reveals that while the phenology (i.e., timing) of spring growth consistently advanced under experimental warming under a diverse group of species, this did not consistently lead to greater annual total growth; instead, responses contrasted among species and among species types. Evergreen gymnosperm tree species tended to experience suppressed growth due to shorter duration of growth with little compensation in growth rate, whereas deciduous angiosperm tree species and exotic shrub species often showed maintained or enhanced annual total growth under warming, mainly due to accelerated growth rate and extended duration of growth. More broadly, these patterns reflect how acquisitive species derive greater growth advantages from warming compared to conservative species. These contrasting responses challenge the assumptions that an earlier start of the growing season universally enhances growth and highlight how warming may shift competitive dynamics within mixed-species forests.

The consistent advancement in spring phenology under warming has been widely reported (Parmesan & Yohe, 2003; Wolkovich et al., 2012). On the contrary, the contrasting responses in growth were only partly consistent with previous studies focused on growth, none of which, however, address the questions of interest in the same way as our study. Previous spatiotemporal analyses generally suggested increased height growth correlated with higher temperatures for species such as trembling aspen (*P. tremuloides*), black spruce (*P. mariana*), common beech (*Fagus sylvatica*), and sessile oak (*Quercus petraea*) (Bontemps et al., 2012; Messaoud & Chen, 2011). In our findings, trembling aspen (*P. tremuloides*) exhibited enhanced height growth under warming, whereas black spruce (*P. mariana*) exhibited suppressed height growth, suggesting a possible difference in responses from observational and experimental settings: observational studies often cannot disentangle the effects of covariation in moisture, ecotypes, plant age, or other sources of variation, which our experiment realistically incorporates or accounts for. Our finding partly aligns with findings of no overall change in diameter growth of temperate deciduous trees under warming (Gao et al., 2022), but our results further reveal substantial and predictable interspecific variation in growth responses. These variations are consistent with previous analyses from the B4WarmED experiment of subsets of the data used herein, which similarly reported contrasting responses in photosynthesis and stem biomass growth among species (Reich et al., 2015). Mechanistically, enhanced growth under warming may result from alleviated thermal limitations on carbon acquisition and wood formation (Gao et al., 2022; Reich et al., 2022), while suppressed growth may be explained by increased evaporative demand (Gao et al., 2022; Mirabel et al., 2023; Reich et al., 2018).

Our findings reconcile the apparent paradox of earlier phenology not always resulting in greater growth by disentangling growth into its duration and rate components. In the case of height growth, while the onset of growth was generally advanced by warming, so was the cessation of growth. As a result, the duration of growth was often unchanged or even compressed (Fig. 2B3). This finding is inconsistent with previous findings that earlier budburst was associated with longer durations of leaf expansion in several deciduous species, including red maple (*A. rubrum*), sugar maple (*A. saccharum*), and red oak (*Q. rubra*) (Klosterman et al., 2018). It is also inconsistent with the correlation between early forest green-up and longer duration of green-up derived from satellite remote sensing (Klosterman et al., 2018), which, however, might be driven by differences in the phenology of leaf development and height growth, as well as interspecific differences in phenology (R. A. Montgomery et al., 2020). These results, in the context of the global greening literature, suggest that despite the extension in overall growing season within a year, the duration of periods critical to woody growth might even shorten, resulting in the negative growth responses of evergreen species. However, note that the duration of height growth we examined here is not the duration of the photosynthetic season. It is possible that extended canopy photosynthesis with earlier cessation of height growth might lead to enhanced growth in diameter, root biomass, or branch shoots through altered carbon allocation (Canham et al., 1999; McKown et al., 2016; R. Montgomery, 2004). On the other hand, the maximum relative height growth rate was often maintained or increased under warming, contributing to the positive growth response of deciduous species. Our analysis of the maximum relative growth rate during active height growth is not directly comparable with the few studies on growth rate or green-up velocity during fixed time windows, which might be confounded by the advancement in start and end of phenological development (Bronson et al., 2009; Hong et al., 2022). The interplay between potentially shorter duration and faster rate explains the contrasting growth responses we observed: some species gained, others lost, depending on which effect dominated.

We further revealed the influence of drying in interaction with warming, both being integral aspects of the composite global change driver of climate change. Consistent with effects of climate warming on soil moisture, some of our warming treatments often induced soil drying (Liang et al., 2024; Reich et al., 2018) (Fig. 5, Table S2). While we could not directly disentangle the direct (thermal) and indirect (e.g., drying) effects of warming, we were able to examine how warming effects on phenology and growth depend on environmental context (Fig. 2, Table S4). For most species, warming was more likely to enhance growth in the closed canopy, under ambient rainfall, where soil moisture was rarely low, and more likely to suppress growth in the open canopy, under reduced rainfall, where soil moisture was often low. This could be because the closed canopy likely buffered the microclimate and reduced evapotranspiration, alleviating water limitation (soil moisture is consistently higher in this treatment), whereas the reduced precipitation treatments directly imposed water limitation (Fig. 5).

**Figure 5.**
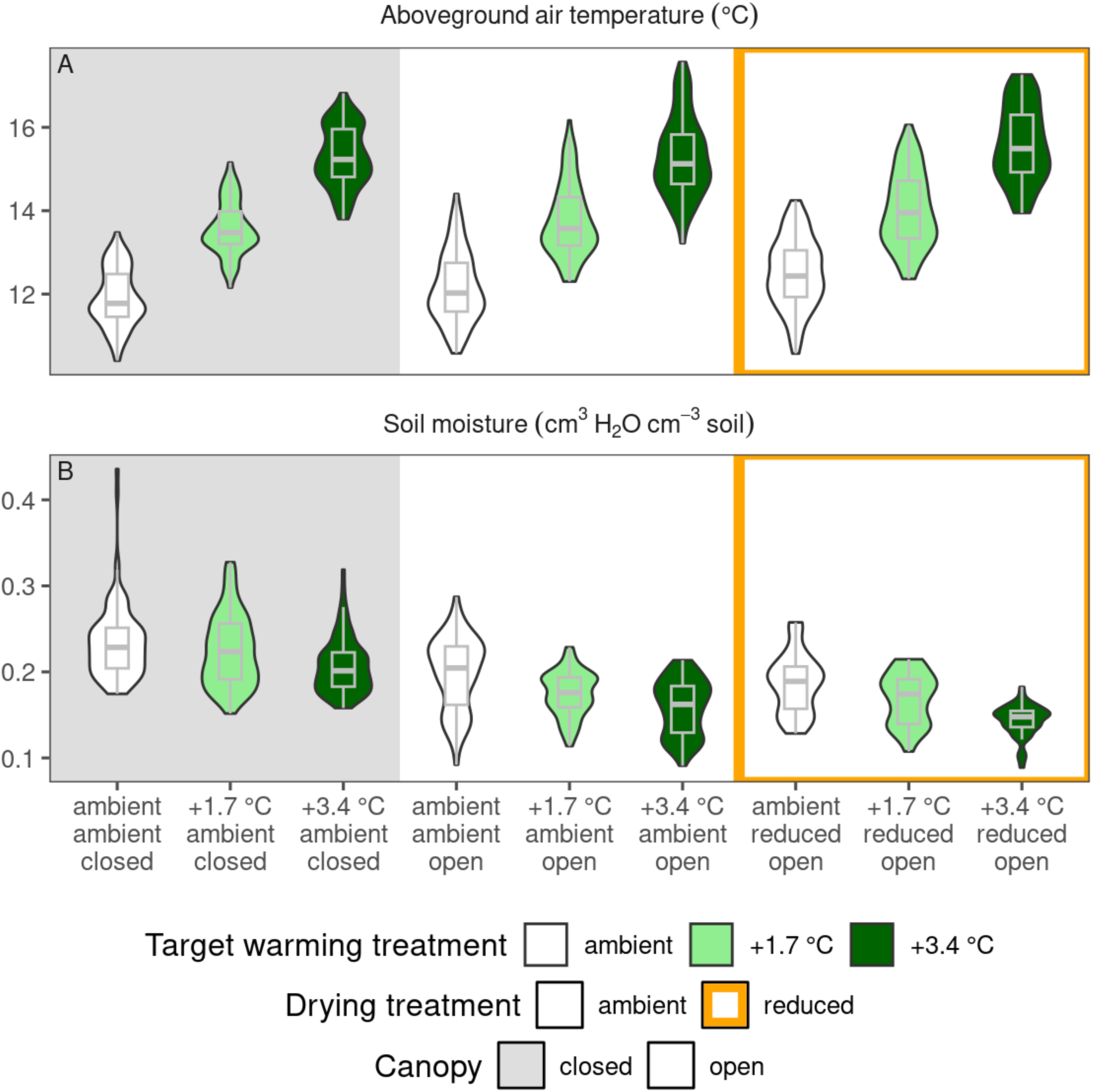
Environmental conditions, including **(A)** aboveground air temperature (°C) and **(B)** soil moisture (mm^3^ H_2_O / mm^3^ soil) under different target warming treatments, rainfall treatments, and canopy conditions. Environmental variables were measured hourly in each experimental plot. For each plot and year, we averaged environmental variables during the time period when the experimental heating system was on (generally March to October, Fig. S3), spanning the plants’ growing season.

Our findings highlight how species’ lifestyles, reflected through their distributions and functional traits, mediate their growth and phenological responses to warming. Previous studies have shown that acquisitive tree species, while having concentrated and rapid periods of development, do not necessarily grow taller than conservative tree species under harsh environmental conditions in the field (Augusto et al., 2025), which is validated by our results (Fig. S26). With a series of species’ properties, encompassing climatic niches and leaf traits, we were able to predict species’ phenological and growth responses to warming (Fig. 4, Fig. S18-25) (Pérez-Ramos et al., 2020). We validated (Fig. S20D) and expanded (Figs. S22D,S24D) the previous understanding that growth responses differ by species’ geographical range (Reich et al., 2015, 2022), demonstrating distinct plant strategies under warming. We reveal that acquisitive species might gain an advantage in growth under warming, while conservative species might be at a disadvantage. One possible explanation is that acquisitive species are better able to capitalize on brief periods of favorable conditions through rapid resource uptake and allocation to growth, while conservative species may be more constrained by their slower physiological rates and longer development windows (Dhaila et al., 1995). Our results suggested that warming might diminish or reverse, rather than maintain or reinforce, the existing growth rank (Fig. S28), consistent with prior experimental and observational results (Fisichelli et al., 2012; Reich et al., 2015, 2022). By altering the growth rank among co-occurring species, warming has the potential to shift competitive dynamics and community composition (Bontemps et al., 2012; Chamberlain & Wolkovich, 2021), having important implications for future forest biodiversity under climate change.

We acknowledge several caveats that should be considered when interpreting these results. First, despite being the rare experiment that warms in a controlled fashion aboveground and belowground without chambers, our warming treatments were paused during winter months. This could have limited our ability to detect phenological delays driven by insufficient winter chilling (Ettinger et al., 2020; Suonan et al., 2017), although it is unlikely that chilling would have been reduced enough to become limiting. Second, our study focused on seedlings and saplings rather than adult trees, which differ in phenology by ontogenetic stage (Mediavilla & Escudero, 2009) or position in the canopy (Augspurger & Bartlett, 2003), due to different ecological strategies, light conditions, and microclimate. Additional data, such as forest inventory and fluxes, could validate whether shifts in seedling phenology transfer to adult trees, and further scale up to influence community dynamics and ecosystem functioning.

Overall, our findings advance understanding of juvenile regeneration dynamics and projecting future forest composition, particularly in biomes with frequent natural and anthropogenic disturbances. We anticipate that warming will not universally enhance growth, but instead restructure seasonal dynamics, shift competitive hierarchies, and alter community composition. The differential changes could be understood with the interplay of changes in duration and rate of growth, and predicted by species’ lifestyles. These insights highlight the value of long-term global change experiments in providing realistic and generalizable predictions of forest responses to climate change.

## Methods

### Study sites

This study was conducted at two research sites in northern Minnesota as part of the Boreal Forest Warming at an Ecotone in Danger (B4WarmED) experiment, a long-term, open-air global change experiment established in 2006 (Fig. S1A). In this study, we use data collected from 2010 to 2023. The sites were approximately 150 km apart, spanning the transition zone between temperate and boreal biomes. The southern site is located at the Cloquet Forestry Center (CFC; 46°40′46″ N, 92°31′12″ W; 382 m a.s.l.; mean annual temperature (MAT) from 2010 to 2023 4.78 ± 1.20 °C; total annual precipitation (TAP) 912 ± 178 mm; mean ± standard deviation), and the northern site is at the Hubachek Wilderness Research Center (HWRC; 47°56′46″ N, 91°45′29″ W; 415 m a.s.l.; MAT 3.84 ± 1.19 °C; TAP 716 ± 107 mm). Both sites are characterized by coarse-textured upland soils. Experimental plots were scattered in 40 to 60 y old mixed aspen–birch–fir stands at the start of the experiment.

### Experimental treatments

We employed an incomplete factorial design (Fig. S2) that crossed site (CFC and HWRC), canopy condition (closed, open), warming treatment (targets of ambient, +1.7 °C, +3.4 °C), and drying treatment when under open canopy (ambient, 40% rainfall reduction for the summer period from June 1 to September 30) (Stefanski et al., 2020), resulting in a 2 × 2 × 3 (× 2) factorial structure. Each combination was replicated across three blocks, leading to a total of 72 plots (3 m diameter, 7.1 m² each; Fig. S1A,S2, Table S1). Canopy conditions reflect differences in habitat under natural and anthropogenic disturbances: closed-canopy plots received approximately 5–10% of full sunlight, while open-canopy plots received 40–60%.

Open-canopy blocks were created through overstory harvests and brush clearing prior to plot establishment. To simulate future climate warming, both soil and air temperatures were elevated using a combination of aboveground and belowground heating methods. Air warming was achieved using overhead ceramic infrared heating elements (Model FTE-1000, 240V, 245 mm × 60 mm; Mor Electric Heating Assoc.), mounted in open-air conditions without enclosures to preserve natural precipitation and airflow. Soil warming was implemented via resistance heating cables (146 m, 240V, GX, Devi A/B, Denmark) installed by hand at 10 cm and 20 cm depths in a concentric spiral within each warmed plot. These systems provided continuous heating generally from March to October (other than brief interruptions for a variety of causes), to achieve near the average target increases of +1.7 °C and +3.4 °C relative to the ambient plot in the same block, matching projected warming scenarios over the next 75 to 100 years. The two levels of warming treatments raised aboveground air temperature by +1.62 °C and +3.26 °C, respectively (Fig. 5, Table S2) during the periods when treatments were deployed. To account for possible effects from the experimental setup, ambient plots were also equipped with non-functional heating elements and PVC tubes. Drying treatments were applied to a subset of plots using manually operated rainout shelters. These shelters, mounted on frames above each plot, were extended manually during individual rainfall events in the summer from Jun 1st to Sep 30th, to intercept and exclude precipitation. This approach reduced total annual precipitation input by approximately 40%, simulating the drier growing season conditions expected under future climate scenarios. As rainout shelters could not be installed under a closed canopy, the three temperature treatments were replicated in each closed canopy block. The factorial combination of warming, drying, and canopy conditions enables direct testing of individual and interactive effects of multiple global change drivers on forest trees.

### Seedling planting

We planted seedlings of multiple tree and shrub species common to the boreal-temperate ecotone (Table S1). Among these, 11 species received the complete factorial design as described in Fig. 1, while the other species were planted in a subset of treatment combinations. All were two-year-old bare-root seedlings, sourced through the Minnesota Department of Natural Resources nursery program using local seed stock collected within 80 km of the Cloquet site. Initial plantings occurred in May 2008. Since then, harvesting and replanting have been performed in multiple years (2012, 2014, 2017) to replace individuals that had grown too tall, exceeding the height of the aboveground heating elements. There were replacements of individuals lost due to disturbance, herbivory, disease, or other reasons in some other years (2011, 2013, and 2020) to maintain a consistent number of seedlings and species representation in each plot. On average, each seedling remained in the plot for three years.

### Height growth measurement

To study the phenology of seedling height growth in spring, we measured the length of terminal shoots, which offers unique information beyond discrete phenophase status data on the continuous trajectory of development. Two individuals per species per plot were marked as “phenology trees” and monitored regularly throughout the growing season. Height increment was measured from the base of the current-year terminal shoot to the terminal shoot apex (Fig. S1B). Measurements began in early spring when warming treatments were initiated and continued through the end of leaf expansion in midsummer. During spring and fall, measurements were taken every 3–4 days, and weekly from mid-June to mid-August. Observers received standardized training and regular calibration by long-term staff to ensure accuracy and consistency in measurements.

### Data pre-processing

We pre-processed the height increment data to improve data quality in several steps.

1. We removed records with non-positive growth values (likely errors) or missing seedling identifiers (barcodes).
2. To remove the effects of mortality, we excluded all data points associated with a seedling in a year if the seedling had been marked as dead at any point in the year.
3. We grouped data into three environmental contexts defined by canopy condition and rainfall treatment: (1) closed canopy, ambient rainfall, (2) open canopy, ambient rainfall, and (3) open canopy, reduced rainfall.
4. We used the measured temperature increase (°C) as the covariate representing warming treatment (Table S2). For each plot and year, we averaged aboveground air temperature during the time period when the experimental heating system was on (generally March to October), spanning the plants’ growing season. We then calculated temperature increases as the difference in aboveground air temperature between each plot and the average from ambient plots in the same experimental block and year. We summarized the temperature increases by environmental contexts.
5. We created a grouping variable to represent the unique combinations of seedling identifier and year, allowing us to account for individual-level variations in growth parameters. We created another grouping variable to represent the unique combinations of site and year, to account for site-year-level variations.

### Bayesian hierarchical model

We adopted a Bayesian hierarchical modeling framework to synthesize complex and variable growth responses to warming on multiple aspects of height growth (Zhu et al., 2018). This approach provides the flexibility to model the nonlinear dynamics of height growth, explicitly accounting for variation among sites, years, and individual seedlings, and improving estimates in the presence of sparse or noisy height increment data. To infer species-specific responses, we fit a model for each species instead of pooling data from all species. Within each species, we also repeated the inference for three environmental contexts, to examine the context-dependency of warming effects. For each species and context, the model predicts the height increment *y* of an individual seedling *i*, at a site *s*, in a year *t*, on a day of year *d*. The models for all species follow the same structure, consisting of a data model, a process model, and a parameter model (Equation 1).

In the data model, we assumed that the response variable height increment *y_i,s,t,d_* followed a log-normal distribution, with mean *μ_i,s,t,d_* and variance *σ*^2^ (Eqn 1). This mean *μ_i,s,t,d_* is a latent variable that intuitively represents height increment *y* after log transformation. We used a log-normal distribution because height growth is inherently a multiplicative process exhibiting an exponential increase in height increment. The height increment variable is also non-negative and typically right-skewed, with most individuals having small or moderate growth and a few having strongly elongated growth. The log-normal distribution also provides a flexible way to accommodate heteroscedasticity in the dataset, where errors were larger when the mean values of height increment were larger (Figs. S4,S5).

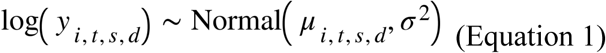

In the process model, we characterized the latent variable *μ_i,s,t,d_* as a logistic growth function over day of year *d* (Fig. 3) (Dow et al., 2022). This logistic function is controlled by minimum value *c*, midpoint *x*_0*i,s,t*_, rate *k_i,s,t_*, and amplitude *A_i,s,t_* (Eqn 2). The minimum value *c* intuitively represents the log-transformed length of the winder bud, and we assumed that it is consistent within a species. The midpoint *x*_0*i,s,t*_, also called the inflection point, represents the DOY with the fastest increase in log-transformed height increment. The rate *k_i,s,t_* is conceptually related to the intrinsic growth rate in logistic population growth, and we later used it to derive the maximum relative growth rate. The amplitude *A_i,s,t_* represents the difference between the minimum value *c* and the maximum log-transformed height increment.

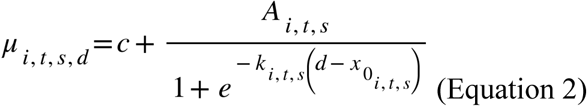

We assumed that the three key parameters (midpoint *x*_0*i,s,t*_, rate *k_i,s,t_*, and amplitude *A_i,s,t_*) that represent the timing, pace, and magnitude of growth are specific to each individual seedling, leading to variations in height growth curves among individuals (Eqn 3). We assumed that these individual-level parameters follow normal distributions, centered around the sum of an intercept *μ_p_*, fixed effects *δ_p,i_*, and random effects *α_p,s,t_*, with *p* indicating one of the three parameters of interest (Eqn 3). Note that we modeled rate *k_i,t,s_* with log transformation, as it is an inherently positive and right-skewed parameter.

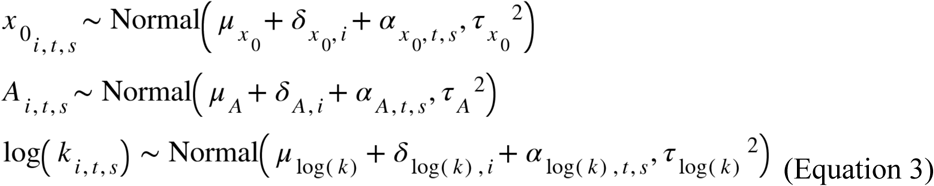

The parameter model specified the structures of fixed effects and random effects. We modeled fixed effects *δ_p,i_* as a linear function of the warming treatment *W_i_* received by seedling *i* (Eqn 4). Warming treatment was the measured temperature increase compared to the ambient plots (°C). The coefficients *β_p_* were the focus of interpretation, allowing us to infer the effects of treatments on key aspects of height growth.

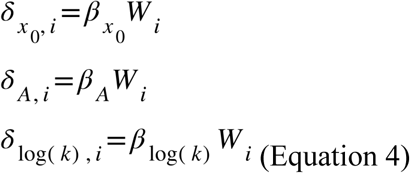

Random effects *α_p,s,t_* were normally distributed among combinations of site *s* and year *t*, to account for habitat differences and interannual variations (Eqn 5).

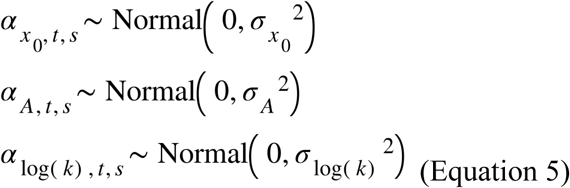

We set uniform or weakly informative normal priors for parameters (Note S1 Eqn S1).

We implemented model fitting using the *nimble* package in *R*, which provides a flexible and efficient framework for building and sampling from Bayesian hierarchical models. For each species, we ran the Markov chain Monte Carlo (MCMC) sampler in a single chain for 50,000 iterations, with a burn-in of 5,000 iterations and thinning every 5 steps.

### Model evaluation and post-processing

We evaluated the performance of the Bayesian hierarchical models through several approaches. First, we assessed MCMC convergence by inspecting trace plots and posterior distributions. We then compared conditional predictions with observed height growth to evaluate predictive accuracy, calculating Spearman’s rank correlation coefficient (*ρ*). To further examine model behavior, we generated marginal predictions across all combinations of treatments. After confirming both convergence and predictive performance, we proceeded to interpret the estimated coefficients.

To facilitate a more intuitive interpretation of model results, we processed the MCMC chains post-hoc to derive the effects of treatments on a few ecologically meaningful metrics: start of growth, end of growth, duration of growth, maximum relative growth rate, and total annual growth. We approximated treatment effects by adding MCMC chains of coefficients *β_p_* to those of baseline parameters *μ_p_*, generating perturbed parameters *θ_p_*. Posterior samples of derived metrics were computed for both *μ_p_* and *θ_p_*, and their differences informed the estimated effect sizes (Note S2, Eqns S2-4).

### Detecting patterns in model parameters

To summarize patterns in treatment effects, we performed linear regressions, with the modeled effects of 3 °C warming on parameters (e.g., annual total growth) as dependent variables. We used species type, environmental context, and their interactions as independent variables. Species type was coded as a three-level categorical variable with levels of “evergreen gymnosperm trees” (reference level), “deciduous angiosperm trees,” and “exotic shrubs.” Environmental context was coded as a three-level categorical variable: “closed canopy, ambient rainfall,” “open canopy, ambient rainfall” (reference level), and “open canopy, reduced rainfall,” such that we can easily interpret the effects of closed canopy and reduced rainfall, respectively. After fitting the linear model, we further assessed the overall effect of species type with ANOVA. To summarize the parameters of species without warming treatment, we repeated a similar analysis with the baseline parameters as response variables.

To understand the relationship between growth and phenology, we placed species’ responses in a space described by responses in duration of growth and responses in relative growth rate (both per 3 °C warming) across environmental contexts. We summarized the responses of the three species types with 50% concentration ellipses, assuming multivariate normal distributions. We inspected if the three groups fall into one of the four hypothesized response types.

### Predicting responses with species’ lifestyle

We assembled two types of variables that likely reflect the acquisitive or conservative lifestyle of species.

First, we used species’ distributions, including their biome (boreal or temperate), northern range limit (°N), southern range limit (°N), range center (°N), temperature niche (°C), precipitation niche (mm), and origin (exotic or native). Range limits and centers were defined with methods in Reich et al. (2015). We calculated species’ climatic niche that represent the climatic conditions they are associated with in their distribution (Feeley et al., 2020; Zhu et al., 2024). We did this by (1) retrieving occurrence records within North America from the Global Biodiversity Information Facility (GBIF) (Derived dataset GBIF.org, 2025), (2) calculating long-term mean annual temperature (MAT, °C) and total annual precipitation (TAP, mm) using climatic data from WorldClim (Fick & Hijmans, 2017), (3) retrieving long-term MAT and TAP at all occurrences of the species, (4) calculating the medians of all long-term MAT and TAP as the species’ temperature and precipitation niches.

Second, we estimated the functional traits, including leaf habit (deciduous or evergreen), shoot growth type (determinate or indeterminate), shade tolerance index, drought tolerance index, specific leaf area (mm^2^ mg^−1^), leaf carbon (C, %), leaf nitrogen (N, %), leaf C:N ratio, net photosynthetic rate (*μ*mol C m^-2^ s^−1^), stomatal conductance (*μ*mol H_2_O m^−2^ s^−1^), and maximum carboxylation rate (Vcmax, *μ*mol CO_2_ m^−2^ s^−1^). Shade and drought tolerance were extracted from Niinemets & Valladares (2006). We calculated specific leaf area from the leaf dry mass and leaf size measured as part of the B4WarmED experiment. Leaf carbon, leaf nitrogen, leaf C:N ratio, net photosynthetic rate, stomatal conductance, and maximum carboxylation rates were also measured in the B4WarmED experiment. We summarized leaf- or plant-level measurements to species-level traits by taking the mean. For traits with skewed distributions, we instead calculated the geometric means instead of arithmetic means. The transformed traits include specific leaf area, photosynthetic rate, stomatal conductance, and maximum carboxylation rate (Díaz et al., 2016).

We performed linear regression to model species’ responses per 3 °C warming (e.g., responses in annual total growth) with each of these lifestyle variables (one at a time), environmental context, and their interaction. Some lifestyle variables are two-level categorical variables. We standardized the continuous lifestyle variables (mean-centered and scaled by one standard deviation). To simplify the model and facilitate interpretation, we coded the environmental context here as an integer variable, with −1 corresponding to “closed canopy, ambient rainfall,” 0 corresponding to “open canopy, ambient rainfall,” and 1 corresponding to “open canopy, reduced rainfall.” The coefficients of this integer context variable conceptually represent the effects of water limitation on growth responses.

All analyses were performed using *R* (version 4.4.3).

## Data availability

Processed data and R code will be available upon request.

## Acknowledgments

The authors would like to thank members of the Reich and Montgomery labs at the University of Minnesota for helping to support the B4WarmED project over the 2008 to present period, and the Zhu Lab and Chen Lab at the University of Michigan, for comments on the manuscript. In particular, we thank Dr. Kara Dobson for providing data and insights during exploring the role of functional traits. Y.S. was supported by the Eric and Wendy Schmidt AI in Science Postdoctoral Fellowship, a Schmidt Sciences program. K.Z. and Y.S. were supported by the National Science Foundation [grant number 2306198 (CAREER)] and the Michigan Institute for Data and AI in Society. This research was supported by the National Science Foundation ASCEND Biology Integration Institute (NSF-DBI-2021898) and the US Department of Energy, Office of Science, and Office of Biological and Environmental Research award number DE-FG02-07ER64456; Minnesota Agricultural Experiment Station MN-42-030 and MN-42-060; and the College of Food, Agricultural and Natural Resources Sciences and Wilderness Research Foundation, University of Minnesota.

## Author contributions

Y.S., P.B.R., R.A.M., and K.Z. conceptualized this study. P.B.R. and R.A.M. conceived and designed the original B4WarmED experiment, including the collection of the long-term growth and phenology dataset presented herein. A.S. and R.B. implemented the experiment; supervised or performed acquisition of all tree growth and phenology data, as well as associated temperature and rainfall data; and curated all data. Y.S., K.Z., and Y.C. developed the methodology for modeling height growth data, including the Bayesian hierarchical models and post-hoc analyses. All authors contributed to the analytical framework. Y.S. performed the formal data analysis. All authors contributed to the interpretation of the results. Y.S. wrote the first draft of the manuscript. All authors reviewed and edited the manuscript. K.Z., Y.C., R.A.M., and P.B.R. acquired financial support for this study.

