## Supplementary Information for "Warming makes 20 boreal species grow earlier but not always taller"

**Note S1.** Priors of Bayesian hierarchical models

$$
\begin{aligned}
&c \sim \text{Uniform}(0,3) \\
&\mu_A \sim \text{Uniform}(1,6) \\
&\beta_A \sim \text{Normal}(0,0.5) \\
&\tau_A \sim \text{Truncated Normal}(0,0.5,0,\infty) \\
&\sigma_A \sim \text{Truncated Normal}(0,0.25,0,\infty) \\
&\mu_{x_0} \sim \text{Uniform}(120,180) \\
&\beta_{x_0} \sim \text{Normal}(0,5) \\
&\tau_{x_0} \sim \text{Truncated Normal}(0,5,0,\infty) \\
&\sigma_{x_0} \sim \text{Truncated Normal}(0,2.5,0,\infty) \\
&\mu_{\log(k)} \sim \text{Uniform}(-3.5,-1) \\
&\beta_{\log(k)} \sim \text{Normal}(0,0.1) \\
&\tau_{\log(k)} \sim \text{Truncated Normal}(0,0.1,0,\infty) \\
&\sigma_{\log(k)} \sim \text{Truncated Normal}(0,0.05,0,\infty) \\
&\sigma^2 \sim \text{Truncated Normal}(0,1,0,\infty) \qquad \text{(Equation S1)}
\end{aligned}
$$

**Note S2.** Deriving ecologically meaningful parameters

We derived the treatment effects on these ecologically meaningful traits in three steps.

1) For each of the original parameter $p$, including $x_0$, $A$, and $\log(k)$, we generated altered model parameters $\theta_p$ by adding the treatment coefficients $\beta_p$ multiplied by three to the baseline parameters $\mu_p$ (Eqn S2).
   $$\Delta_p = 3\beta_p$$
   $$\theta_p = \mu_p + \Delta_p \quad \text{(Equation S2)}$$
   The multiplier of three was chosen to predict the effect of 3 °C warming treatment on parameters. This step was repeated for every iteration in the MCMC chain; we omit these indices here for clarity.
2) Using both the MCMC chains of baseline parameters $\mu_p$ and the altered parameters $\theta_p$, we computed the posterior samples of baseline and altered derived parameters $q$. Annual total growth was defined as the asymptote of the growth curve. Start of growth was defined as the time of 5% logarithmic growth. End of growth was defined as the time of 95% logarithmic growth. Duration of growth was defined as the time interval between start and end of growth. Because the slope of the log-transformed growth curve is the relative growth rate (i.e., rate of increase per unit of existing height increment), maximum relative growth rate is the slope at 50% logarithmic growth. Their relationships could be derived analytically.
   $$\mu_{\text{annual total growth}} = \mu_A$$
   $$\mu_{\text{duration of growth}} = \frac{2\log(19)}{\exp(\mu_{\log(k)})}$$
   $$\mu_{\text{start of growth}} = \mu_{x_0} - \frac{\mu_{\text{duration of growth}}}{2}$$
   $$\mu_{\text{end of growth}} = \mu_{x_0} + \frac{\mu_{\text{duration of growth}}}{2}$$
   $$\mu_{\text{relative growth rate}} = \frac{\mu_A \exp(\mu_{\log(k)})}{4} \quad \text{(Equation S3)}$$
   The same relationships apply for altered parameters $\theta_q$. The equations are omitted.
3) For each posterior sample, we subtracted the derived traits under ambient conditions (from $\mu_q$) from the corresponding trait under treatment conditions (from $\theta_q$).
   $$\Delta_q = \theta_q - \mu_q \quad \text{(Equation S4)}$$

The derived MCMC chains for $\mu_q$ and $\Delta_q$ were then summarized in the same way as regular MCMC chains.

**Note S3.** Two-stage approach

Apart from the full Bayesian hierarchical models described, we used an alternative NLS-LME two-stage approach to test the robustness of our findings. This alternative approach consists of a first stage of nonlinear least squares (NLS) regression for fitting individual growth curves and a second stage of linear mixed-effects (LME) models for estimated parameters.

First, we performed NLS regression to the log-transformed growth trajectories of each combination of individual seedling and year. The model was a logistic curve with the parameters of minimum $c$, midpoint $x_0$, amplitude $A$, and rate $k$, similar to the structure of Eqn 2. Parameter estimation was performed by minimizing the residual sum of squares using the Gauss–Newton algorithm, with the maximum number of iterations of 200. With the fitted parameters, we derived ecologically interpretable parameters (e.g., total annual growth, duration of growth, relative growth rate) following Eqn. S3.

Second, we modeled the fitted individual-level parameters as response variables, warming treatment as the independent variable, and site-year combination as random intercepts. This is conceptually similar to the structure of Eqn 3-5. We fitted the LME models using the *brms* R package, where coefficients were estimated in a Bayesian framework via Hamiltonian Monte Carlo sampling implemented in Stan (Bürkner, 2017). The posterior distributions of LME coefficients, representing responses to warming treatment, were then analyzed in ways similar to the fully Bayesian hierarchical models.

This approach differed from the fully Bayesian hierarchical models in several ways. First, the NLS step produced parameter estimates independently for each seedling–year growth trajectory, without assuming any distribution of individual parameters or how responses scale between 1.7 °C and 3.4 °C target warming treatments. Results from this step served as a robustness check for the assumptions of the fully hierarchical models. Second, because many growth trajectories were incomplete or noisy, the NLS step often failed to yield stable or biologically realistic parameter estimates. This led to large variations in individual-level parameters and large uncertainties in the estimated treatment effects. By contrast, the "partial pooling" mechanism of Bayesian hierarchical models were able to "borrow strength" across seedlings, improving inference with incomplete trajectories while reducing the influence of outliers. Third, the LME step treated the NLS-derived parameters as fixed observations and only accounted for site–year variations. It therefore did not propagate uncertainty in individual-level parameter estimates to the uncertainty in treatment effects. Given these differences, we expected the Bayesian hierarchical approach and the two-stage NLS-LME approach to produce qualitatively similar parameter estimates, with the Bayesian hierarchical models offering improved efficiency in parameter estimation.

(A)

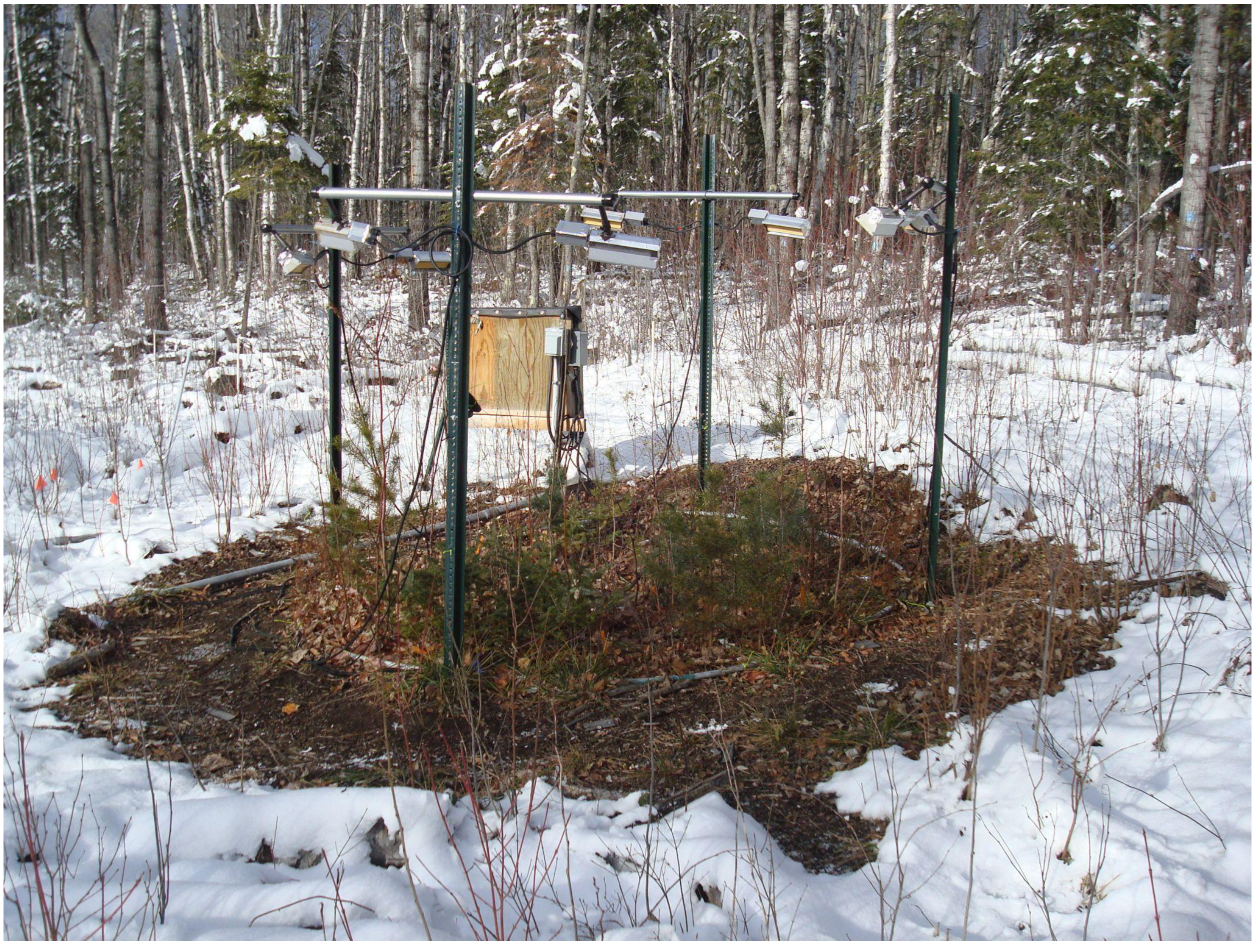

(B)

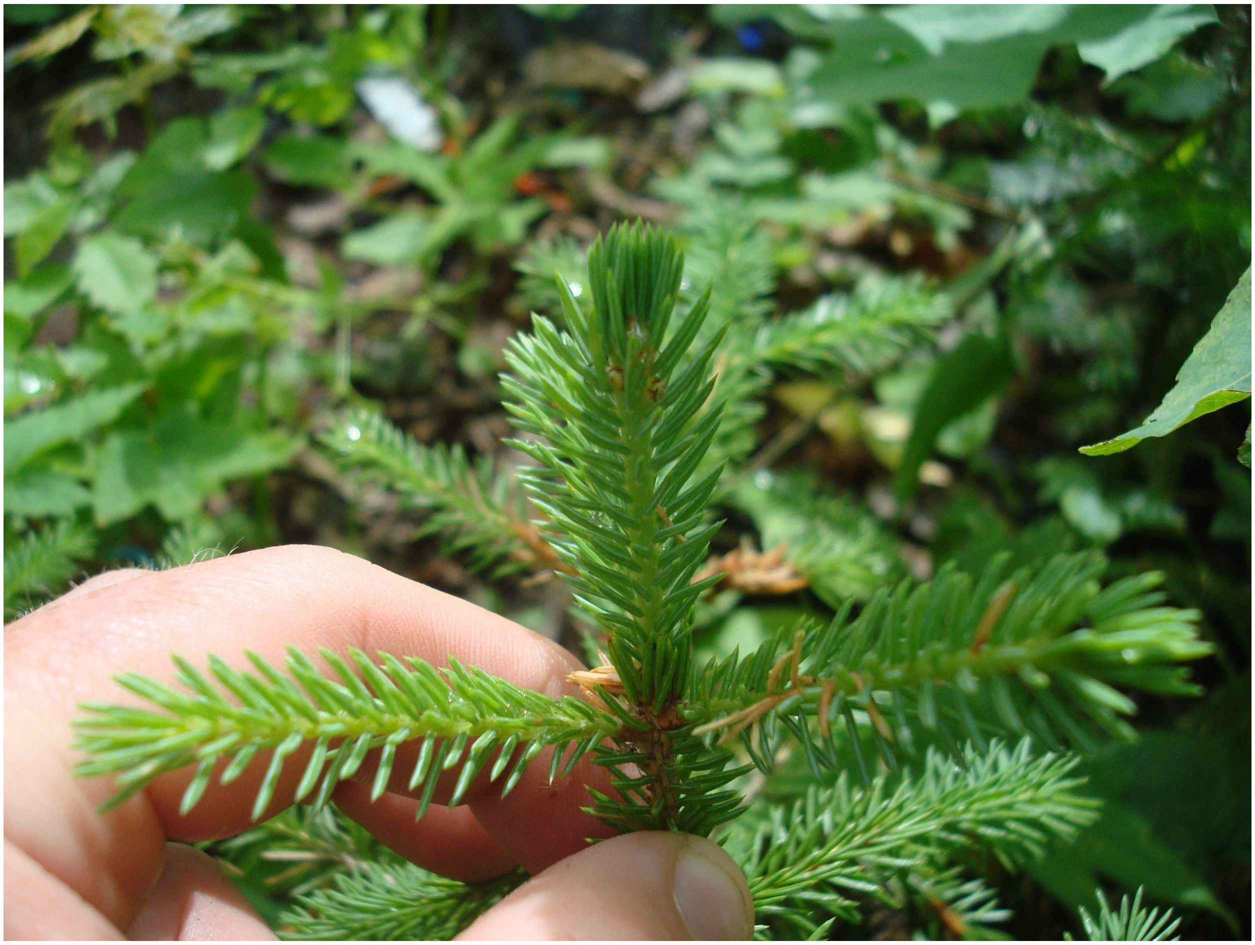

**Figure S1. (A)** 3-m diameter circular plot with boreal and temperate tree saplings planted in the B4WarmED experiment. (Photo credit: Artur Stefanski) **(B)** The difference in color between shoot segments grown in the previous year and the current year allows measurement of height growth during spring. Photo of a white spruce (*Picea glauca*) seeding on June 15, 2012. (Photo credit: Kyle Gill)

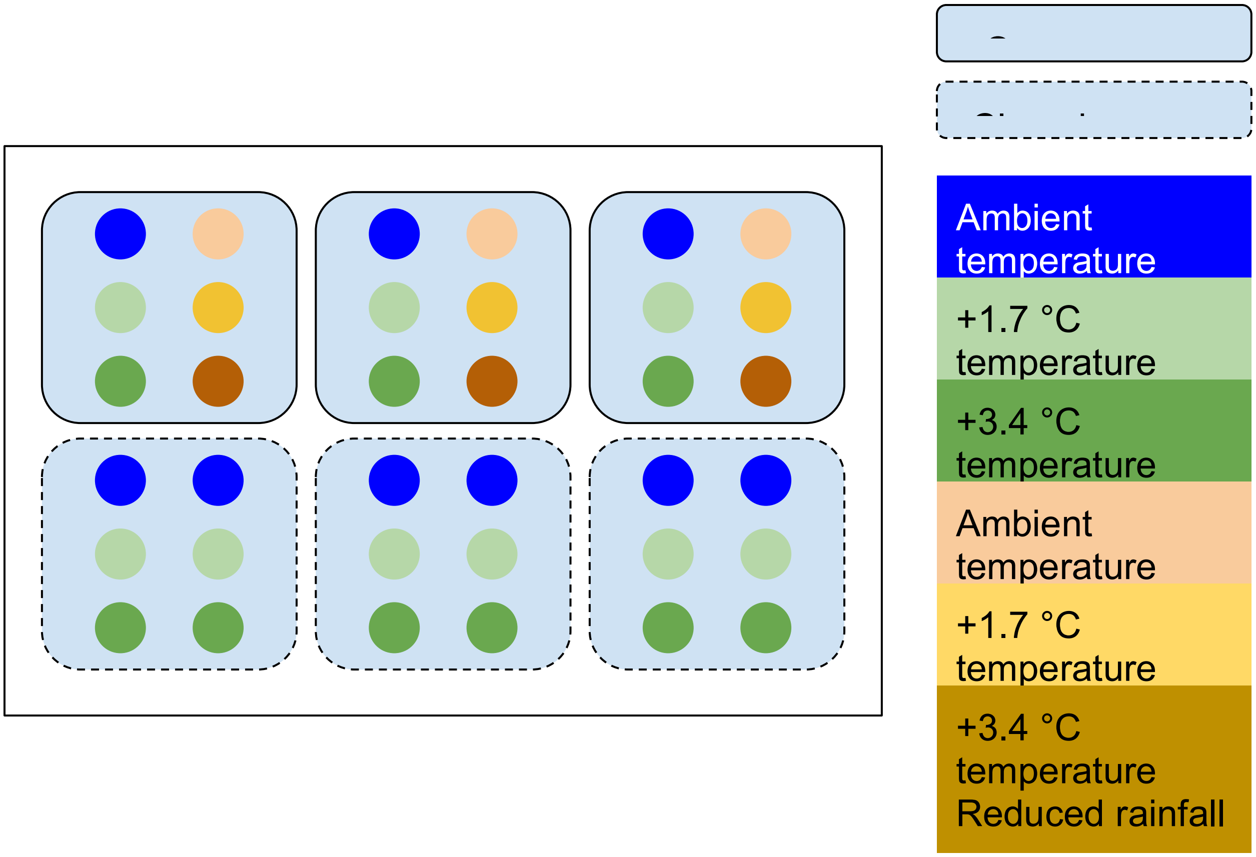


**Figure S2.** Factorial design of the B4WarmED experiment, crossing canopy condition, warming treatment, and drying treatment, with each combination replicated across three blocks. Each circle represents a circular plot. Symbols are not drawn to scale. Two experimental sites (CFC and HWRC) share the same design.

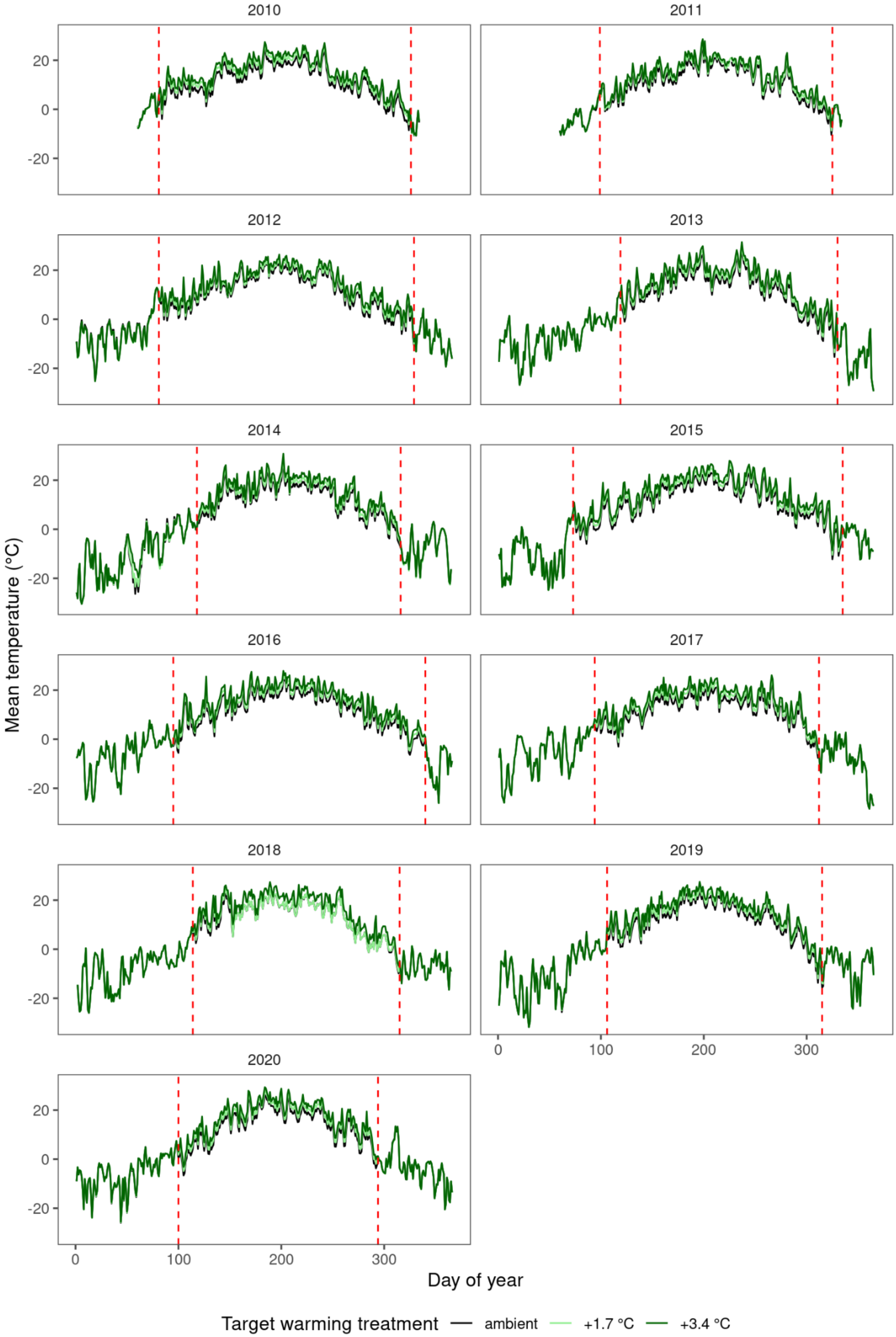

2010
2011
2012
2013
2014
2015
2016
2017
2018
2019
2020
20
0
-20
0
100
200
300
Mean temperature (°C)
Day of year
Target warming treatment
ambient
+1.7 °C
+3.4 °C

**Figure S3.** Mean aboveground air temperature (°C) under different target warming treatments, recorded with temperature loggers. Vertical lines indicate start and end of the operation of experimental heating systems.

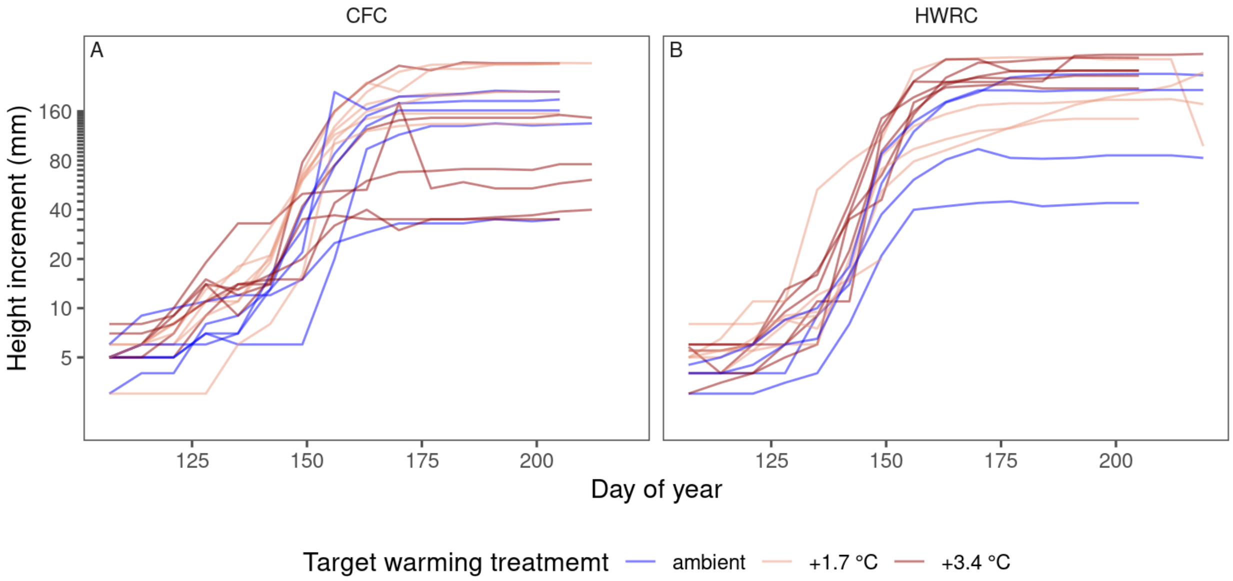


**Figure S4.** Example height increment (mm) data for red maple (*Acer rubrum*) seedlings at the Cloquet Forestry Center (CFC) and the Hubachek Wilderness Research Center (HWRC) in 2020.

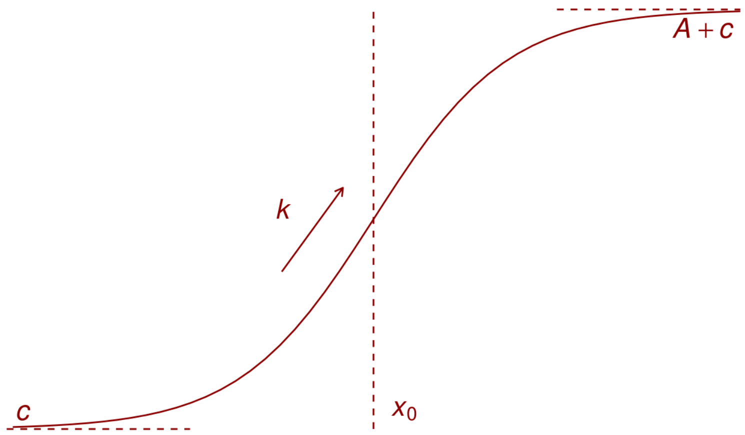


**Figure S5.** Conceptual representation of the logistic function characterizing height growth phenology, with the parameters of minimum value $c$, midpoint $x_0$, amplitude $A$, and rate $k$.

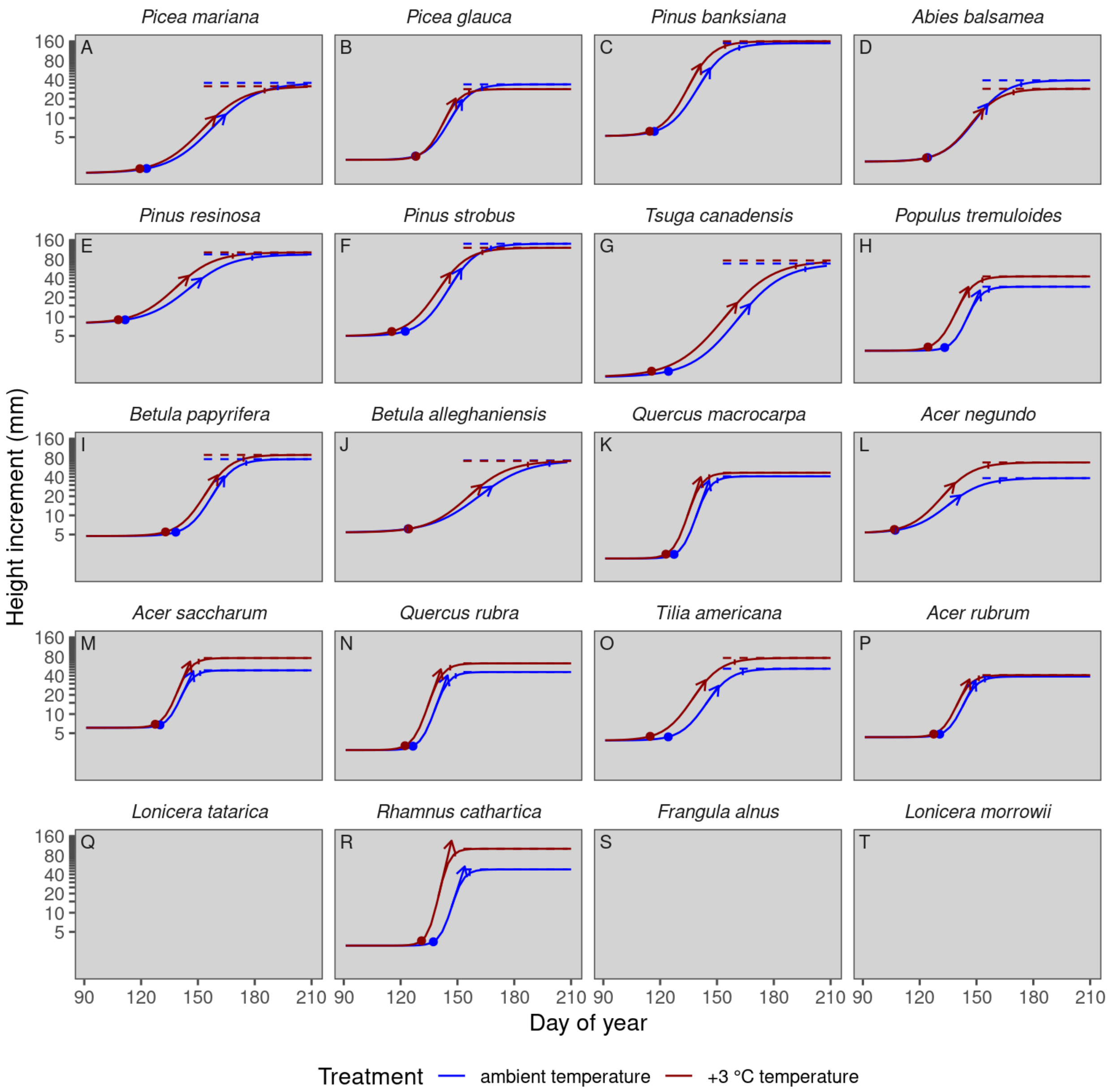


**Figure S6.** Modeled curves of height growth across species under ambient temperature and 3 °C warming, in closed canopy, under ambient rainfall. (Not all species were planted in this environmental context.) Logistic curves show modeled height increments (mm) on a log scale. Annotations follow Fig. 1. Order of species follows that in Fig. 2A.

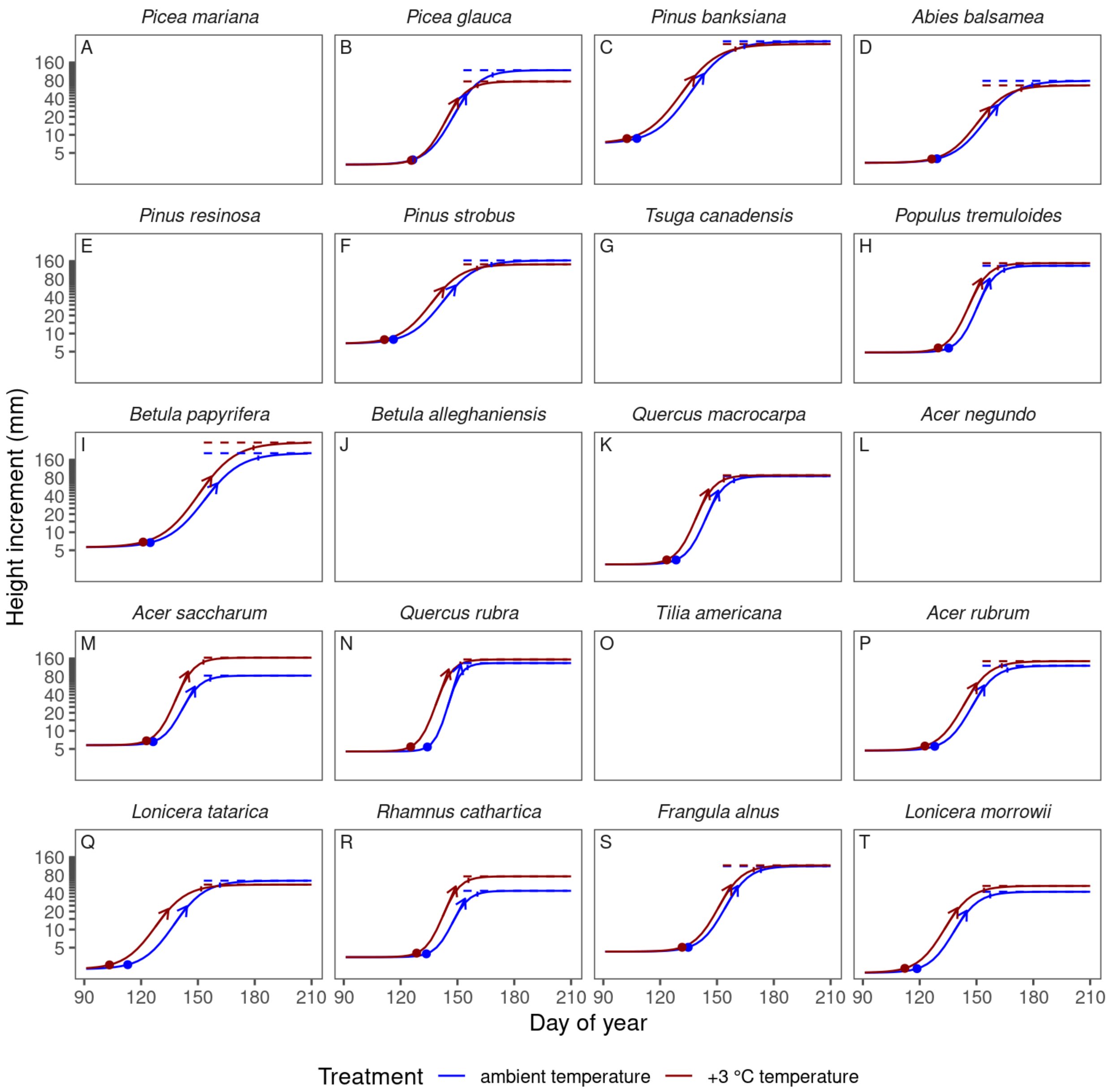


**Figure S7.** Modeled curves of height growth across species under ambient temperature and 3 °C warming, in open canopy, under ambient rainfall. (Not all species were planted in this environmental context.) Logistic curves show modeled height increments (mm) on a log scale. Annotations follow Fig. 1. Order of species follows that in Fig. 2A.

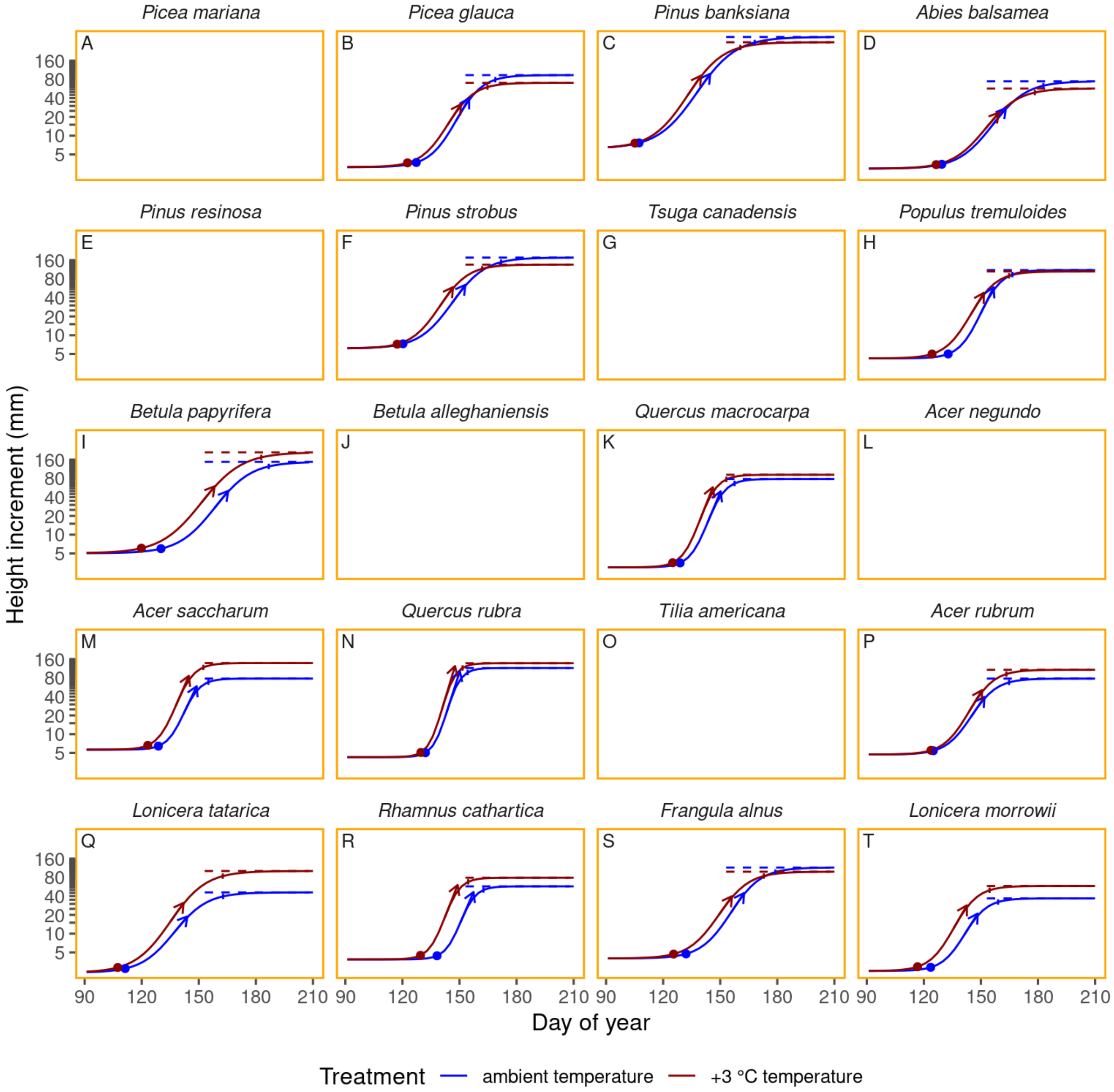


**Figure S8.** Modeled curves of height growth across species under ambient temperature and 3 °C warming, in open canopy, under reduced rainfall. (Not all species were planted in this environmental context.) Logistic curves show modeled height increments (mm) on a log scale. Annotations follow Fig. 1. Order of species follows that in Fig. 2A.

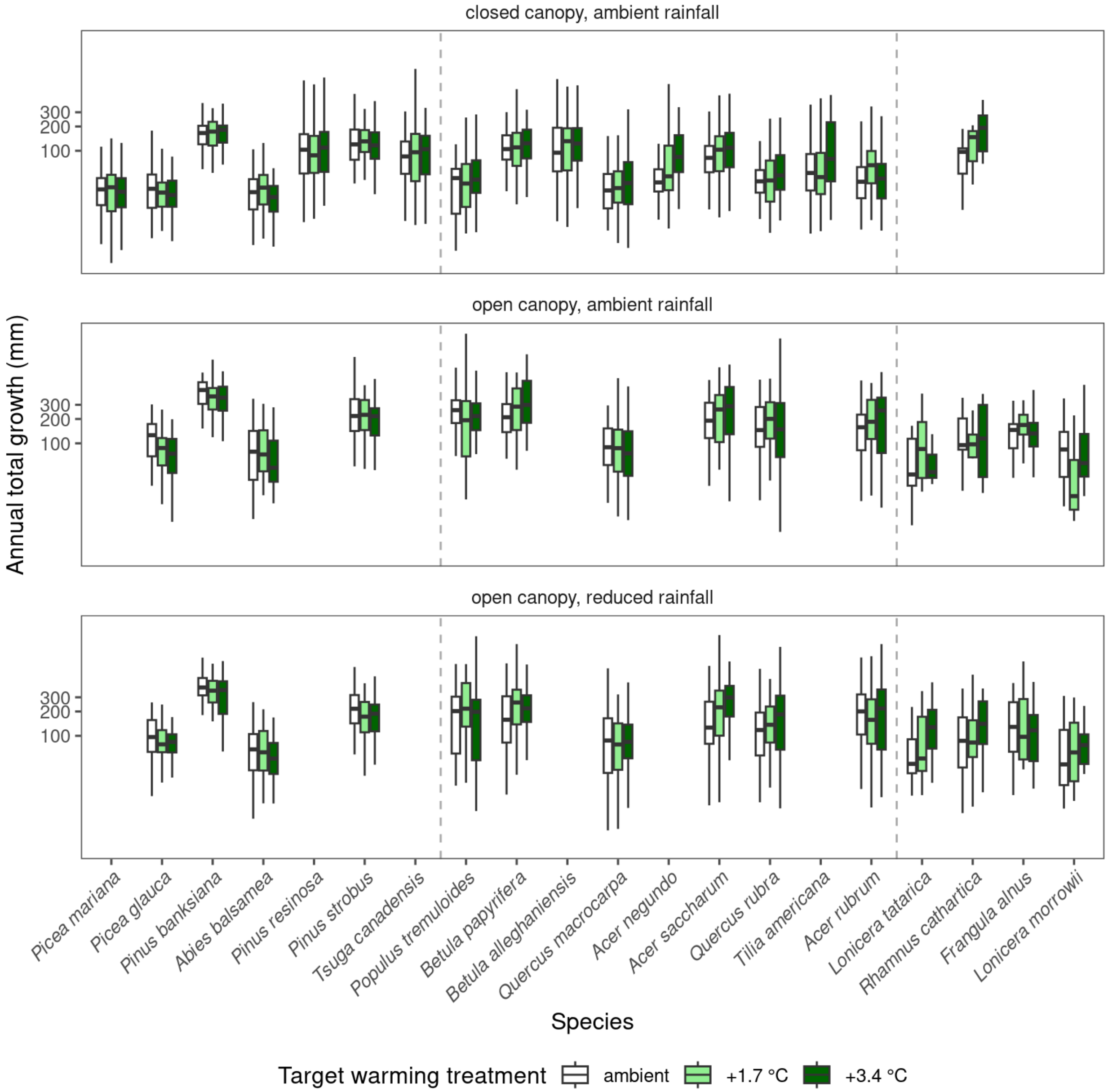


**Figure S9.** Responses in log-transformed annual total growth (mm) did not exhibit consistent nonlinearity against the strength of warming treatment across species. Boxplots summarize the annual total growth fitted with nonlinear least squares (NLS) regression for each growth trajectory.

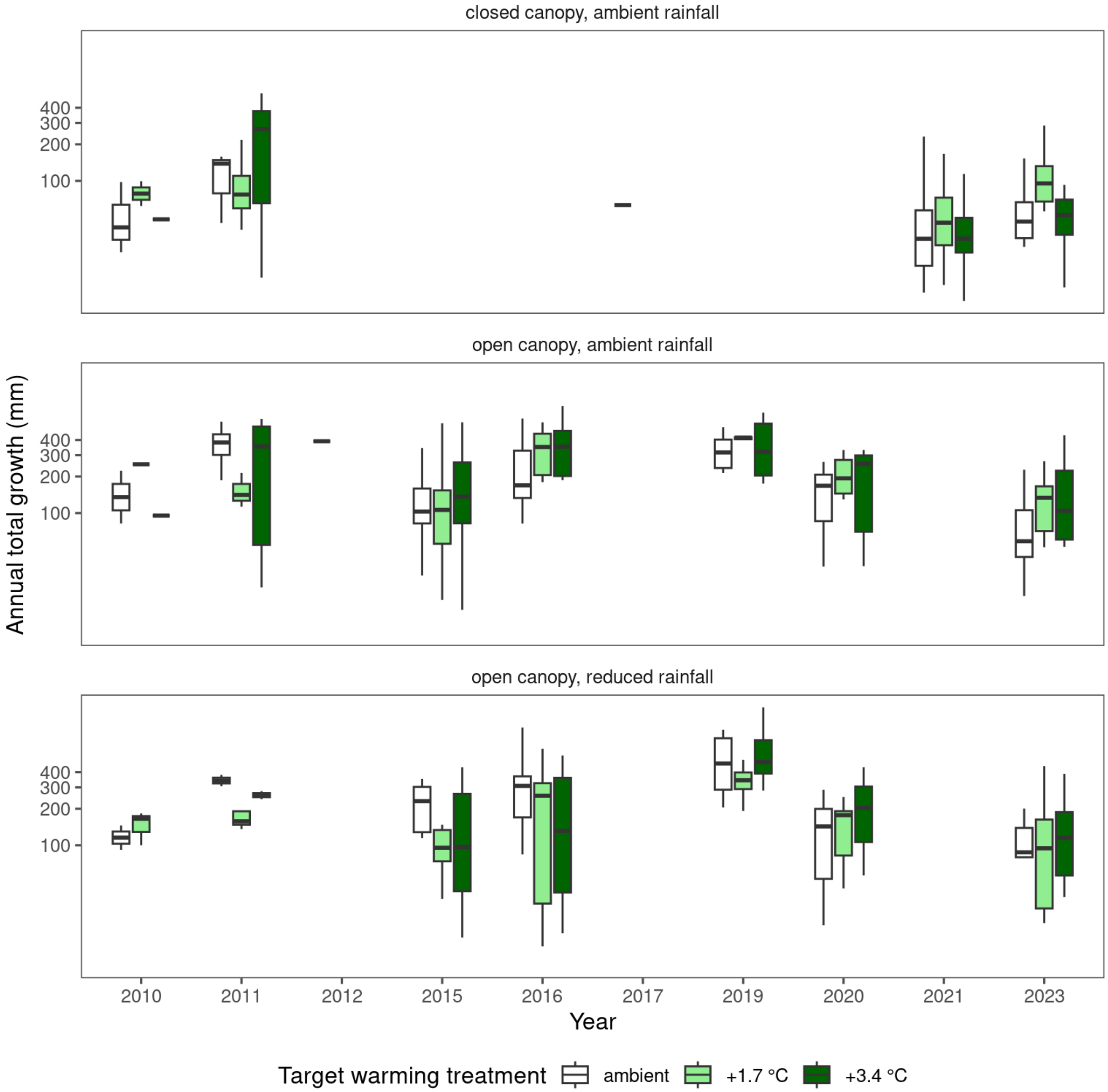


**Figure S10.** Responses in log-transformed annual total growth (mm) did not exhibit consistent trends over years. Boxplots summarize the annual total growth fitted with nonlinear least squares (NLS) regression for each growth trajectory in the example species of red maple (*A. rubrum*).

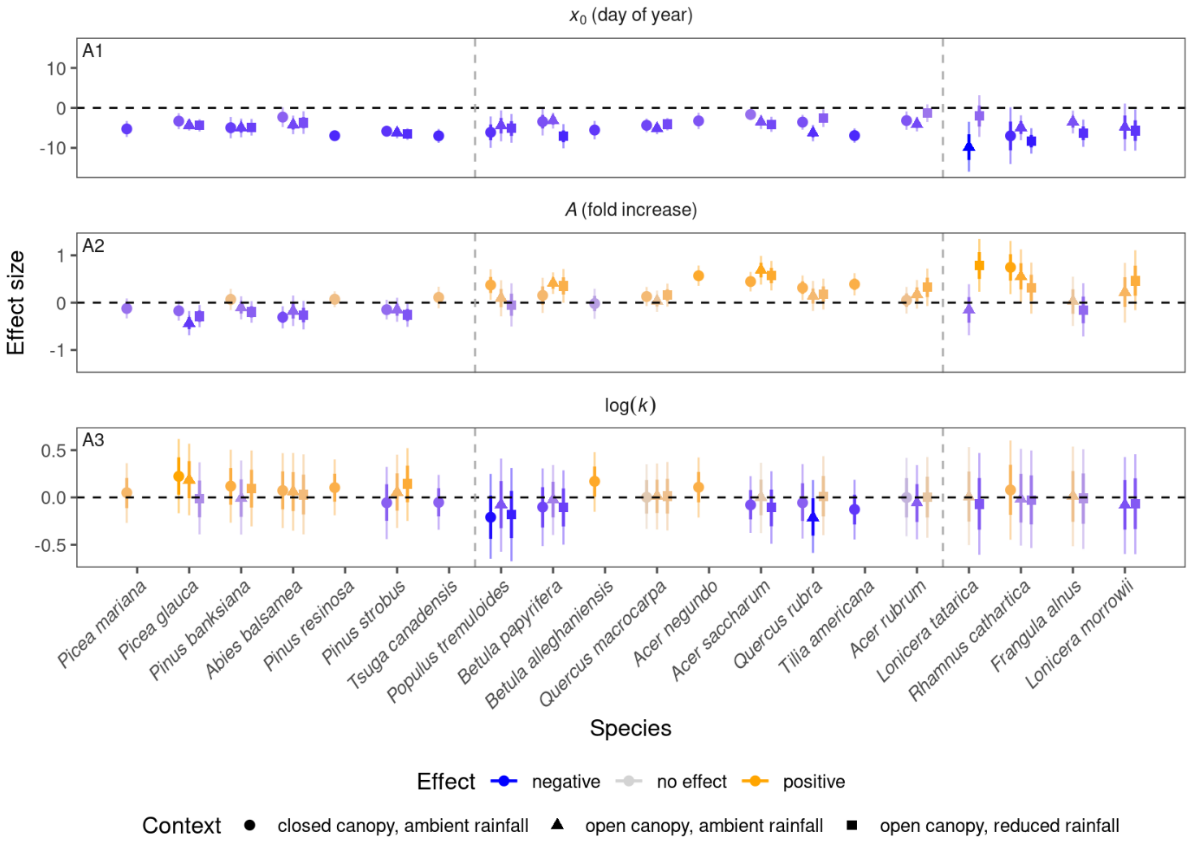

$x_0$ (day of year)
A1
A (fold increase)
A2
log(k)
A3
Effect size
Picea mariana
Picea glauca
Pinus banksiana
Abies balsamea
Pinus resinosa
Pinus strobus
Tsuga canadensis
Populus tremuloides
Betula papyrifera
Betula alleghaniensis
Quercus macrocarpa
Acer negundo
Acer saccharum
Quercus rubra
Tilia americana
Acer rubrum
Lonicera tatarica
Rhamnus cathartica
Frangula alnus
Lonicera morrowii
Species
Effect negative no effect positive
Context closed canopy, ambient rainfall open canopy, ambient rainfall open canopy, reduced rainfall


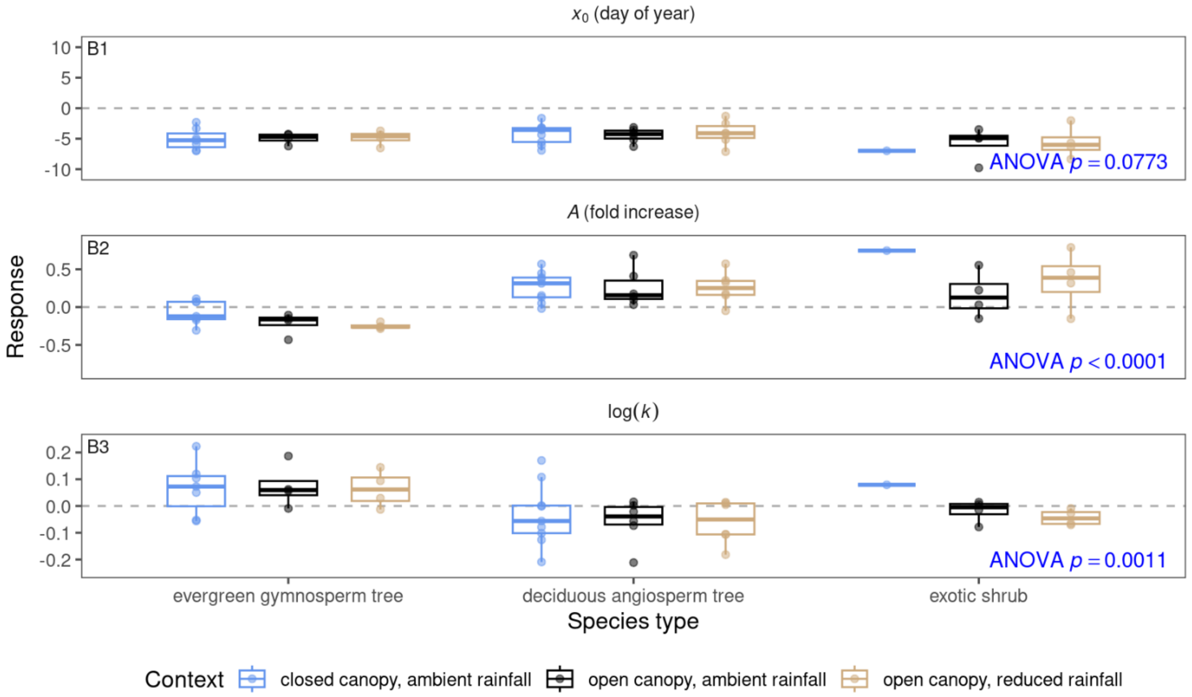

$x_0$ (day of year)
B1
ANOVA $p = 0.0773$
A (fold increase)
B2
ANOVA $p < 0.0001$
log(k)
B3
ANOVA $p = 0.0011$
Response
evergreen gymnosperm tree
deciduous angiosperm tree
exotic shrub
Species type
Context closed canopy, ambient rainfall open canopy, ambient rainfall open canopy, reduced rainfall

**Figure S11.** Effects of 3 °C warming on the parameters $x_0$ (time of 50% logarithmic growth; day of year), *A* (difference between minimum and maximum height increment on a log scale), and log(*k*) (describing the rate of height increment) under various environmental contexts. Effect sizes were inferred with Bayesian hierarchical models. **(A)** Points show medians of effect sizes, with thick and thin error bars showing 68% and 95% credible intervals of effect sizes, respectively. Colors indicate the direction and magnitude of warming effects on growth. Order of species follows that in Fig. 2A. **(B)** Species' height growth responses to warming treatment are summarized into three species types across environmental contexts. Boxplots show the median (line), interquartile range (box), and 95% quantile range (whiskers) of species' posterior median estimates. *p*-values are from ANOVA for the predictor of species type, derived from linear mixed-effects models.

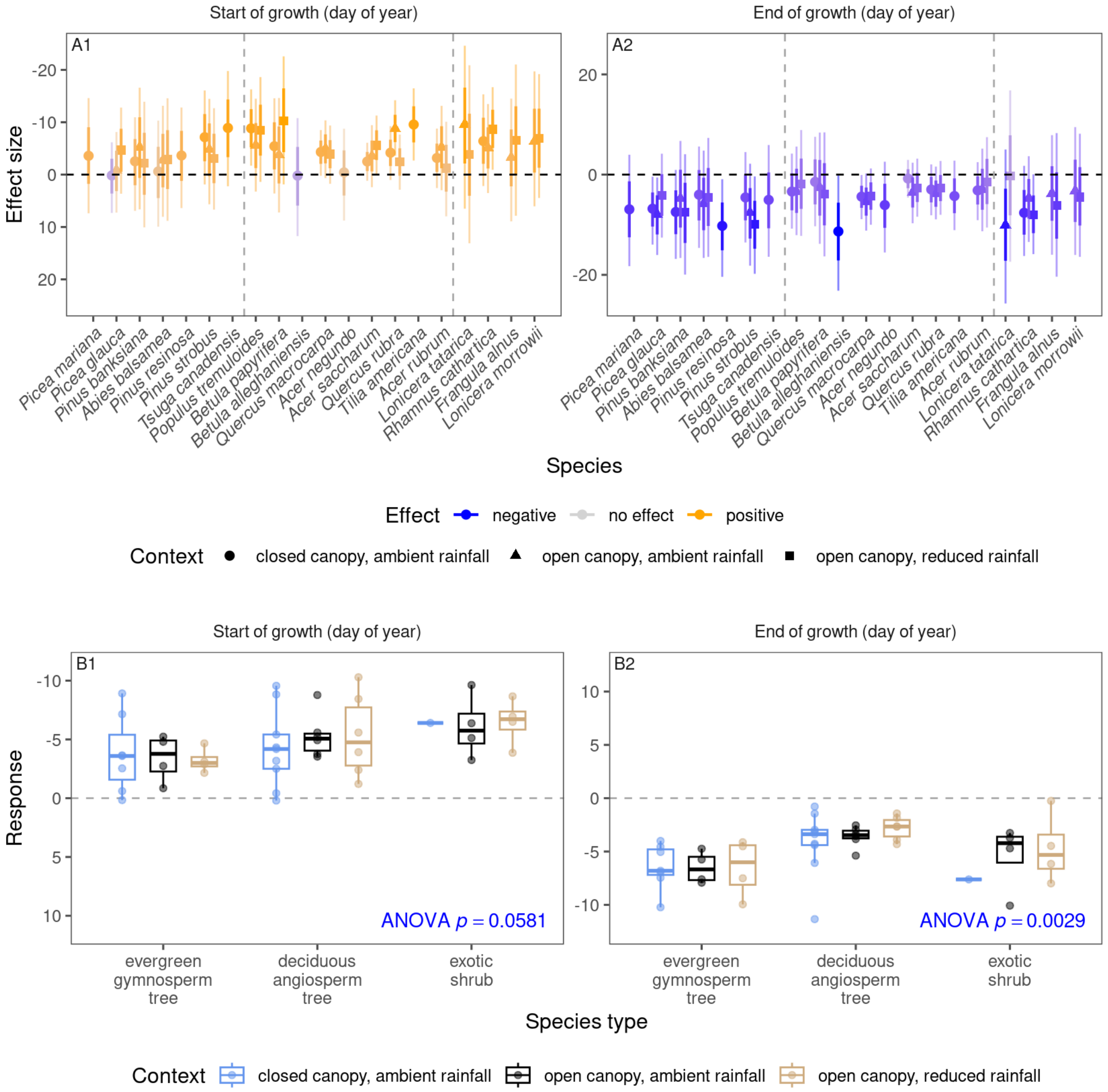


**Figure S12.** Effects of 3 °C warming on the start of growth (time of 5% logarithmic growth; day of year; with effects shown on a reversed scale to highlight advancement) and end of growth (time of 95% logarithmic growth; day of year) under various environmental contexts. Effect sizes were inferred with Bayesian hierarchical models. **(A)** Points show medians of effect sizes, with thick and thin error bars showing 68% and 95% credible intervals of effect sizes, respectively. Colors indicate the direction and magnitude of warming effects on growth, with positive effects associated with earlier start and later end of growth. Order of species follows that in Fig. 2A. **(B)** Species' height growth responses to warming treatment are summarized into three species types across environmental contexts. Boxplots show the median (line), interquartile range (box), and 95% quantile range (whiskers) of species' posterior median estimates. *p*-values are from ANOVA for the predictor of species type, derived from linear mixed-effects models.

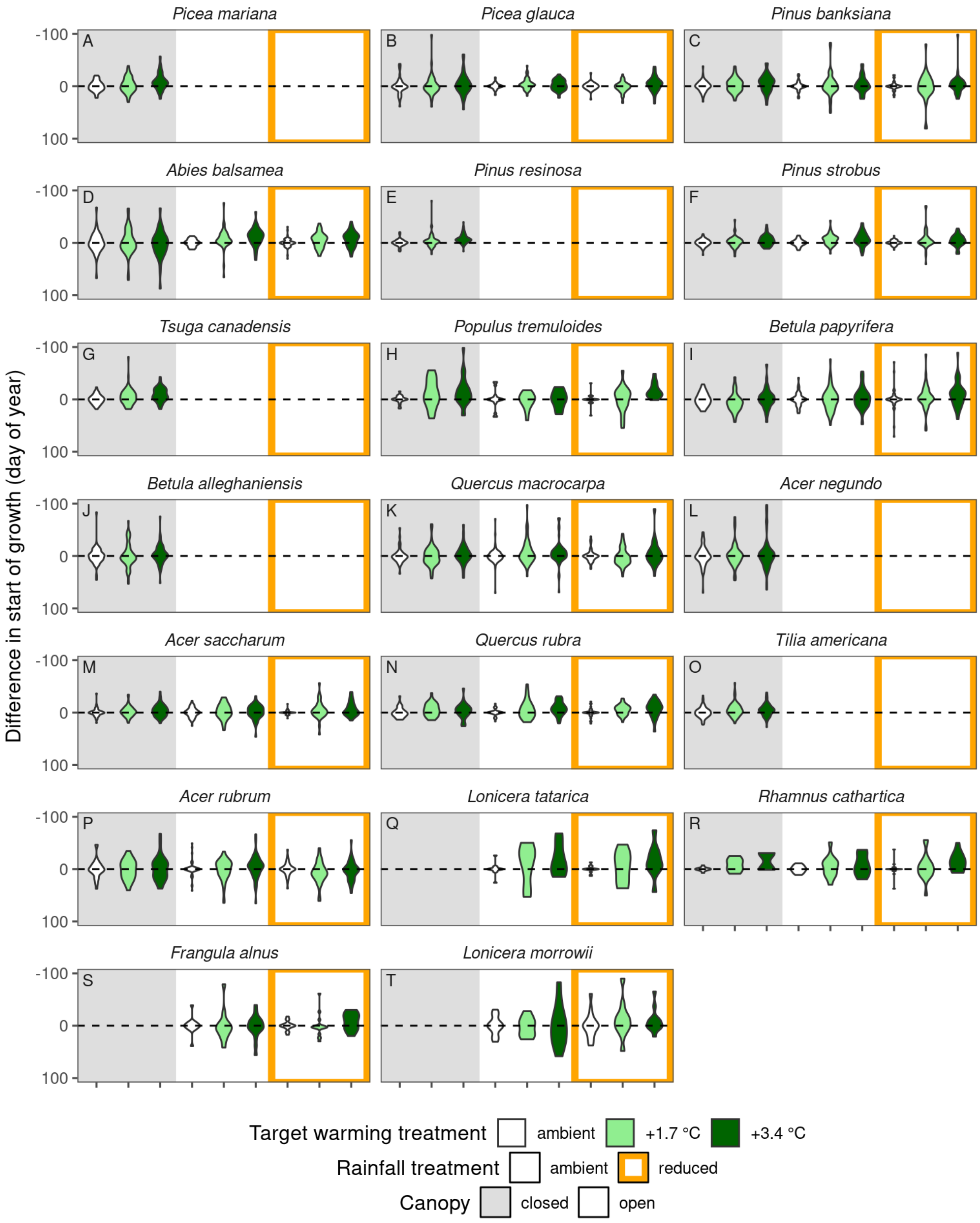


**Figure S13.** Effects of warming on the start of growth (time of 5% logarithmic growth; day of year; effect sizes shown on a reversed scale to highlight advancement), estimated with nonlinear least squares (NLS). Violins show the deviations from the seedlings receiving ambient warming treatment in the same experimental block.

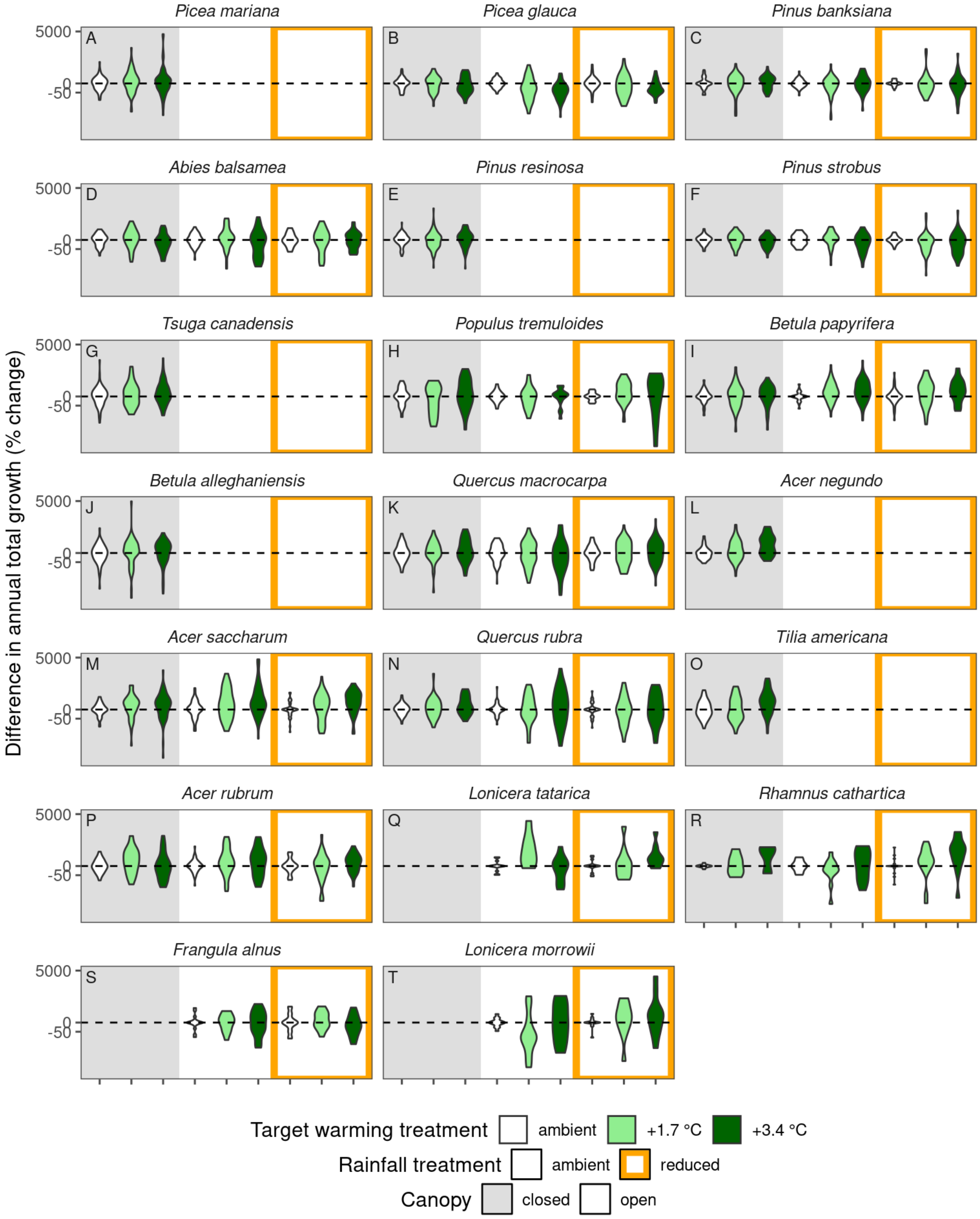


**Figure S14.** Effects of warming on annual total growth (mm; differences on a log scale shown as equivalent percentage changes), estimated with nonlinear least squares (NLS). Violins show the deviations from the seedlings receiving ambient warming treatment in the same experimental block.

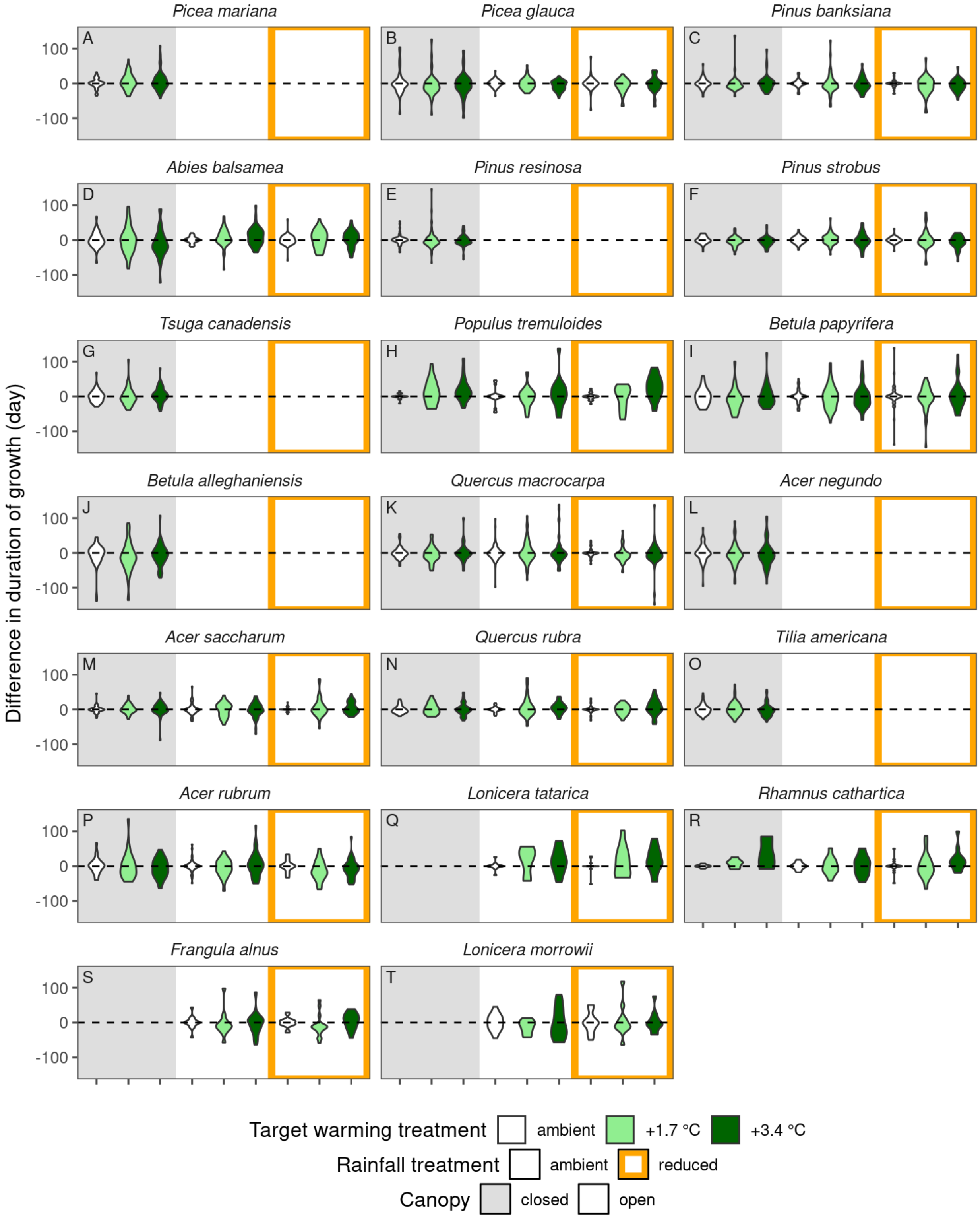


**Figure S15.** Effects of warming on duration of growth (duration from 5% to 95% logarithmic growth, day), estimated with nonlinear least squares (NLS). Violins show the deviations from the seedlings receiving ambient warming treatment in the same experimental block.

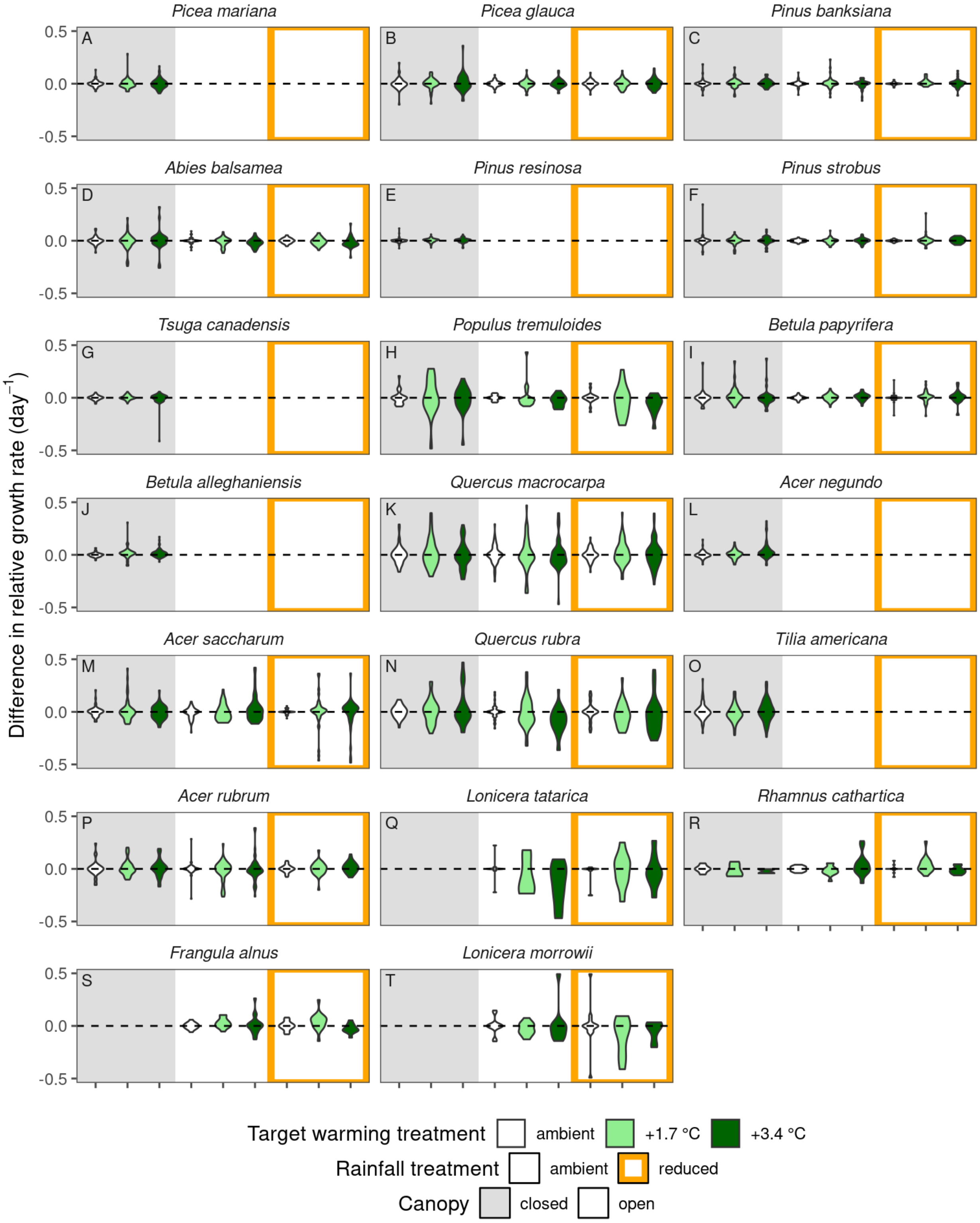


**Figure S16.** Effects of warming on maximum relative growth rate ($day^{-1}$), estimated with nonlinear least squares (NLS). Violins show the deviations from the seedlings receiving ambient warming treatment in the same experimental block.

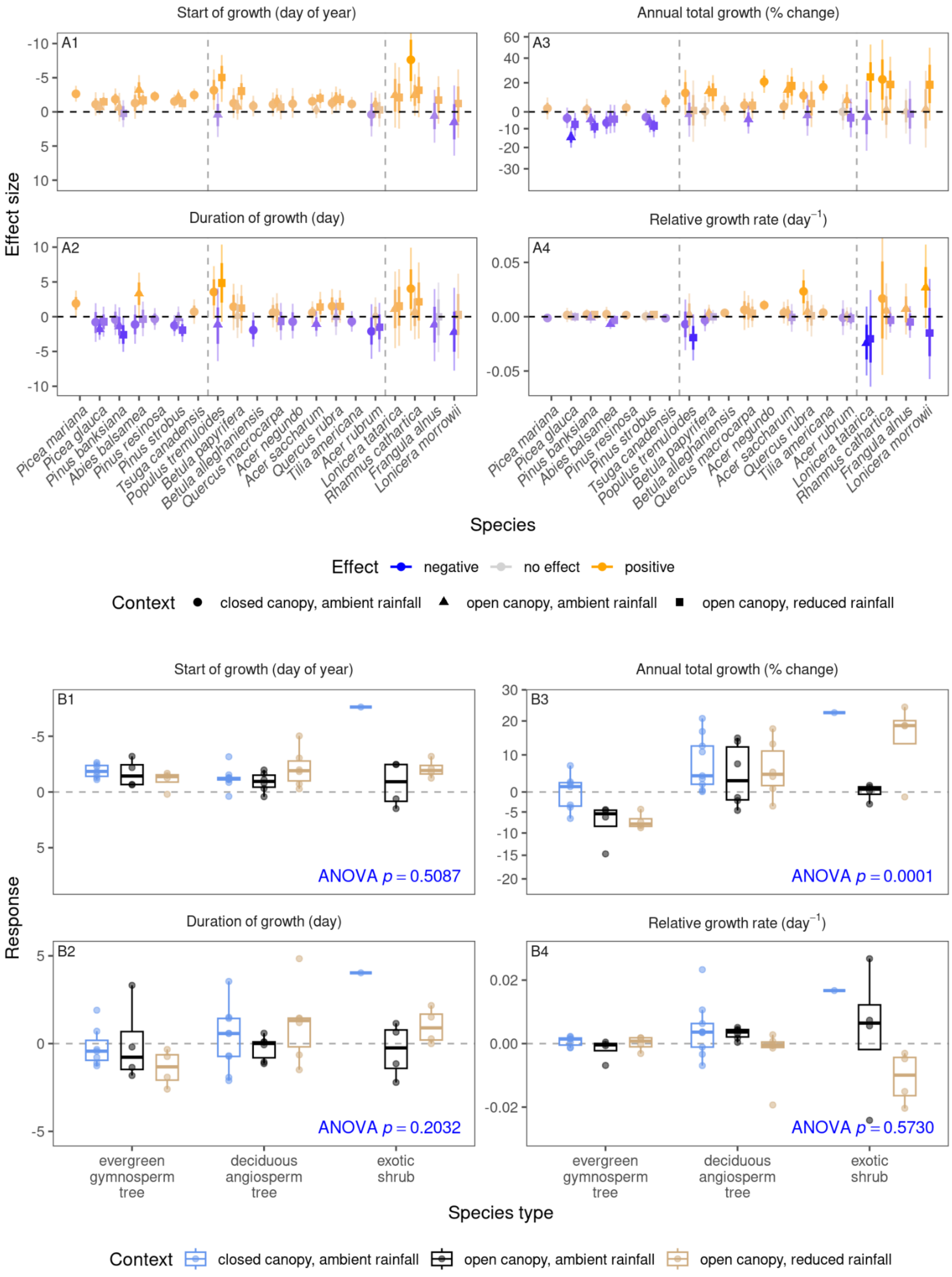

Start of growth (day of year)
Annual total growth (% change)
Duration of growth (day)
Relative growth rate (day$^{-1}$)
A1
A2
A3
A4
Effect size
Picea mariana
Picea glauca
Pinus banksiana
Abies balsamea
Pinus resinosa
Pinus strobus
Tsuga canadensis
Populus tremuloides
Betula papyrifera
Betula alleghaniensis
Quercus macrocarpa
Acer negundo
Acer saccharum
Quercus rubra
Tilia americana
Acer rubrum
Lonicera tatarica
Rhamnus cathartica
Frangula alnus
Lonicera morrowii
Species
Effect negative no effect positive
Context closed canopy, ambient rainfall open canopy, ambient rainfall open canopy, reduced rainfall
B1
B2
B3
B4
Response
ANOVA $p = 0.5087$
ANOVA $p = 0.0001$
ANOVA $p = 0.2032$
ANOVA $p = 0.5730$
evergreen gymnosperm tree
deciduous angiosperm tree
exotic shrub
Species type
Context closed canopy, ambient rainfall open canopy, ambient rainfall open canopy, reduced rainfall

**Figure S17.** Effects of 3 °C warming treatment on the start of growth (time of 5% logarithmic growth; day of year; effect sizes shown on a reversed scale to highlight advancement), annual total growth (mm; effect sizes on a log scale shown as equivalent percentage changes), duration of growth (duration from 5% to 95% logarithmic growth; day), and maximum relative growth rate ($day^{-1}$) under various environmental contexts. Effect sizes were inferred with a NLS-LME two-stage approach. **(A)** Points show medians of effect sizes, with segments showing 95% credible intervals of effect sizes. Colors indicate the direction and magnitude of warming effects on growth, with positive effects associated with earlier growth, higher annual total growth, longer duration, and faster speed. Order of species follows that in Fig. 2A. **(B)** Species' height growth responses to warming treatment summarized into three species types and across environmental contexts. Boxplots show the median (line), interquartile range (box), and 95% quantile range (whiskers) of species' posterior median estimates. *p*-values are from ANOVA for the predictor of species type, derived from linear mixed-effects models.

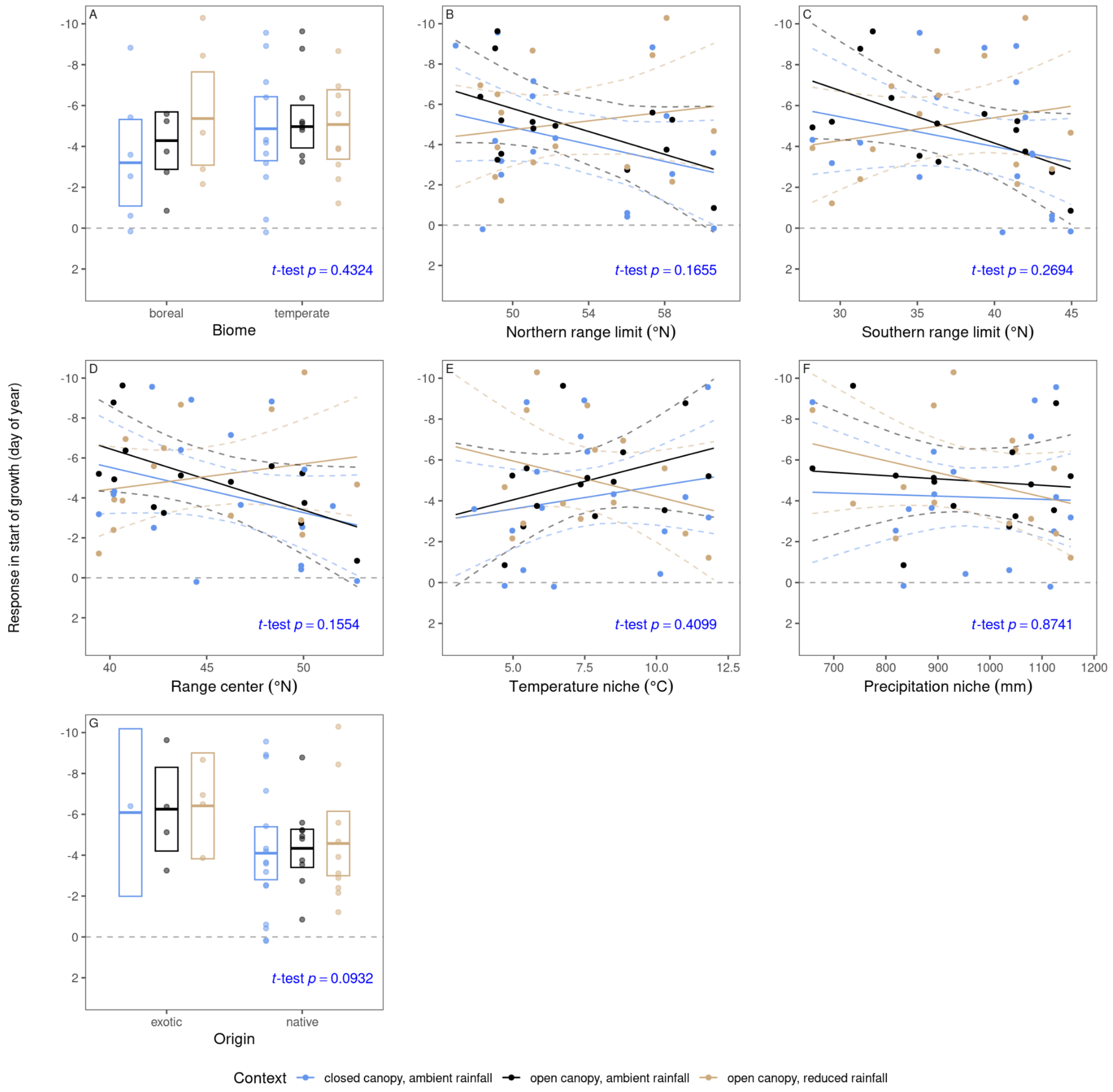


**Figure S18.** Relationship between responses in the start of growth (time of 5% logarithmic growth; day of year; effect sizes shown on a reversed scale to highlight advancement) per 3 °C warming treatment and species' distributions. *p*-values were derived from linear mixed-effects models (*t*-test). Median and 95% credible intervals of predictions are shown with the middle and end segments of boxes for discrete predictors and with the regression lines and ribbons for continuous predictors.

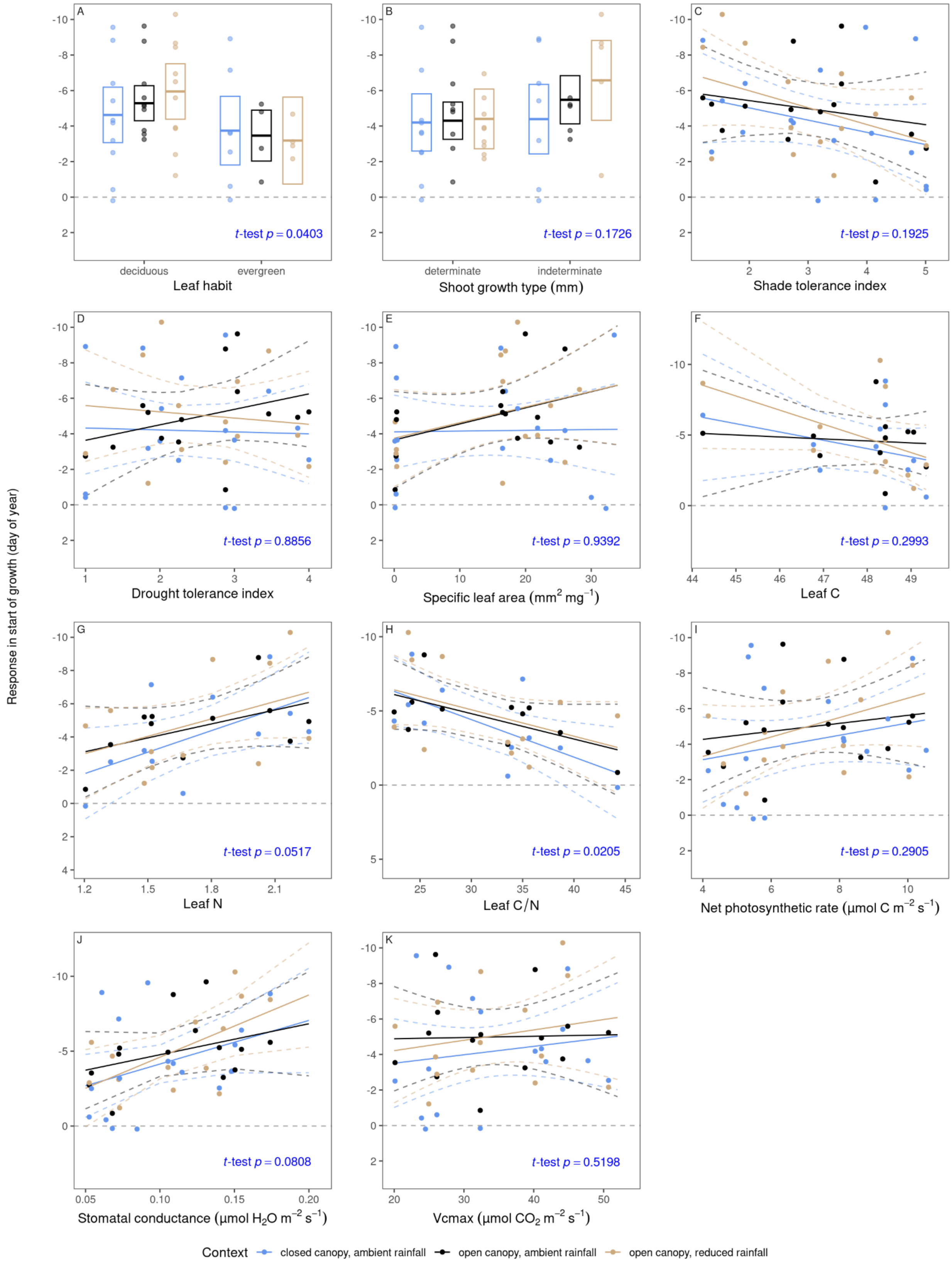

Response in start of growth (day of year)
A
t-test p = 0.0403
deciduous
evergreen
Leaf habit
B
t-test p = 0.1726
determinate
indeterminate
Shoot growth type (mm)
C
t-test p = 0.1925
Shade tolerance index
D
t-test p = 0.8856
Drought tolerance index
E
t-test p = 0.9392
Specific leaf area (mm2 mg−1)
F
t-test p = 0.2993
Leaf C
G
t-test p = 0.0517
Leaf N
H
t-test p = 0.0205
Leaf C/N
I
t-test p = 0.2905
Net photosynthetic rate (μmol C m−2 s−1)
J
t-test p = 0.0808
Stomatal conductance (μmol H2O m−2 s−1)
K
t-test p = 0.5198
Vcmax (μmol CO2 m−2 s−1)
Context
closed canopy, ambient rainfall
open canopy, ambient rainfall
open canopy, reduced rainfall

**Figure S19.** Relationship between responses in the start of growth (time of 5% logarithmic growth; day of year; effect sizes shown on a reversed scale to highlight advancement) per 3 °C warming treatment and species' functional traits. *p*-values were derived from linear mixed-effects models (*t*-test). Median and 95% credible intervals of predictions are shown with the middle and end segments of boxes for discrete predictors and with the regression lines and ribbons for continuous predictors.

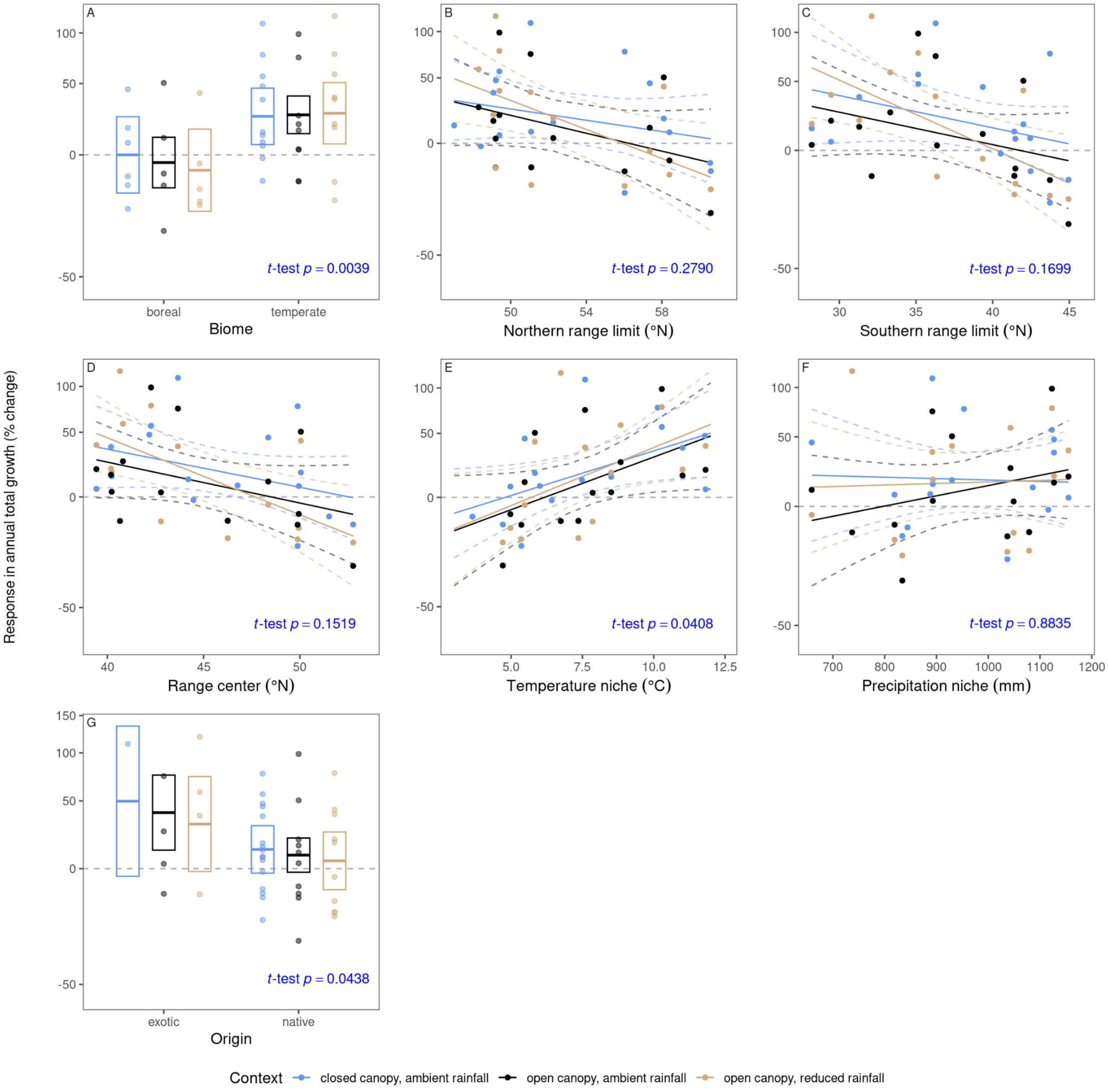


**Figure S20.** Relationship between responses in annual total growth (mm; effect sizes on a log scale shown as equivalent percentage changes) per 3 °C warming treatment and species' distributions. *p*-values were derived from linear mixed-effects models (*t*-test). Median and 95% credible intervals of predictions are shown with the middle and end segments of boxes for discrete predictors and with the regression lines and ribbons for continuous predictors.

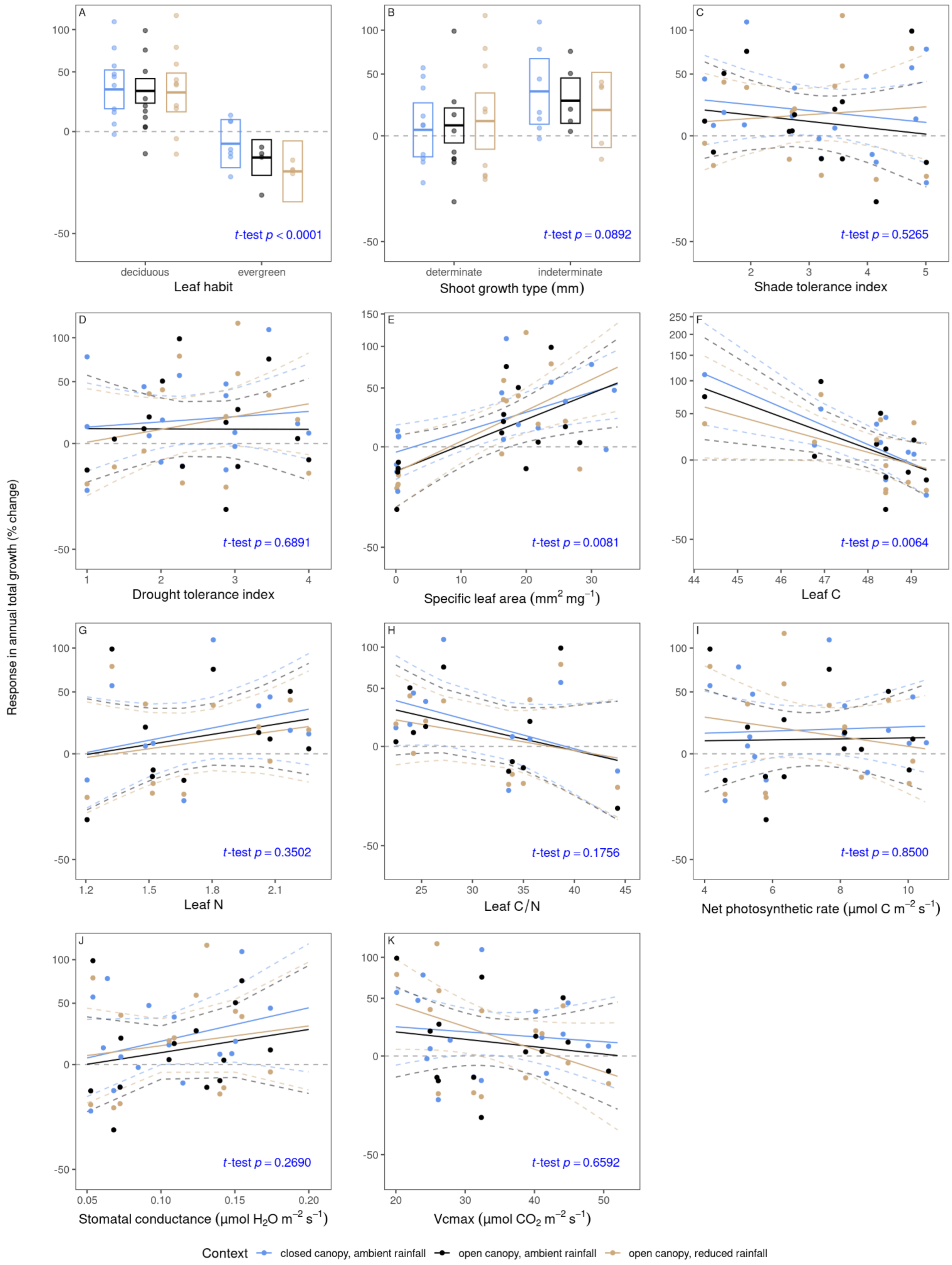

Response in annual total growth (% change)
A
t-test p < 0.0001
deciduous
evergreen
Leaf habit
B
t-test p = 0.0892
determinate
indeterminate
Shoot growth type (mm)
C
t-test p = 0.5265
Shade tolerance index
D
t-test p = 0.6891
Drought tolerance index
E
t-test p = 0.0081
Specific leaf area (mm2 mg−1)
F
t-test p = 0.0064
Leaf C
G
t-test p = 0.3502
Leaf N
H
t-test p = 0.1756
Leaf C/N
I
t-test p = 0.8500
Net photosynthetic rate (μmol C m−2 s−1)
J
t-test p = 0.2690
Stomatal conductance (μmol H2O m−2 s−1)
K
t-test p = 0.6592
Vcmax (μmol CO2 m−2 s−1)
Context
closed canopy, ambient rainfall
open canopy, ambient rainfall
open canopy, reduced rainfall

**Figure S21.** Relationship between responses in annual total growth (mm; effect sizes on a log scale shown as equivalent percentage changes) per 3 °C warming treatment and species' functional traits. *p*-values were derived from linear mixed-effects models (*t*-test). Median and 95% credible intervals of predictions are shown with the middle and end segments of boxes for discrete predictors and with the regression lines and ribbons for continuous predictors.

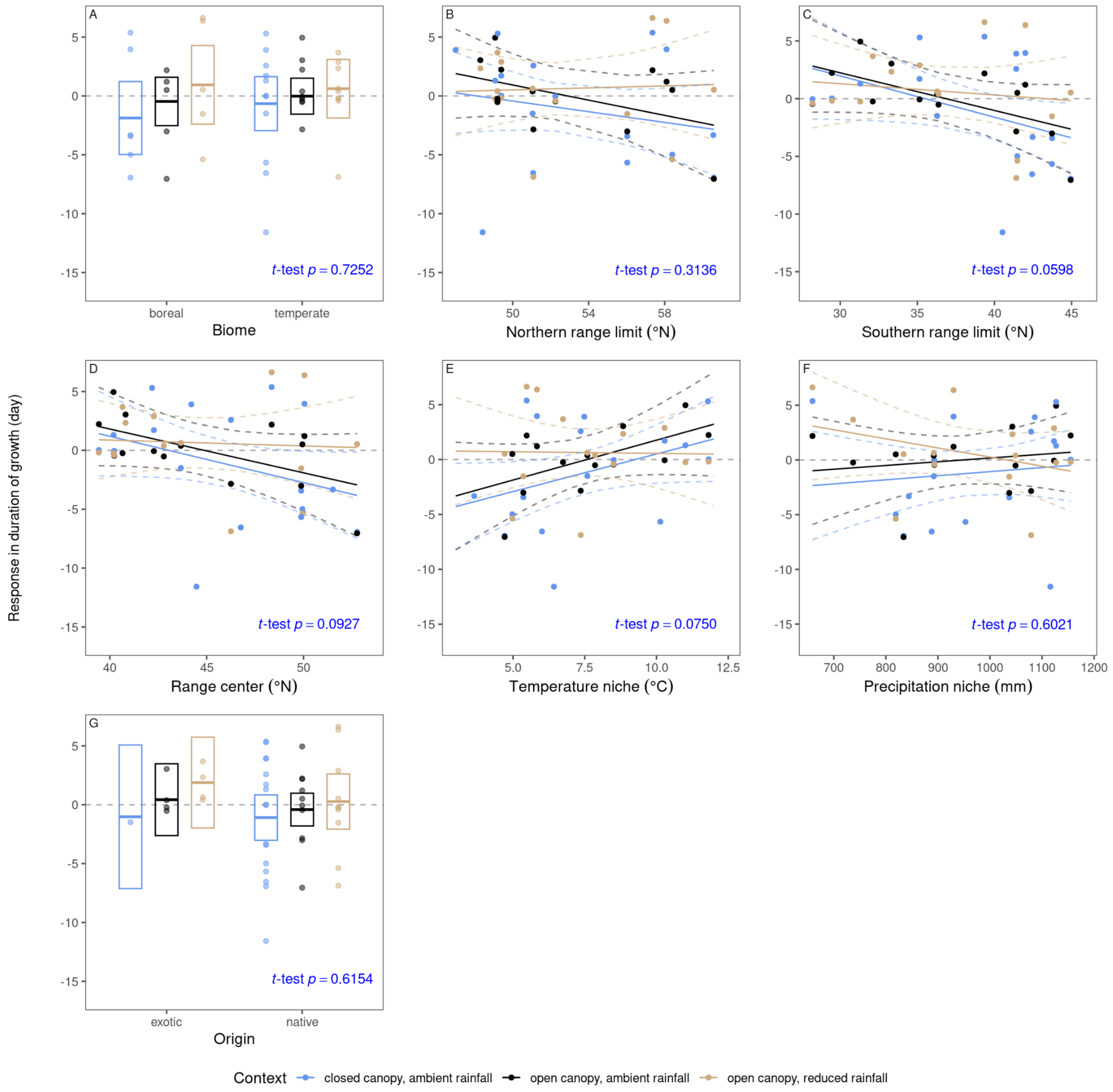


**Figure S22.** Relationship between responses in duration of growth (duration from 5% to 95% logarithmic growth; day) per 3 °C warming treatment and species' distributions. *p*-values were derived from linear mixed-effects models (*t*-test). Median and 95% credible intervals of predictions are shown with the middle and end segments of boxes for discrete predictors and with the regression lines and ribbons for continuous predictors.

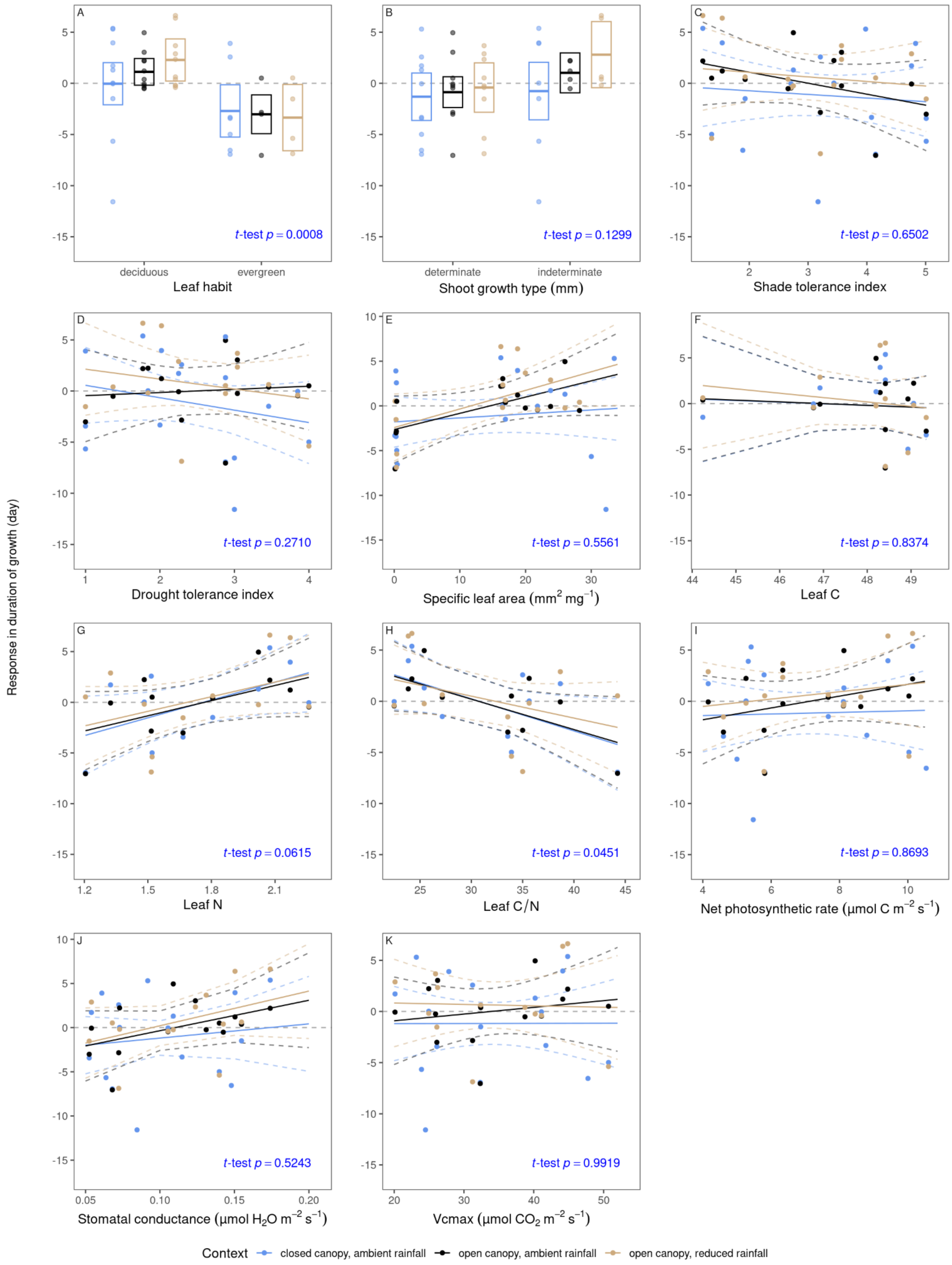
A
B
C
D
E
F
G
H
I
J
K
t-test p = 0.0008
t-test p = 0.1299
t-test p = 0.6502
t-test p = 0.2710
t-test p = 0.5561
t-test p = 0.8374
t-test p = 0.0615
t-test p = 0.0451
t-test p = 0.8693
t-test p = 0.5243
t-test p = 0.9919
deciduous
evergreen
Leaf habit
determinate
indeterminate
Shoot growth type (mm)
Shade tolerance index
Drought tolerance index
Specific leaf area (mm² mg⁻¹)
Leaf C
Leaf N
Leaf C/N
Net photosynthetic rate (μmol C m⁻² s⁻¹)
Stomatal conductance (μmol H₂O m⁻² s⁻¹)
Vcmax (μmol CO₂ m⁻² s⁻¹)
Response in duration of growth (day)
Context
closed canopy, ambient rainfall
open canopy, ambient rainfall
open canopy, reduced rainfall

**Figure S23.** Relationship between responses in duration of growth (duration from 5% to 95% logarithmic growth; day) per 3 °C warming treatment and species' functional traits. *p*-values were derived from linear mixed-effects models (*t*-test). Median and 95% credible intervals of predictions are shown with the middle and end segments of boxes for discrete predictors and with the regression lines and ribbons for continuous predictors.

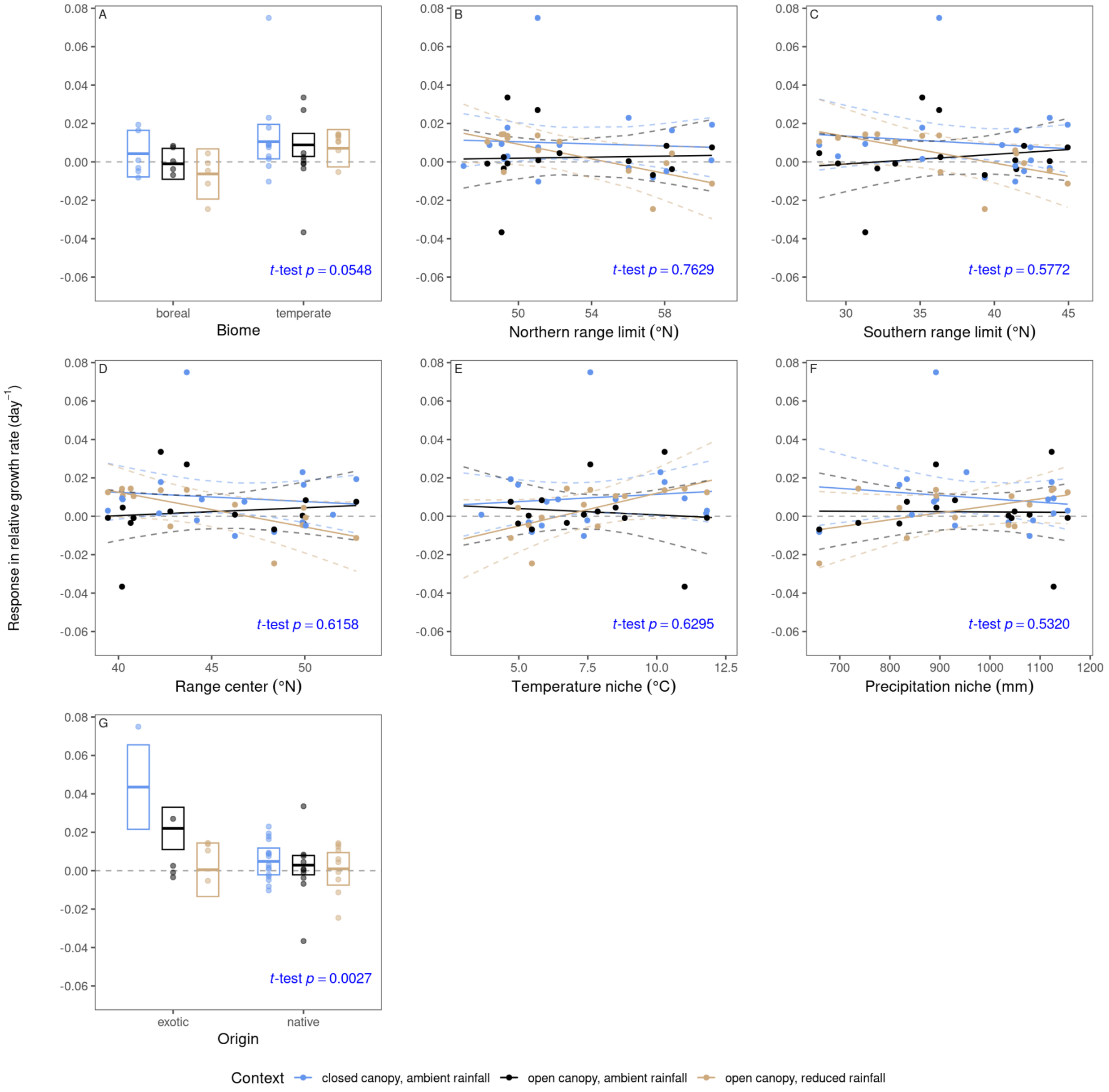


**Figure S24.** Relationship between responses in maximum relative growth rate (day$^{-1}$) per 3 °C warming treatment and species' distributions. *p*-values were derived from linear mixed-effects models (*t*-test). Median and 95% credible intervals of predictions are shown with the middle and end segments of boxes for discrete predictors and with the regression lines and ribbons for continuous predictors.

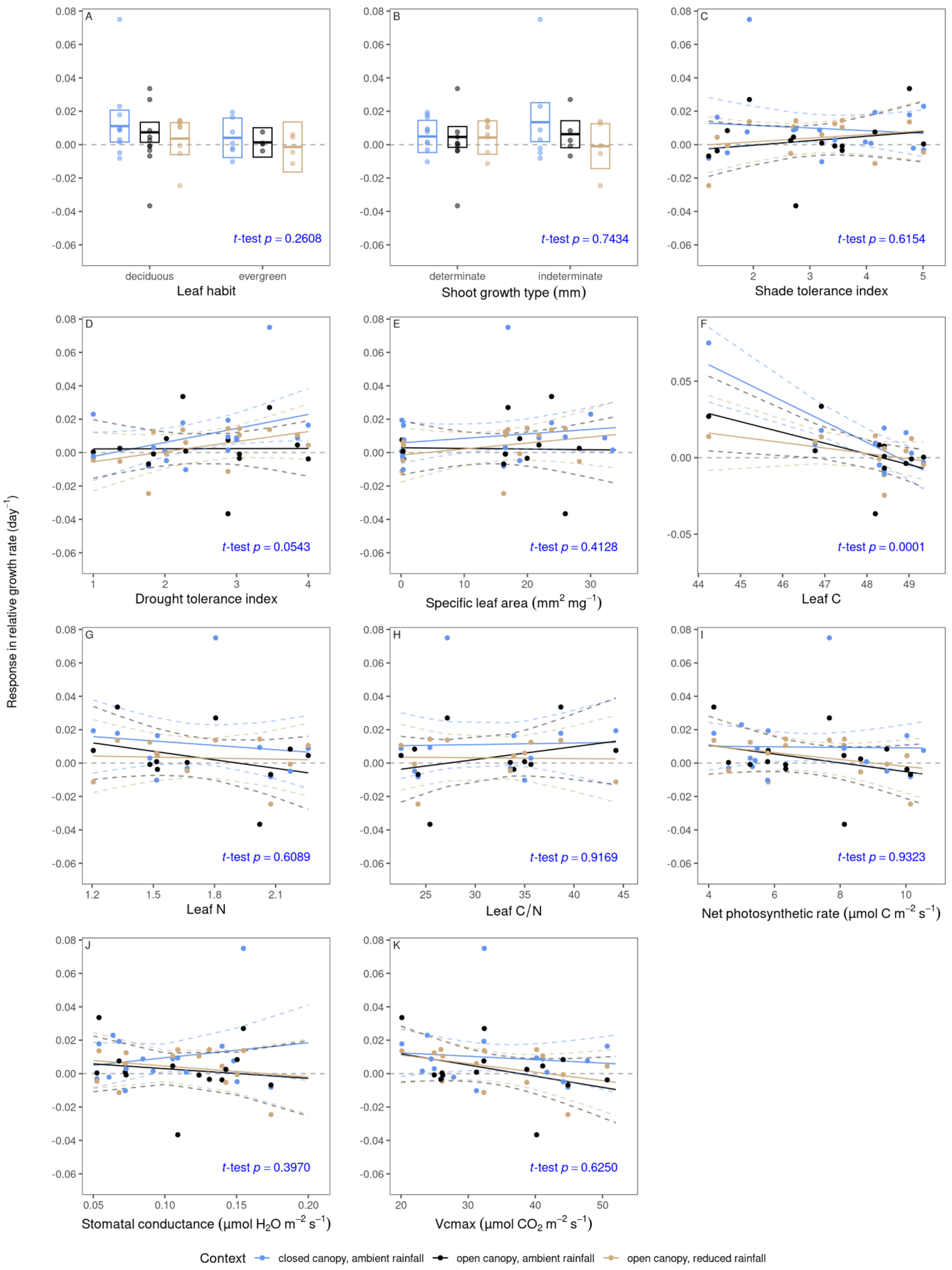

Response in relative growth rate (day⁻¹)
A
t-test p = 0.2608
deciduous
evergreen
Leaf habit
B
t-test p = 0.7434
determinate
indeterminate
Shoot growth type (mm)
C
t-test p = 0.6154
Shade tolerance index
D
t-test p = 0.0543
Drought tolerance index
E
t-test p = 0.4128
Specific leaf area (mm² mg⁻¹)
F
t-test p = 0.0001
Leaf C
G
t-test p = 0.6089
Leaf N
H
t-test p = 0.9169
Leaf C/N
I
t-test p = 0.9323
Net photosynthetic rate (μmol C m⁻² s⁻¹)
J
t-test p = 0.3970
Stomatal conductance (μmol H₂O m⁻² s⁻¹)
K
t-test p = 0.6250
Vcmax (μmol CO₂ m⁻² s⁻¹)
Context
closed canopy, ambient rainfall
open canopy, ambient rainfall
open canopy, reduced rainfall

**Figure S25.** Relationship between responses in maximum relative growth rate ($day^{-1}$) per 3 °C warming treatment and species' functional traits. *p*-values were derived from linear mixed-effects models (*t*-test). Median and 95% credible intervals of predictions are shown with the middle and end segments of boxes for discrete predictors and with the regression lines and ribbons for continuous predictors.

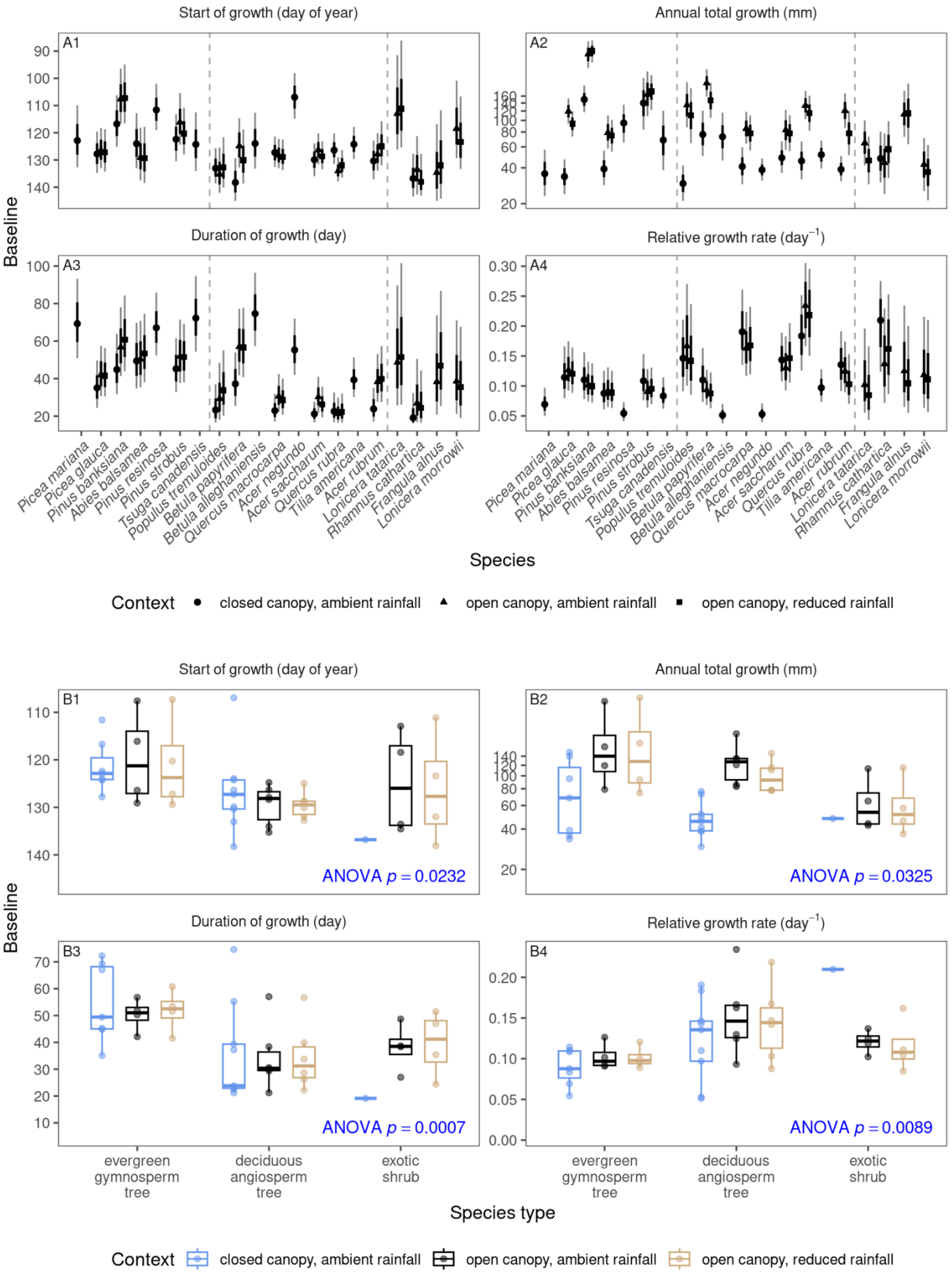

Start of growth (day of year)
Annual total growth (mm)
Duration of growth (day)
Relative growth rate (day$^{-1}$)
A1
A2
A3
A4
Baseline
Picea mariana
Picea glauca
Pinus banksiana
Abies balsamea
Pinus resinosa
Pinus strobus
Tsuga canadensis
Populus tremuloides
Betula papyrifera
Betula alleghaniensis
Quercus macrocarpa
Acer negundo
Acer saccharum
Quercus rubra
Tilia americana
Acer rubrum
Lonicera tatarica
Rhamnus cathartica
Frangula alnus
Lonicera morrowii
Species
Context
closed canopy, ambient rainfall
open canopy, ambient rainfall
open canopy, reduced rainfall
Start of growth (day of year)
Annual total growth (mm)
Duration of growth (day)
Relative growth rate (day$^{-1}$)
B1
B2
B3
B4
ANOVA $p = 0.0232$
ANOVA $p = 0.0325$
ANOVA $p = 0.0007$
ANOVA $p = 0.0089$
Baseline
evergreen gymnosperm tree
deciduous angiosperm tree
exotic shrub
Species type
Context
closed canopy, ambient rainfall
open canopy, ambient rainfall
open canopy, reduced rainfall

**Figure S26.** Baselines of the start of growth (time of 5% logarithmic growth; day of year), annual total growth (mm; visualized on a log scale), duration of growth (duration from 5% to 95% logarithmic growth; day), and maximum relative growth rate ($day^{-1}$) in ambient temperature under various environmental contexts. Baselines were inferred with Bayesian hierarchical models. **(A)** Points show medians of baselines, with thick and thin error bars showing 68% and 95% credible intervals of baselines, respectively. **(B)** Species' baseline parameters are summarized into three species types across environmental contexts. Boxplots show the median (line), interquartile range (box), and 95% quantile range (whiskers) of species' posterior median estimates. *p*-values are from ANOVA for the predictor of species type, derived from linear mixed-effects models. Order of species follows that in Fig. 2A.

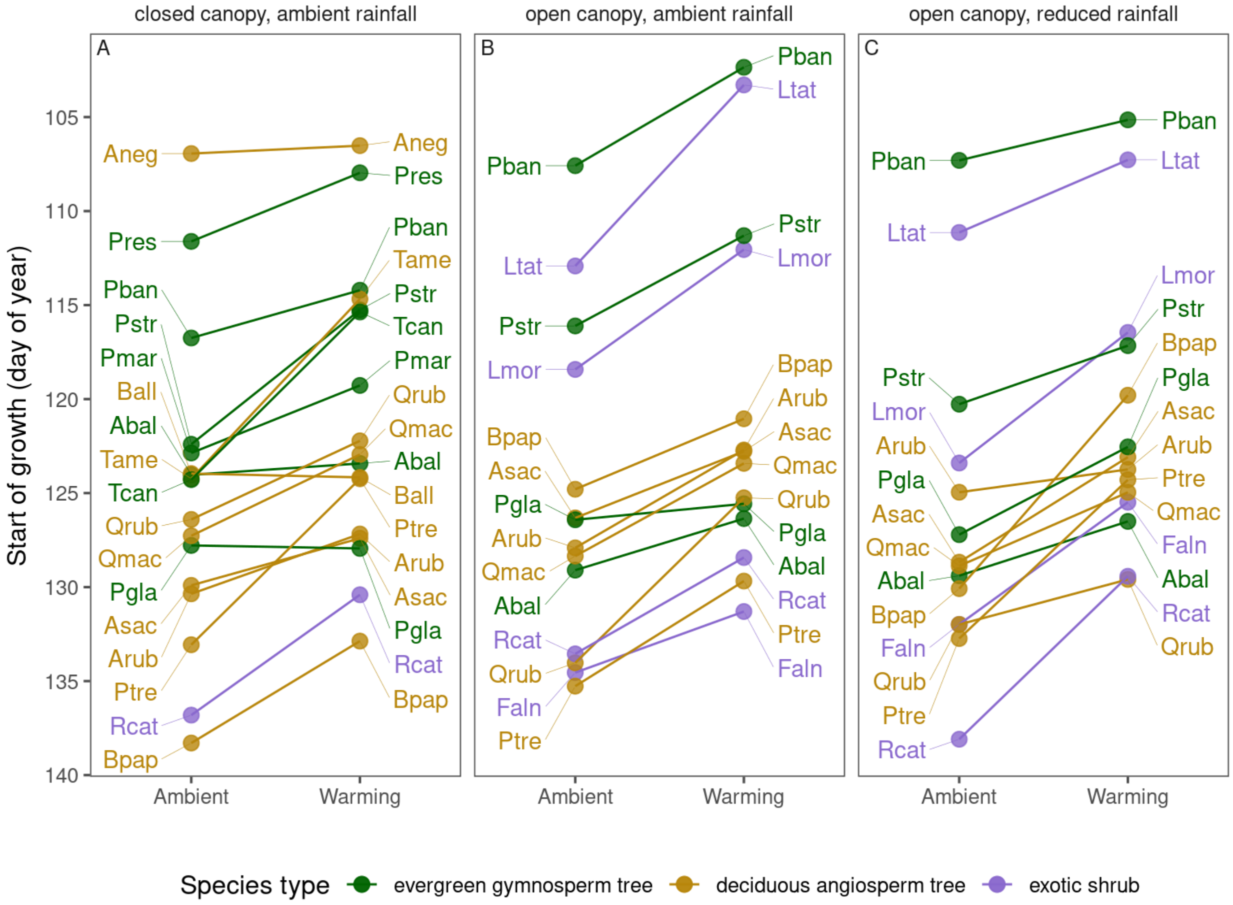


**Figure S27.** Model-predicted change in phenological rank (order of start of height growth among species) between ambient temperature and 3 °C warming across environmental contexts. Species acronyms are provided in Table S1.

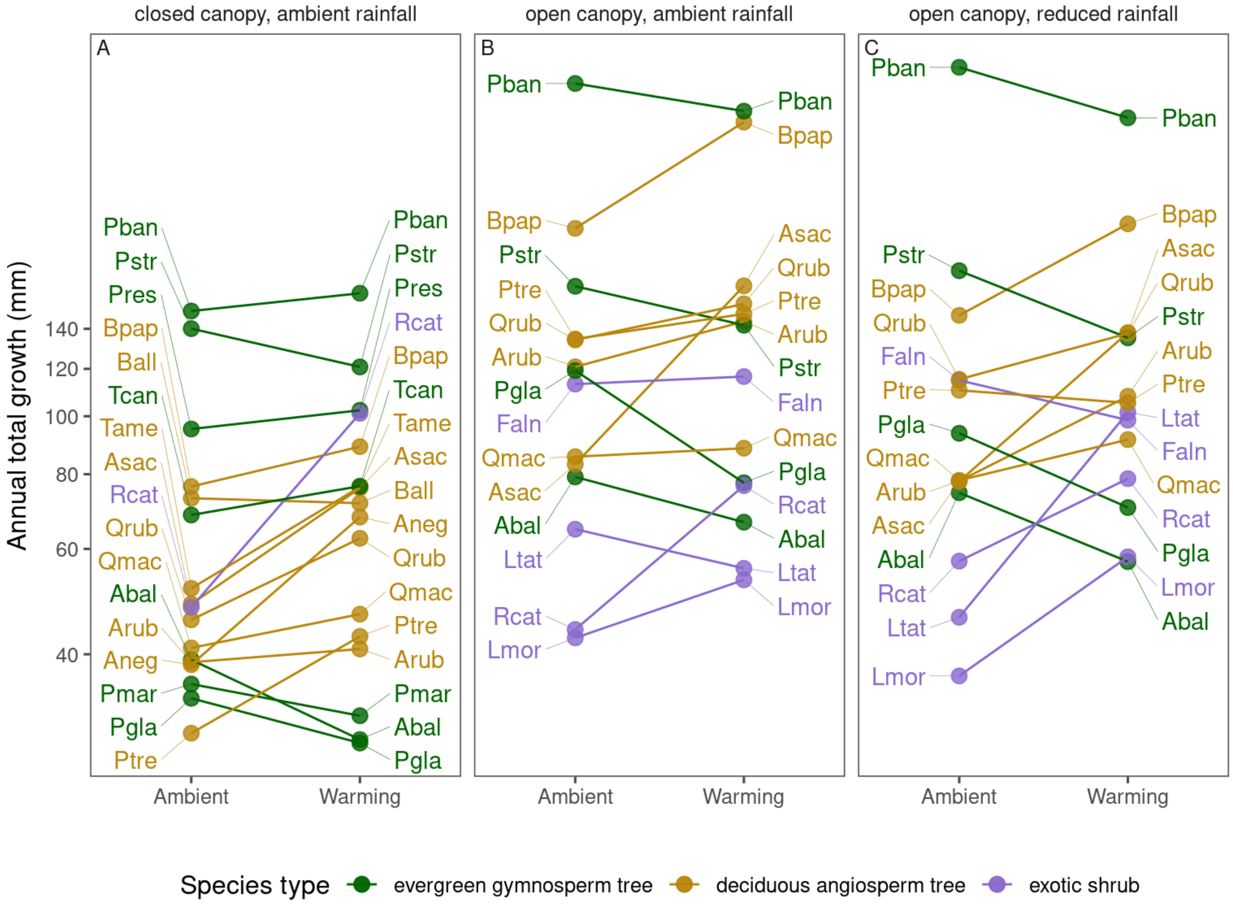


**Figure S28.** Model-predicted change in growth rank (order of annual total growth among species) between ambient temperature and 3 °C warming across environmental contexts. Species acronyms are provided in Table S1.

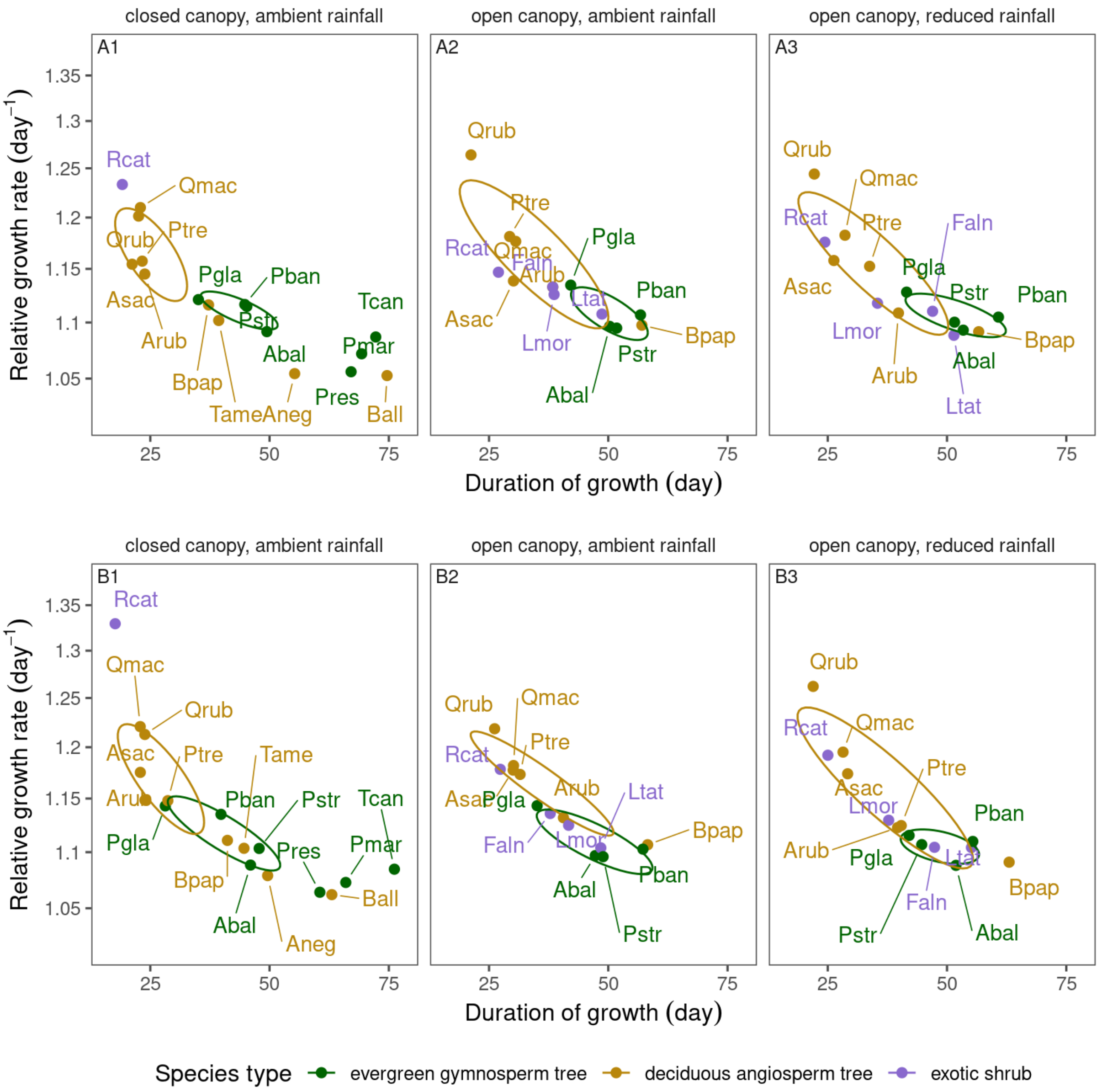


**Figure S29.** Model-predicted changes in duration of growth (day) and maximum relative growth rate (day$^{-1}$) under **(A)** ambient temperature and **(B)** 3 °C warming treatments across environmental contexts. Ellipses represent 50% concentration regions for each species type, assuming multivariate normal distributions. Species acronyms are provided in Table S1.

**Table S1.** Temperate and boreal tree species planted in the B4WarmED experiment. We highlighted 11 species that received the complete factorial design (Fig. 1), while the other species were planted in a subset of treatment combinations.

| Scientific name | Acronym | Common name | Biome | Leaf habit | Shoot growth type |
|---|---|---|---|---|---|
| *Acer negundo* | Aneg | box elder | temperate | deciduous | indeterminate |
| *Acer rubrum* | Arub | red maple | temperate | deciduous | indeterminate |
| *Acer saccharum* | Asac | sugar maple | temperate | deciduous | determinate |
| *Betula alleghaniensis* | Ball | yellow birch | temperate | deciduous | indeterminate |
| *Betula papyrifera* | Bpap | paper birch | boreal | deciduous | indeterminate |
| *Frangula alnus* | Faln | glossy buckthorn | exotic | deciduous | indeterminate |
| *Lonicera morrowii* | Lmor | Morrow's honeysuckle | exotic | deciduous | determinate |
| *Lonicera tatarica* | Ltat | tatarian honeysuckle | exotic | deciduous | determinate |
| *Populus tremuloides* | Ptre | trembling aspen | boreal | deciduous | indeterminate |
| *Quercus macrocarpa* | Qmac | bur oak | temperate | deciduous | determinate |
| *Quercus rubra* | Qrub | red oak | temperate | deciduous | determinate |
| *Rhamnus cathartica* | Rcat | common buckthorn | exotic | deciduous | indeterminate |
| *Tilia americana* | Tame | american basswood | temperate | deciduous | determinate |
| *Abies balsamea* | Abal | balsam fir | boreal | evergreen | determinate |
| *Picea glauca* | Pgla | white spruce | boreal | evergreen | determinate |
| *Picea mariana* | Pmar | black spruce | boreal | evergreen | determinate |
| *Pinus banksiana* | Pban | jack pine | boreal | evergreen | determinate |
| *Pinus resinosa* | Pres | red pine | temperate | evergreen | determinate |
| *Pinus strobus* | Pstr | eastern white pine | temperate | evergreen | determinate |
| *Tsuga canadensis* | Tcan | canadian hemlock | temperate | evergreen | indeterminate |

**Table S2.** Treatment effects on environmental variables, including aboveground air temperature (°C) and soil moisture ($mm^3$ water $mm^{-3}$ soil). For each plot and year, we averaged environmental variables during the time period when the experimental heating system was on (generally March to October), spanning the plants' growing season. Treatment effects were estimated by calculating the difference in these environmental variables between each plot and the average from ambient plots in the same experimental block and year. We summarized the temperature increases by environmental contexts. Mean ± standard deviations (among plot-year combinations) in differences are reported.

| warming treatment target | watering treatment | canopy | difference in aboveground air temperature | difference in soil moisture |
|---|---|---|---|---|
| +1.7 °C | ambient | closed | 1.63 ± 0.434 | −0.000848 ± 0.0258 |
| +3.4 °C | ambient | closed | 3.47 ± 0.153 | −0.0223 ± 0.0217 |
| +1.7 °C | ambient | open | 1.64 ± 0.179 | −0.0260 ± 0.0180 |
| +3.4 °C | ambient | open | 3.23 ± 0.176 | −0.0445 ± 0.0189 |
| +1.7 °C | reduced | open | 1.61 ± 0.199 | −0.0194 ± 0.0150 |
| +3.4 °C | reduced | open | 3.22 ± 0.196 | −0.0463 ± 0.0301 |

**Table S3.** Effects of 3 °C warming treatment on the **(A)** start of growth (time of 5% logarithmic growth; day of year), **(B)** annual total growth (mm; effect sizes on a log scale shown as equivalent percentage changes), **(C)** duration of growth (duration from 5% to 95% logarithmic growth; day), and **(D)** maximum relative growth rate ($day^{-1}$) under various environmental contexts. Medians and 95% credible intervals of effect sizes are reported. Effect sizes that are statistically significant ($p \leq 0.05$) are bolded. Order of species follows that in Fig. 2A.

(A) start of growth

| species | closed canopy, ambient rainfall | open canopy, ambient rainfall | open canopy, reduced rainfall |
|---|---|---|---|
| *Picea mariana* | −3.59<br>(−14.6, 7.39) | | |
| *Picea glauca* | 0.160<br>(−6.16, 7.31) | −0.851<br>(−8.13, 7.25) | −4.67<br>(−12.8, 3.65) |
| *Pinus banksiana* | −2.54<br>(−11.0, 6.91) | −5.23<br>(−16.6, 6.63) | −2.16<br>(−13.9, 10.1) |
| *Abies balsamea* | −0.606<br>(−10.3, 9.94) | −2.74<br>(−13.1, 8.05) | −2.89<br>(−14.6, 8.81) |
| *Pinus resinosa* | −3.65<br>(−12.8, 6.38) | | |
| *Pinus strobus* | −7.15<br>(−16.1, 1.88) | −4.80<br>(−14.5, 5.66) | −3.11<br>(−12.7, 6.68) |
| *Tsuga canadensis* | −8.91<br>(−19.8, 2.22) | | |
| *Populus tremuloides* | **−8.83**<br>**(−16.3, −1.71)** | −5.59<br>(−14.5, 3.23) | −8.44<br>(−18.6, 1.38) |
| *Betula papyrifera* | −5.42<br>(−14.6, 3.60) | −3.75<br>(−14.6, 7.22) | −10.3<br>(−22.6, 1.87) |
| *Betula alleghaniensis* | 0.202<br>(−10.8, 11.7) | | |
| *Quercus macrocarpa* | −4.32<br>(−8.51, 0.216) | −4.93<br>(−10.4, 0.677) | −3.92<br>(−9.37, 1.74) |
| *Acer negundo* | −0.420<br>(−8.76, 8.77) | | |
| *Acer saccharum* | −2.50<br>(−6.09, 1.17) | −3.54<br>(−9.44, 2.40) | **−5.59**<br>**(−11.3, −0.0441)** |

| | | | |
|---|---|---|---|
| *Quercus rubra* | −4.18 (−9.25, 1.03) | **−8.78 (−14.2, −3.60)** | −2.40 (−7.64, 2.88) |
| *Tilia americana* | **−9.56 (−16.5, −2.70)** | | |
| *Acer rubrum* | −3.18 (−8.79, 2.56) | −5.21 (−13.1, 2.97) | −1.21 (−10.1, 7.96) |
| *Lonicera tatarica* | | −9.63 (−24.6, 6.51) | −3.86 (−20.9, 13.1) |
| *Rhamnus cathartica* | −6.40 (−15.3, 2.18) | −5.12 (−13.5, 3.52) | **−8.67 (−16.7, −0.926)** |
| *Frangula alnus* | | −3.25 (−14.5, 8.92) | −6.50 (−21.0, 7.85) |
| *Lonicera morrowii* | | −6.37 (−19.7, 6.05) | −6.95 (−19.2, 4.46) |

(B) annual total growth

| species | closed canopy, ambient rainfall | open canopy, ambient rainfall | open canopy, reduced rainfall |
|---|---|---|---|
| *Picea mariana* | −11.5 (−28.2, 9.23) | | |
| *Picea glauca* | −15.8 (−31.6, 4.20) | **−35.1 (−49.7, −16.2)** | **−24.9 (−40.4, −4.72)** |
| *Pinus banksiana* | 7.07 (−14.5, 33.7) | −10.1 (−29.7, 14.4) | −17.7 (−34.5, 3.14) |
| *Abies balsamea* | **−26.5 (−41.9, −7.45)** | −16.0 (−38.9, 16.2) | −23.3 (−43.2, 3.78) |
| *Pinus resinosa* | 7.48 (−8.91, 27.2) | | |
| *Pinus strobus* | −13.7 (−29.8, 6.44) | −13.9 (−33.1, 10.8) | **−22.8 (−39.9, −1.21)** |
| *Tsuga canadensis* | 11.7 (−11.1, 40.1) | | |
| *Populus tremuloides* | **45.1 (6.16, 102)** | 10.1 (−25.1, 60.4) | −4.75 (−39.5, 50.5) |
| *Betula papyrifera* | 16.6 (−19.4, 69.2) | **50.5 (20.2, 89.6)** | 42.2 (−0.177, 104) |

| | | | |
|---|---|---|---|
| *Betula alleghaniensis* | −1.89<br>(−28.9, 34.0) | | |
| *Quercus macrocarpa* | 13.9<br>(−6.89, 39.3) | 3.31<br>(−17.7, 29.1) | 17.0<br>(−7.88, 49.6) |
| *Acer negundo* | **76.6**<br>**(42.6, 119)** | | |
| *Acer saccharum* | **56.2**<br>**(26.9, 91.0)** | **98.7**<br>**(46.4, 168)** | **77.3**<br>**(29.5, 142)** |
| *Quercus rubra* | **36.9**<br>**(5.61, 77.4)** | 14.9<br>(−16.2, 58.0) | 19.3<br>(−13.4, 66.2) |
| *Tilia americana* | **47.7**<br>**(16.5, 86.3)** | | |
| *Acer rubrum* | 5.22<br>(−20.4, 39.2) | 19.1<br>(−11.7, 61.3) | 38.5<br>(−6.78, 105) |
| *Lonicera tatarica* | | −14.0<br>(−49.7, 47.5) | **120**<br>**(25.9, 284)** |
| *Rhamnus cathartica* | **111**<br>**(19.8, 268)** | 74.0<br>(−0.477, 208) | 37.4<br>(−20.6, 133) |
| *Frangula alnus* | | 2.95<br>(−38.8, 72.8) | −14.2<br>(−51.0, 50.8) |
| *Lonicera morrowii* | | 25.0<br>(−34.2, 132) | 58.1<br>(−14.7, 202) |

(C) duration of growth

| species | closed canopy, ambient rainfall | open canopy, ambient rainfall | open canopy, reduced rainfall |
|---|---|---|---|
| *Picea mariana* | −3.32<br>(−25.4, 18.0) | | |
| *Picea glauca* | −6.93<br>(−20.9, 5.07) | −7.05<br>(−23.1, 7.14) | 0.530<br>(−16.0, 16.8) |
| *Pinus banksiana* | −4.99<br>(−23.0, 11.2) | 0.508<br>(−23.0, 22.8) | −5.38<br>(−29.7, 17.5) |
| *Abies balsamea* | −3.43<br>(−24.2, 15.3) | −3.01<br>(−24.2, 17.2) | −1.53<br>(−24.8, 21.2) |
| *Pinus resinosa* | −6.55<br>(−26.7, 11.9) | | |

| | | | |
|---|---|---|---|
| *Pinus strobus* | 2.58<br>(−15.3, 20.5) | −2.84<br>(−23.9, 16.4) | −6.87<br>(−26.0, 12.0) |
| *Tsuga canadensis* | 3.90<br>(−18.4, 25.5) | | |
| *Populus tremuloides* | 5.38<br>(−6.87, 18.5) | 2.19<br>(−13.7, 18.6) | 6.63<br>(−12.5, 26.5) |
| *Betula papyrifera* | 3.96<br>(−12.5, 20.9) | 1.21<br>(−20.7, 22.5) | 6.38<br>(−17.3, 30.1) |
| *Betula alleghaniensis* | −11.6<br>(−34.3, 10.2) | | |
| *Quercus macrocarpa* | −0.0377<br>(−8.46, 7.72) | −0.457<br>(−11.6, 10.2) | −0.402<br>(−11.3, 9.95) |
| *Acer negundo* | −5.66<br>(−23.7, 10.8) | | |
| *Acer saccharum* | 1.72<br>(−5.19, 8.48) | −0.0657<br>(−11.7, 11.3) | 2.89<br>(−7.88, 13.8) |
| *Quercus rubra* | 1.30<br>(−8.61, 10.5) | 4.95<br>(−4.47, 15.1) | −0.242<br>(−10.1, 9.41) |
| *Tilia americana* | 5.31<br>(−7.93, 19.0) | | |
| *Acer rubrum* | 0.0230<br>(−10.8, 10.3) | 2.23<br>(−14.0, 17.8) | −0.182<br>(−18.0, 16.8) |
| *Lonicera tatarica* | | −0.241<br>(−30.1, 27.1) | 3.68<br>(−29.2, 35.8) |
| *Rhamnus cathartica* | −1.49<br>(−10.6, 11.0) | 0.385<br>(−15.8, 15.9) | 0.629<br>(−14.0, 15.7) |
| *Frangula alnus* | | −0.519<br>(−24.5, 21.8) | 0.404<br>(−27.3, 28.4) |
| *Lonicera morrowii* | | 3.04<br>(−19.3, 26.8) | 2.34<br>(−18.7, 25.7) |

(D) relative growth rate

| species | closed canopy, ambient rainfall | open canopy, ambient rainfall | open canopy, reduced rainfall |
|---|---|---|---|
| *Picea mariana* | 0.000822<br>(−0.0225, 0.0231) | | |

| | | | |
|---|---|---|---|
| *Picea glauca* | 0.0194 (−0.0301, 0.0722) | 0.00756 (−0.0447, 0.0607) | −0.0113 (−0.0579, 0.0330) |
| *Pinus banksiana* | 0.0164 (−0.0301, 0.0638) | −0.00376 (−0.0439, 0.0369) | 0.00443 (−0.0377, 0.0465) |
| *Abies balsamea* | −0.00311 (−0.0396, 0.0319) | 0.000366 (−0.0399, 0.0404) | −0.00461 (−0.0435, 0.0333) |
| *Pinus resinosa* | 0.00761 (−0.00990, 0.0257) | | |
| *Pinus strobus* | −0.0102 (−0.0533, 0.0294) | 0.000850 (−0.0368, 0.0377) | 0.00603 (−0.0341, 0.0461) |
| *Tsuga canadensis* | −0.00211 (−0.0282, 0.0222) | | |
| *Populus tremuloides* | −0.00815 (−0.0738, 0.0600) | −0.00683 (−0.0982, 0.0756) | −0.0245 (−0.105, 0.0405) |
| *Betula papyrifera* | −0.00484 (−0.0552, 0.0408) | 0.00838 (−0.0292, 0.0441) | −0.000650 (−0.0382, 0.0349) |
| *Betula alleghaniensis* | 0.00880 (−0.0101, 0.0280) | | |
| *Quercus macrocarpa* | 0.00881 (−0.0601, 0.0775) | 0.00455 (−0.0574, 0.0620) | 0.0106 (−0.0553, 0.0754) |
| *Acer negundo* | **0.0230 (0.00201, 0.0447)** | | |
| *Acer saccharum* | 0.0178 (−0.0307, 0.0675) | 0.0336 (−0.0242, 0.0904) | 0.0136 (−0.0477, 0.0775) |
| *Quercus rubra* | 0.00936 (−0.0642, 0.0913) | −0.0366 (−0.120, 0.0488) | 0.0143 (−0.0785, 0.115) |
| *Tilia americana* | 0.00154 (−0.0314, 0.0332) | | |
| *Acer rubrum* | 0.00293 (−0.0556, 0.0644) | −0.000815 (−0.0520, 0.0502) | 0.0125 (−0.0380, 0.0633) |
| *Lonicera tatarica* | | −0.00343 (−0.0686, 0.0550) | 0.0144 (−0.0396, 0.0787) |
| *Rhamnus cathartica* | 0.0750 (−0.0464, 0.271) | 0.0270 (−0.0541, 0.128) | 0.0138 (−0.0739, 0.120) |
| *Frangula alnus* | | 0.00252 (−0.0766, 0.0788) | −0.00528 (−0.0731, 0.0539) |

| | | |
|---|---|---|
| *Lonicera morrowii* | −0.000906<br>(−0.0733, 0.0693) | 0.0105<br>(−0.0602, 0.0903) |

**Table S4.** Effects of species type and environmental context on the responses in **(A)** start of growth (time of 5% logarithmic growth; day of year), **(B)** annual total growth (mm; effect sizes on a log scale shown as equivalent percentage changes), **(C)** duration of growth (duration from 5% to 95% logarithmic growth; day), and **(D)** maximum relative growth rate ($day^{-1}$) per 3 °C warming. Mean ± standard deviations (*p*-values, *t*-test) are reported. Coefficients that are statistically significant ($p \leq 0.05$) are bolded.

| (A) start of growth | |
|---|---|
| term | estimate |
| intercept | **−3.41 ± 1.40 (0.0202)** |
| deciduous angiosperm tree | −1.89 ± 1.81 (0.3029) |
| exotic shrub | −2.69 ± 1.98 (0.1838) |
| closed canopy | −0.348 ± 1.76 (0.8440) |
| reduced rainfall | 0.200 ± 1.98 (0.9203) |
| deciduous angiosperm tree:closed canopy | 1.40 ± 2.30 (0.5456) |
| exotic shrub:closed canopy | 0.0401 ± 3.59 (0.9912) |
| deciduous angiosperm tree:reduced rainfall | −0.208 ± 2.56 (0.9358) |
| exotic shrub:reduced rainfall | −0.599 ± 2.80 (0.8320) |

| (B) annual total growth | |
|---|---|
| term | estimate |
| intercept | −0.216 ± 0.111 (0.0609) |
| deciduous angiosperm tree | **0.472 ± 0.144 (0.0023)** |
| exotic shrub | **0.380 ± 0.158 (0.0213)** |
| closed canopy | 0.144 ± 0.140 (0.3079) |
| reduced rainfall | −0.0354 ± 0.158 (0.8237) |
| deciduous angiosperm tree:closed canopy | −0.133 ± 0.182 (0.4696) |
| exotic shrub:closed canopy | 0.438 ± 0.286 (0.1338) |
| deciduous angiosperm tree:reduced rainfall | 0.0350 ± 0.203 (0.8643) |
| exotic shrub:reduced rainfall | 0.224 ± 0.223 (0.3212) |

| (C) duration of growth | |
|---|---|
| term | estimate |
| intercept | −3.10 ± 1.88 (0.1074) |
| deciduous angiosperm tree | 4.77 ± 2.42 (0.0565) |
| exotic shrub | 3.76 ± 2.65 (0.1646) |
| closed canopy | 0.422 ± 2.35 (0.8587) |
| reduced rainfall | −0.215 ± 2.65 (0.9360) |
| deciduous angiosperm tree:closed canopy | −2.05 ± 3.07 (0.5084) |
| exotic shrub:closed canopy | −2.58 ± 4.81 (0.5946) |
| deciduous angiosperm tree:reduced rainfall | 1.05 ± 3.43 (0.7609) |
| exotic shrub:reduced rainfall | 1.31 ± 3.75 (0.7286) |

| (D) relative growth rate | |
|---|---|
| term | estimate |
| intercept | 0.00125 ± 0.00665 (0.8515) |
| deciduous angiosperm tree | −0.000887 ± 0.00859 (0.9183) |
| exotic shrub | 0.00504 ± 0.00941 (0.5951) |
| closed canopy | 0.00285 ± 0.00834 (0.7344) |
| reduced rainfall | −0.00262 ± 0.00941 (0.7823) |
| deciduous angiosperm tree:closed canopy | 0.00337 ± 0.0109 (0.7590) |
| exotic shrub:closed canopy | **0.0658 ± 0.0171 (0.0005)** |
| deciduous angiosperm tree:reduced rainfall | 0.00656 ± 0.0121 (0.5925) |
| exotic shrub:reduced rainfall | 0.00467 ± 0.0133 (0.7275) |

**Table S5.** Summary statistics from linear models for responses in **(A)** start of growth (time of 5% logarithmic growth; day of year), **(B)** annual total growth (mm; effect sizes on a log scale shown as equivalent percentage changes), **(C)** duration of growth (duration from 5% to 95% logarithmic growth; day), and **(D)** maximum relative growth rate (day$^{-1}$) per 3 °C warming. From each model, we estimated an intercept, the effect of species' lifestyle (distribution or trait), the effect of environmental context, and the interaction between species' lifestyle and context. Effects of categorical lifestyle variables are modeled as differences between two categories. Effects of continuous lifestyle variables are modeled as change in responses every one standard deviation increase in the continuous variable. Effects of context are modeled as changes in responses moving from the wettest context (closed canopy, ambient rainfall), to the intermediate context (open canopy, ambient rainfall), and to the driest context (open canopy, reduced rainfall). Mean ± standard deviations (*p*-values, *t*-test) are reported. Coefficients that are statistically significant ($p \leq 0.05$) are bolded.

(A) start of growth

| lifestyle predictor | intercept | lifestyle effect | context effect | interaction |
|---|---|---|---|---|
| biome | **−4.28 ± 0.696 (< 0.0001)** | −0.687 ± 0.867 (0.4324) | −1.08 ± 0.839 (0.2044) | 0.979 ± 1.04 (0.3541) |
| northern range limit | **−4.75 ± 0.405 (< 0.0001)** | 0.526 ± 0.412 (0.2088) | −0.414 ± 0.488 (0.4013) | −0.648 ± 0.491 (0.1944) |
| southern range limit | **−4.81 ± 0.411 (< 0.0001)** | 0.505 ± 0.416 (0.2311) | −0.318 ± 0.496 (0.5249) | −0.672 ± 0.503 (0.1888) |
| range center | **−4.79 ± 0.403 (< 0.0001)** | 0.594 ± 0.407 (0.1520) | −0.337 ± 0.486 (0.4922) | −0.777 ± 0.495 (0.1239) |
| temperature niche | **−4.73 ± 0.414 (< 0.0001)** | −0.142 ± 0.426 (0.7408) | −0.446 ± 0.499 (0.3767) | 0.606 ± 0.501 (0.2331) |
| precipitation niche | **−4.71 ± 0.417 (< 0.0001)** | 0.413 ± 0.420 (0.3313) | −0.428 ± 0.502 (0.3985) | 0.367 ± 0.512 (0.4779) |
| origin | **−6.25 ± 1.01 (< 0.0001)** | 1.92 ± 1.12 (0.0932) | −0.164 ± 1.36 (0.9050) | −0.0737 ± 1.47 (0.9601) |
| leaf habit | **−5.29 ± 0.489 (< 0.0001)** | **1.83 ± 0.862 (0.0403)** | −0.660 ± 0.598 (0.2767) | 0.939 ± 1.02 (0.3639) |
| shoot growth type | **−4.30 ± 0.519 (< 0.0001)** | −1.18 ± 0.849 (0.1726) | −0.0998 ± 0.630 (0.8748) | −0.992 ± 1.02 (0.3356) |
| shade tolerance index | **−4.71 ± 0.403 (< 0.0001)** | **0.877 ± 0.412 (0.0393)** | −0.361 ± 0.486 (0.4609) | 0.127 ± 0.488 (0.7964) |

| | | | | |
|---|---|---|---|---|
| drought tolerance index | **−4.73 ± 0.424 (< 0.0001)** | −0.100 ± 0.430 (0.8172) | −0.440 ± 0.510 (0.3938) | 0.0740 ± 0.514 (0.8863) |
| specific leaf area | **−4.71 ± 0.411 (< 0.0001)** | −0.642 ± 0.432 (0.1445) | −0.432 ± 0.495 (0.3881) | −0.501 ± 0.499 (0.3213) |
| leaf C | **−4.51 ± 0.437 (< 0.0001)** | 0.812 ± 0.444 (0.0776) | −0.380 ± 0.535 (0.4833) | 0.291 ± 0.544 (0.5969) |
| leaf N | **−4.51 ± 0.405 (< 0.0001)** | **−1.23 ± 0.412 (0.0058)** | −0.380 ± 0.496 (0.4499) | 0.144 ± 0.504 (0.7771) |
| leaf C/N | **−4.51 ± 0.388 (< 0.0001)** | **1.38 ± 0.394 (0.0015)** | −0.380 ± 0.475 (0.4296) | −0.255 ± 0.482 (0.6005) |
| net photosynthetic rate | **−4.72 ± 0.407 (< 0.0001)** | −0.764 ± 0.416 (0.0735) | −0.424 ± 0.490 (0.3920) | −0.175 ± 0.493 (0.7239) |
| stomatal conductance | **−4.70 ± 0.381 (< 0.0001)** | **−1.22 ± 0.385 (0.0029)** | −0.301 ± 0.459 (0.5155) | −0.221 ± 0.465 (0.6373) |
| Vcmax | **−4.73 ± 0.420 (< 0.0001)** | −0.361 ± 0.427 (0.4035) | −0.443 ± 0.505 (0.3855) | −0.0268 ± 0.508 (0.9582) |

(B) annual total growth

| lifestyle predictor | intercept | lifestyle effect | context effect | interaction |
|---|---|---|---|---|
| biome | −0.0437 ± 0.0712 (0.5428) | **0.271 ± 0.0888 (0.0039)** | −0.0435 ± 0.0859 (0.6154) | 0.0524 ± 0.107 (0.6272) |
| northern range limit | **0.129 ± 0.0425 (0.0042)** | **−0.129 ± 0.0432 (0.0047)** | −0.0160 ± 0.0512 (0.7558) | −0.0591 ± 0.0515 (0.2576) |
| southern range limit | **0.122 ± 0.0422 (0.0063)** | **−0.137 ± 0.0427 (0.0025)** | −0.0334 ± 0.0510 (0.5152) | −0.0600 ± 0.0516 (0.2520) |
| range center | **0.125 ± 0.0415 (0.0046)** | **−0.142 ± 0.0419 (0.0015)** | −0.0281 ± 0.0501 (0.5777) | −0.0561 ± 0.0509 (0.2775) |
| temperature niche | **0.131 ± 0.0408 (0.0026)** | **0.154 ± 0.0420 (0.0007)** | −0.0117 ± 0.0492 (0.8127) | 0.0206 ± 0.0494 (0.6791) |
| precipitation niche | **0.132 ± 0.0469 (0.0076)** | 0.0299 ± 0.0473 (0.5307) | −0.00857 ± 0.0566 (0.8802) | 0.0124 ± 0.0577 (0.8308) |
| origin | **0.335 ± 0.111 (0.0044)** | **−0.255 ± 0.122 (0.0438)** | −0.0684 ± 0.149 (0.6495) | 0.0341 ± 0.161 (0.8333) |

| | | | | |
|---|---|---|---|---|
| leaf habit | **0.276 ± 0.0413 (< 0.0001)** | **−0.453 ± 0.0728 (< 0.0001)** | −0.0103 ± 0.0505 (0.8389) | −0.0838 ± 0.0864 (0.3380) |
| shoot growth type | 0.0685 ± 0.0569 (0.2355) | 0.162 ± 0.0931 (0.0892) | 0.0287 ± 0.0691 (0.6795) | −0.0895 ± 0.112 (0.4271) |
| shade tolerance index | **0.133 ± 0.0469 (0.0069)** | −0.0196 ± 0.0480 (0.6845) | −0.0133 ± 0.0565 (0.8150) | 0.0352 ± 0.0568 (0.5389) |
| drought tolerance index | **0.130 ± 0.0468 (0.0083)** | 0.0358 ± 0.0476 (0.4563) | −0.0126 ± 0.0564 (0.8250) | 0.0212 ± 0.0569 (0.7115) |
| specific leaf area | **0.130 ± 0.0373 (0.0012)** | **0.193 ± 0.0392 (< 0.0001)** | −0.0165 ± 0.0449 (0.7152) | 0.0418 ± 0.0453 (0.3606) |
| leaf C | **0.113 ± 0.0428 (0.0131)** | **−0.192 ± 0.0435 (0.0001)** | −0.0366 ± 0.0524 (0.4903) | 0.0412 ± 0.0532 (0.4452) |
| leaf N | **0.113 ± 0.0537 (0.0441)** | 0.0786 ± 0.0546 (0.1603) | −0.0366 ± 0.0658 (0.5822) | −0.0133 ± 0.0668 (0.8438) |
| leaf C/N | **0.113 ± 0.0519 (0.0374)** | **−0.109 ± 0.0527 (0.0481)** | −0.0366 ± 0.0635 (0.5688) | 0.0239 ± 0.0645 (0.7138) |
| net photosynthetic rate | **0.132 ± 0.0469 (0.0075)** | −0.0160 ± 0.0479 (0.7397) | −0.0101 ± 0.0565 (0.8590) | −0.0367 ± 0.0567 (0.5209) |
| stomatal conductance | **0.132 ± 0.0462 (0.0067)** | 0.0653 ± 0.0467 (0.1699) | −0.0191 ± 0.0557 (0.7331) | −0.0158 ± 0.0564 (0.7806) |
| Vcmax | **0.132 ± 0.0453 (0.0059)** | −0.0766 ± 0.0462 (0.1046) | −0.00936 ± 0.0546 (0.8647) | −0.0543 ± 0.0549 (0.3286) |

(C) duration of growth

| lifestyle predictor | intercept | lifestyle effect | context effect | interaction |
|---|---|---|---|---|
| biome | −0.466 ± 1.02 (0.6496) | 0.449 ± 1.27 (0.7252) | 1.41 ± 1.23 (0.2577) | −0.774 ± 1.53 (0.6154) |
| northern range limit | −0.160 ± 0.595 (0.7897) | −0.698 ± 0.606 (0.2556) | 0.867 ± 0.717 (0.2340) | 0.537 ± 0.722 (0.4615) |
| southern range limit | −0.107 ± 0.571 (0.8529) | **−1.38 ± 0.577 (0.0216)** | 0.611 ± 0.689 (0.3803) | 0.680 ± 0.699 (0.3359) |
| range center | −0.117 ± 0.577 (0.8407) | **−1.20 ± 0.582 (0.0464)** | 0.714 ± 0.696 (0.3107) | 0.751 ± 0.708 (0.2947) |

| | | | | |
|---|---|---|---|---|
| temperature niche | −0.171 ± 0.574 (0.7668) | 1.04 ± 0.590 (0.0844) | 0.897 ± 0.691 (0.2017) | −0.779 ± 0.694 (0.2680) |
| precipitation niche | −0.214 ± 0.601 (0.7237) | −0.0677 ± 0.606 (0.9116) | 0.920 ± 0.724 (0.2112) | −0.875 ± 0.738 (0.2427) |
| origin | 0.430 ± 1.51 (0.7773) | −0.840 ± 1.66 (0.6154) | 1.45 ± 2.03 (0.4776) | −0.772 ± 2.18 (0.7254) |
| leaf habit | 1.13 ± 0.646 (0.0890) | **−4.15 ± 1.14 (0.0008)** | 1.16 ± 0.791 (0.1499) | −1.49 ± 1.35 (0.2780) |
| shoot growth type | −0.859 ± 0.745 (0.2558) | 1.89 ± 1.22 (0.1299) | 0.448 ± 0.905 (0.6232) | 1.33 ± 1.46 (0.3671) |
| shade tolerance index | −0.187 ± 0.602 (0.7573) | −0.738 ± 0.616 (0.2377) | 0.835 ± 0.726 (0.2563) | −0.0969 ± 0.729 (0.8948) |
| drought tolerance index | −0.180 ± 0.604 (0.7666) | −0.592 ± 0.613 (0.3395) | 0.956 ± 0.728 (0.1960) | 0.157 ± 0.733 (0.8312) |
| specific leaf area | −0.200 ± 0.562 (0.7232) | **1.55 ± 0.591 (0.0120)** | 0.865 ± 0.677 (0.2088) | 0.949 ± 0.683 (0.1720) |
| leaf C | −0.0432 ± 0.656 (0.9479) | −0.396 ± 0.666 (0.5570) | 0.199 ± 0.803 (0.8064) | −0.212 ± 0.816 (0.7969) |
| leaf N | −0.0432 ± 0.571 (0.9402) | **1.81 ± 0.580 (0.0041)** | 0.199 ± 0.699 (0.7785) | −0.182 ± 0.710 (0.7997) |
| leaf C/N | −0.0432 ± 0.561 (0.9391) | **−1.89 ± 0.570 (0.0025)** | 0.199 ± 0.687 (0.7746) | 0.338 ± 0.698 (0.6320) |
| net photosynthetic rate | −0.184 ± 0.602 (0.7609) | 0.652 ± 0.615 (0.2950) | 0.889 ± 0.725 (0.2270) | 0.319 ± 0.728 (0.6632) |
| stomatal conductance | −0.224 ± 0.584 (0.7029) | 1.19 ± 0.590 (0.0513) | 0.774 ± 0.704 (0.2779) | 0.458 ± 0.712 (0.5236) |
| Vcmax | −0.175 ± 0.610 (0.7752) | 0.141 ± 0.621 (0.8222) | 0.907 ± 0.735 (0.2243) | −0.0348 ± 0.739 (0.9627) |

(D) relative growth rate

| lifestyle predictor | intercept | lifestyle effect | context effect | interaction |
|---|---|---|---|---|

| | | | | |
|---|---|---|---|---|
| biome | −0.000985 ± 0.00399 (0.8060) | 0.00982 ± 0.00497 (0.0548) | −0.00529 ± 0.00481 (0.2780) | 0.00357 ± 0.00599 (0.5545) |
| northern range limit | **0.00522 ± 0.00242 (0.0374)** | −0.00304 ± 0.00247 (0.2245) | −0.00312 ± 0.00292 (0.2926) | −0.00331 ± 0.00294 (0.2670) |
| southern range limit | 0.00497 ± 0.00248 (0.0516) | −0.00227 ± 0.00251 (0.3702) | −0.00331 ± 0.00299 (0.2759) | −0.00263 ± 0.00303 (0.3910) |
| range center | **0.00503 ± 0.00245 (0.0464)** | −0.00268 ± 0.00247 (0.2836) | −0.00327 ± 0.00295 (0.2748) | −0.00305 ± 0.00300 (0.3157) |
| temperature niche | **0.00532 ± 0.00244 (0.0349)** | 0.00303 ± 0.00251 (0.2343) | −0.00303 ± 0.00294 (0.3086) | 0.00269 ± 0.00295 (0.3676) |
| precipitation niche | **0.00551 ± 0.00244 (0.0295)** | 0.000913 ± 0.00246 (0.7130) | −0.00302 ± 0.00295 (0.3120) | 0.00392 ± 0.00300 (0.1988) |
| origin | **0.0220 ± 0.00545 (0.0002)** | **−0.0191 ± 0.00599 (0.0027)** | **−0.0215 ± 0.00731 (0.0053)** | **0.0196 ± 0.00787 (0.0170)** |
| leaf habit | **0.00736 ± 0.00299 (0.0182)** | −0.00603 ± 0.00528 (0.2608) | −0.00375 ± 0.00367 (0.3124) | 0.00101 ± 0.00627 (0.8732) |
| shoot growth type | 0.00458 ± 0.00310 (0.1472) | 0.00167 ± 0.00507 (0.7434) | −0.000307 ± 0.00376 (0.9353) | −0.00688 ± 0.00608 (0.2642) |
| shade tolerance index | **0.00551 ± 0.00248 (0.0319)** | 0.00131 ± 0.00254 (0.6094) | −0.00294 ± 0.00299 (0.3316) | 0.00237 ± 0.00300 (0.4343) |
| drought tolerance index | **0.00538 ± 0.00239 (0.0300)** | 0.00447 ± 0.00243 (0.0730) | −0.00336 ± 0.00288 (0.2496) | −0.00123 ± 0.00290 (0.6733) |
| specific leaf area | **0.00533 ± 0.00247 (0.0369)** | 0.00236 ± 0.00260 (0.3679) | −0.00308 ± 0.00298 (0.3059) | 0.000181 ± 0.00300 (0.9523) |
| leaf C | **0.00581 ± 0.00238 (0.0209)** | **−0.0113 ± 0.00242 (< 0.0001)** | −0.00405 ± 0.00291 (0.1747) | **0.00702 ± 0.00296 (0.0245)** |
| leaf N | 0.00581 ± 0.00326 (0.0855) | −0.00321 ± 0.00331 (0.3407) | −0.00405 ± 0.00400 (0.3188) | 0.00117 ± 0.00406 (0.7748) |

| | | | | |
|---|---|---|---|---|
| leaf C/N | 0.00581 ± 0.00330 (0.0891) | 0.00186 ± 0.00335 (0.5832) | −0.00405 ± 0.00404 (0.3244) | −0.000476 ± 0.00411 (0.9086) |
| net photosynthetic rate | **0.00539 ± 0.00244 (0.0328)** | −0.00318 ± 0.00249 (0.2098) | −0.00292 ± 0.00294 (0.3272) | −0.00206 ± 0.00295 (0.4905) |
| stomatal conductance | **0.00559 ± 0.00248 (0.0295)** | −0.000504 ± 0.00251 (0.8414) | −0.00304 ± 0.00299 (0.3145) | −0.00297 ± 0.00302 (0.3325) |
| Vcmax | **0.00536 ± 0.00241 (0.0318)** | −0.00415 ± 0.00245 (0.0982) | −0.00295 ± 0.00290 (0.3157) | −0.00153 ± 0.00292 (0.6024) |

**Table S6.** Baseline of the **(A)** start of growth (time of 5% logarithmic growth; day of year), **(B)** annual total growth (mm), **(C)** duration of growth (duration from 5% to 95% logarithmic growth; day), and **(D)** maximum relative growth rate (day$^{-1}$) in ambient warming treatment under various environmental contexts. Medians and 95% credible intervals of effect sizes are reported. Order of species follows that in Fig. 2A.

(A) start of growth

| species | closed canopy, ambient rainfall | open canopy, ambient rainfall | open canopy, reduced rainfall |
|---|---|---|---|
| *Picea mariana* | 123<br>(110, 133) | | |
| *Picea glauca* | 128<br>(120, 135) | 126<br>(118, 133) | 127<br>(119, 134) |
| *Pinus banksiana* | 117<br>(106, 125) | 108<br>(96.4, 117) | 107<br>(94.9, 117) |
| *Abies balsamea* | 124<br>(113, 133) | 129<br>(118, 138) | 129<br>(118, 138) |
| *Pinus resinosa* | 112<br>(102, 119) | | |
| *Pinus strobus* | 122<br>(113, 130) | 116<br>(105, 124) | 120<br>(111, 128) |
| *Tsuga canadensis* | 124<br>(113, 134) | | |
| *Populus tremuloides* | 133<br>(126, 139) | 135<br>(125, 142) | 133<br>(122, 140) |
| *Betula papyrifera* | 138<br>(129, 145) | 125<br>(115, 133) | 130<br>(119, 139) |
| *Betula alleghaniensis* | 124<br>(113, 133) | | |
| *Quercus macrocarpa* | 127<br>(121, 133) | 128<br>(122, 133) | 129<br>(123, 134) |
| *Acer negundo* | 107<br>(98.1, 115) | | |
| *Acer saccharum* | 130<br>(124, 136) | 126<br>(120, 132) | 129<br>(123, 133) |
| *Quercus rubra* | 126<br>(120, 131) | 134<br>(129, 138) | 132<br>(126, 136) |

| | | | |
|---|---|---|---|
| *Tilia americana* | 124<br>(118, 130) | | |
| *Acer rubrum* | 130<br>(123, 137) | 128<br>(120, 134) | 125<br>(116, 132) |
| *Lonicera tatarica* | | 113<br>(91.5, 126) | 111<br>(86.1, 125) |
| *Rhamnus cathartica* | 137<br>(128, 143) | 134<br>(121, 143) | 138<br>(128, 143) |
| *Frangula alnus* | | 135<br>(117, 145) | 132<br>(112, 144) |
| *Lonicera morrowii* | | 118<br>(101, 129) | 123<br>(107, 133) |

(B) annual total growth

| species | closed canopy, ambient rainfall | open canopy, ambient rainfall | open canopy, reduced rainfall |
|---|---|---|---|
| *Picea mariana* | 35.7<br>(23.1, 56.8) | | |
| *Picea glauca* | 33.8<br>(24.0, 47.1) | 119<br>(94.1, 153) | 93.6<br>(73.0, 120) |
| *Pinus banksiana* | 150<br>(117, 197) | 360<br>(275, 461) | 383<br>(307, 469) |
| *Abies balsamea* | 39.2<br>(28.5, 55.6) | 79.2<br>(57.9, 110) | 74.5<br>(54.7, 101) |
| *Pinus resinosa* | 95.2<br>(65.1, 136) | | |
| *Pinus strobus* | 140<br>(83.6, 227) | 165<br>(108, 258) | 175<br>(123, 240) |
| *Tsuga canadensis* | 68.4<br>(38.3, 121) | | |
| *Populus tremuloides* | 29.5<br>(21.1, 41.8) | 135<br>(87.4, 209) | 110<br>(64.4, 187) |
| *Betula papyrifera* | 76.3<br>(50.2, 121) | 206<br>(158, 269) | 147<br>(105, 207) |
| *Betula alleghaniensis* | 72.9<br>(46.0, 116) | | |

| | | | |
|---|---|---|---|
| *Quercus macrocarpa* | 41.0<br>(29.1, 59.7) | 85.6<br>(63.8, 118) | 78.1<br>(56.0, 110) |
| *Acer negundo* | 38.4<br>(31.2, 48.0) | | |
| *Acer saccharum* | 48.5<br>(36.3, 66.9) | 83.1<br>(56.6, 123) | 77.8<br>(55.4, 111) |
| *Quercus rubra* | 45.7<br>(31.9, 66.5) | 134<br>(103, 177) | 115<br>(81.9, 161) |
| *Tilia americana* | 51.5<br>(39.4, 67.9) | | |
| *Acer rubrum* | 38.8<br>(30.8, 50.4) | 121<br>(86.6, 168) | 77.9<br>(51.0, 120) |
| *Lonicera tatarica* | | 64.7<br>(42.0, 100) | 46.1<br>(29.1, 73.3) |
| *Rhamnus cathartica* | 47.9<br>(30.2, 80.5) | 44.0<br>(24.1, 77.6) | 57.3<br>(32.9, 98.4) |
| *Frangula alnus* | | 113<br>(72.8, 178) | 115<br>(63.2, 204) |
| *Lonicera morrowii* | | 42.6<br>(25.5, 71.2) | 36.8<br>(21.3, 62.1) |

(C) duration of growth

| species | closed canopy, ambient rainfall | open canopy, ambient rainfall | open canopy, reduced rainfall |
|---|---|---|---|
| *Picea mariana* | 69.3<br>(50.8, 93.2) | | |
| *Picea glauca* | 35.1<br>(24.4, 49.6) | 42.1<br>(30.7, 57.5) | 41.5<br>(30.6, 56.4) |
| *Pinus banksiana* | 44.8<br>(31.7, 63.9) | 56.8<br>(41.4, 78.1) | 60.8<br>(43.9, 84.3) |
| *Abies balsamea* | 49.4<br>(34.8, 69.8) | 50.3<br>(35.5, 70.8) | 53.4<br>(38.0, 74.6) |
| *Pinus resinosa* | 67.1<br>(52.4, 85.9) | | |
| *Pinus strobus* | 45.2<br>(32.9, 61.5) | 51.8<br>(37.7, 71.4) | 51.6<br>(38.1, 69.2) |

| | | | |
|---|---|---|---|
| *Tsuga canadensis* | 72.3<br>(54.3, 94.7) | | |
| *Populus tremuloides* | 23.3<br>(16.6, 37.0) | 29.3<br>(18.1, 48.5) | 33.8<br>(20.3, 55.1) |
| *Betula papyrifera* | 37.2<br>(25.5, 53.9) | 57.0<br>(41.8, 77.2) | 56.6<br>(40.8, 77.8) |
| *Betula alleghaniensis* | 74.6<br>(57.2, 96.4) | | |
| *Quercus macrocarpa* | 22.9<br>(17.1, 31.9) | 30.6<br>(22.4, 42.5) | 28.6<br>(20.8, 40.0) |
| *Acer negundo* | 55.3<br>(42.0, 72.0) | | |
| *Acer saccharum* | 21.2<br>(16.7, 27.6) | 30.1<br>(21.9, 40.9) | 26.3<br>(19.3, 35.7) |
| *Quercus rubra* | 22.6<br>(16.8, 32.6) | 21.2<br>(16.5, 29.9) | 22.2<br>(16.6, 32.0) |
| *Tilia americana* | 39.3<br>(30.2, 51.3) | | |
| *Acer rubrum* | 23.8<br>(17.1, 35.2) | 38.4<br>(27.6, 53.1) | 39.8<br>(27.5, 57.2) |
| *Lonicera tatarica* | | 48.7<br>(25.7, 89.7) | 51.4<br>(26.1, 102) |
| *Rhamnus cathartica* | 19.1<br>(16.1, 32.3) | 27.0<br>(16.9, 50.5) | 24.4<br>(16.5, 44.3) |
| *Frangula alnus* | | 38.3<br>(20.8, 73.8) | 47.0<br>(25.3, 86.8) |
| *Lonicera morrowii* | | 38.6<br>(21.2, 71.1) | 35.4<br>(19.0, 67.7) |

(D) relative growth rate

| species | closed canopy, ambient rainfall | open canopy, ambient rainfall | open canopy, reduced rainfall |
|---|---|---|---|
| *Picea mariana* | 0.0693<br>(0.0497, 0.0969) | | |
| *Picea glauca* | 0.114<br>(0.0794, 0.166) | 0.126<br>(0.0923, 0.175) | 0.120<br>(0.0883, 0.165) |

| | | | |
|---|---|---|---|
| *Pinus banksiana* | 0.110 (0.0779, 0.156) | 0.102 (0.0740, 0.140) | 0.0998 (0.0720, 0.138) |
| *Abies balsamea* | 0.0877 (0.0609, 0.125) | 0.0921 (0.0647, 0.131) | 0.0890 (0.0631, 0.125) |
| *Pinus resinosa* | 0.0544 (0.0410, 0.0727) | | |
| *Pinus strobus* | 0.109 (0.0763, 0.153) | 0.0907 (0.0638, 0.128) | 0.0955 (0.0694, 0.131) |
| *Tsuga canadensis* | 0.0833 (0.0619, 0.115) | | |
| *Populus tremuloides* | 0.146 (0.0914, 0.211) | 0.167 (0.100, 0.271) | 0.142 (0.0851, 0.237) |
| *Betula papyrifera* | 0.110 (0.0734, 0.163) | 0.0931 (0.0682, 0.127) | 0.0876 (0.0629, 0.123) |
| *Betula alleghaniensis* | 0.0513 (0.0375, 0.0697) | | |
| *Quercus macrocarpa* | 0.191 (0.135, 0.261) | 0.163 (0.117, 0.224) | 0.168 (0.119, 0.232) |
| *Acer negundo* | 0.0529 (0.0397, 0.0708) | | |
| *Acer saccharum* | 0.144 (0.108, 0.189) | 0.130 (0.0926, 0.183) | 0.147 (0.105, 0.205) |
| *Quercus rubra* | 0.184 (0.126, 0.252) | 0.234 (0.166, 0.305) | 0.218 (0.150, 0.295) |
| *Tilia americana* | 0.0970 (0.0735, 0.128) | | |
| *Acer rubrum* | 0.135 (0.0915, 0.191) | 0.125 (0.0894, 0.174) | 0.103 (0.0700, 0.153) |
| *Lonicera tatarica* | | 0.102 (0.0553, 0.196) | 0.0847 (0.0431, 0.166) |
| *Rhamnus cathartica* | 0.210 (0.124, 0.275) | 0.137 (0.0716, 0.230) | 0.162 (0.0864, 0.253) |
| *Frangula alnus* | | 0.125 (0.0651, 0.234) | 0.105 (0.0551, 0.200) |
| *Lonicera morrowii* | | 0.119 (0.0634, 0.216) | 0.111 (0.0563, 0.210) |
